# Repeated Transitions in Migratory Behavior Across a Songbird Radiation Are Not Driven by Selection on Shared Variation in Candidate Genes

**DOI:** 10.64898/2026.10.30.763195

**Authors:** Stephanie J. Szarmach, Kevin F. P. Bennett, David P. L. Toews

## Abstract

Understanding the mechanisms facilitating the repeated evolution of traits is a core question in evolutionary biology. While repeated evolution has been observed across many systems, behavioral phenotypes have been less well studied than morphological traits. Migration is a behavioral adaptation to seasonality that has evolved independently in many animals, and repeated gains and losses of migratory behavior have been especially prevalent throughout the diversification of birds. While migratory traits are heritable in many taxa and candidate genes underlying intraspecific variation in migratory behavior have been identified, whether similar genetic changes are responsible for the repeated evolution of migration remains poorly understood. We used a phylogenomic approach to investigate the relationship between variation in candidate genes and migratory propensity across nearly all species in the parulid warbler radiation (*n* = 101), a group of songbirds in which migratory behavior has been lost and gained multiple times. Comparisons of candidate gene tree topologies to the parulid warbler species tree revealed cases of discordance, though these were explained by differences in the number of phylogenetically informative sites among genes rather than by elevated genetic similarity among species sharing migratory strategies. Testing models of coding sequence evolution identified sites with elevated dN/dS in five candidate genes, suggestive of positive selection; however, variation at these sites showed no clear association with migratory strategy, nor did we detect recurrent selection across lineages where transitions in migratory behavior most likely occurred. Together, these results contradict the hypothesis that selection on shared variation (ancestral polymorphisms or introgressed alleles) facilitated the repeated evolution of migration in this clade.

---

Understanding the degree to which evolution is predictable—and identifying what conditions facilitate the repeated evolution of a phenotype or genotype—has long been a central goal of evolutionary biology. Repeated evolution can occur at the phenotypic level when a similar trait arises multiple times in independent taxa, typically in response to similar environmental conditions or selective pressures (Bolnick et al. 2018; Cerca 2023). Similar phenotypes sometimes evolve independently through completely different genetic mechanisms, such as in cases of convergent pigmentation patterns in *Monarcha* flycatchers (Uy et al. 2016, 2019) and *Ranitomeya* poison frogs (Twomey et al. 2023). However, repeated evolution can also occur at the level of a pathway, gene, or genetic variant, with similar molecular mechanisms producing similar phenotypes in independent lineages. In some cases, repeated evolution at the genetic level has occurred among distantly related species, as in the case of mutations in *MC1R* leading to pigmentation changes in mammals, birds, and non-avian reptiles (Arendt and Reznick 2008).

However, repeated evolution at the molecular level is thought to be more prevalent in closely related species, due to shared evolutionary history and genetic or developmental constraints (Conte et al. 2012). Many notable examples of repeated evolution at both the phenotypic and genotypic levels have been observed in adaptive radiations, such as the repeated evolution of morphologically similar ecotypes in three-spined stickleback (*Gasterosteus aculeatus*; Jones et al. 2012) and Arctic charr (*Salvelinus alpinus*; Jacobs et al. 2020), and the parallel evolution of mimetic wing pattern in *Heliconius* butterflies (Kronforst et al. 2006). Associations between specific genetic variants and the independent evolution of phenotypic traits can be due to either repeated selection on standing variation shared between populations or species (Barrett et al. 2008; Jones et al. 2012), or due to hybridization and introgression resulting in the sharing of trait-associated variants (Dasmahapatra et al. 2012; Baiz et al. 2021; Bennett et al. 2025).

Most studies of repeated evolution have examined morphological traits, particularly coloration, while behavioral traits have received less focus (York and Fernald 2017). In cases where the repeated evolution of behaviors has been identified—such as mate guarding in isopod ecotypes (Karlsson et al. 2010), territorial behavior in *Anolis* lizards (Johnson et al. 2010), and activity patterns in *Heliconius* butterflies occupying different elevations (Rivas-Sánchez et al. 2023)—whether similar genetic mechanisms have facilitated the evolution of these parallel traits often remains unknown. One reason for this may be that while some behavioral traits are controlled by a few loci of large effect, like burrowing behavior in *Peromyscus* mice (Weber et al. 2013), many are likely highly polygenic (York 2018). However, advances in whole-genome sequencing now facilitate analyses of variation at many genes simultaneously across a large number of taxa, improving researchers’ ability to explore relationships between genetic variation and the repeated evolution of complex behavioral traits.

Seasonal migration is a behavior that has arisen independently across a broad range of animal taxa, including birds, mammals, fish, and insects. Throughout avian diversification, migratory behavior has been gained and lost many times (Outlaw et al. 2003; Winger et al. 2012, 2014; Gómez-Bahamón et al. 2020). Variation in migratory propensity (i.e., migratory vs. sedentary behavior) is present at both the interspecific and intraspecific level, with some species exhibiting “partial migration” (i.e., some populations or individuals do not migrate, while others do; Chapman et al. 2011). A large body of work has established migratory behavior as a heritable trait in birds (Berthold and Querner 1981; Berthold and Pulido 1994; Pulido 2007), and over the past two decades specific genes have been associated with variation in migratory propensity among individuals and populations (reviewed in Gu et al. 2024). Yet, while some especially well-studied genes, like *CLOCK* and *ADCYAP1* have been linked to intraspecific variation in migratory behavior across multiple taxa (e.g., Mueller et al. 2011; Bazzi et al. 2015; Saino et al. 2015), in other species these genes show no association with migration-related traits (e.g. Peterson et al. 2013; Contina et al. 2018; Sauve et al. 2021).

Genes linked with migratory variation at the individual and population level have rarely been examined across independent lineages in a comparative framework, limiting conclusions about their relationship to variation between species. One study compared 25 candidate migration genes (first proposed by Ruegg et al. 2014) across 70 bird species spanning all avian orders, but found that no genetic variants consistently differed between migratory and sedentary species and that signatures of positive selection were not associated with transitions in migratory phenotype (Lugo Ramos et al. 2017). In hindsight, this result may not be unexpected given the broad phylogenetic scale of the study, which compared birds with diverse migratory strategies that likely evolved through very different mechanisms. For example, nocturnal and diurnal migrants likely require different modes of navigation and distinctive circadian rhythms (Piersma et al. 2005; Zink 2011). Short-lived solitary migrants presumably rely more on an innate migratory program, while long-lived species that migrate in groups may depend more on learning and social interactions (Berthold 1990; Pulido 2007; Mueller et al. 2013).

Identifying instances where parallel genetic mechanisms facilitate the repeated evolution of migratory behavior may be more likely when comparing more closely related species, because the syndrome of adaptations that facilitate migration are more likely to be directly comparable. In addition, species that share a more recent common ancestor are more likely to share genetic variation—either ancestral polymorphisms or alleles introduced through introgression—that selection can act on repeatedly (Ord and Summers 2015; Waters and McCulloch 2021). In line with this, Lugo Ramos et al. (2017) found that variation in certain candidate genes was more closely associated with differences in migratory phenotype when restricting the analysis to a single order (Passeriformes).

Here, we use whole-genome resequencing data from nearly all species in Parulidae to test whether candidate genes associated with migratory phenotype at the intraspecific level are also related to interspecific variation in migratory propensity. The parulid warblers, a radiation of small songbirds that diversified over the last 7 million years (Oliveros et al. 2019), are a valuable system for studying the evolution of migration because they exhibit a diversity of migratory phenotypes, ranging from full residency to long-distance migration (Winger et al. 2012).

Previous analyses using the Lovette et al. (2010) phylogeny of Parulidae found that the family’s common ancestor was most likely migratory, and that migration has been subsequently lost and regained multiple times across the radiation (Winger et al. 2012; Mitchell et al. 2019), making this group well positioned for studying the repeated evolution of migratory behavior. Because hybridization occurs between many parulid warblers and previous work has detected introgression of carotenoid processing genes associated with plumage pigmentation (Baiz et al. 2021; Bennett et al. 2025), this system also offers an opportunity to test whether past gene flow has facilitated the transfer of adaptive variants that facilitate migration. Studies have identified genes associated with migratory variation within parulid species (Bazzi et al. 2016; Ralston et al. 2019; Toews et al. 2019; Zamudio-Beltrán et al. 2024), but to date no work has systematically studied interspecific variation in candidate migration genes across the radiation.

Using low-coverage whole genome resequencing data from 101 parulid warbler taxa, we produced alignments for 24 of the same candidate migration genes tested by Lugo Ramos et al. (2017) plus one additional gene, *VPS13A*, which varies with migratory phenotype and exhibits signatures of selection in *Vermivora* warblers (Toews et al. 2019). We constructed gene trees and compared their topologies to a revised phylogeny of Parulidae inferred from ultra-conserved elements (UCEs; Bennett et al. 2025). If repeated evolution of migratory or sedentary behavior in this clade has been facilitated by selection on shared variation—whether retained ancestral polymorphisms or introgressed alleles—gene trees should exhibit clustering of species according to their migratory strategy more so than phylogenetic relationships, producing discordance with the species tree. We then investigated how rates of coding sequence evolution relate to transitions in migratory phenotype by estimating nonsynonymous to synonymous substitution rate ratios (dN/dS) in PAML (Yang 1997, 2007). If similar genetic mechanisms are responsible for the repeated evolution of migratory behavior across the Parulidae, we expect to detect signatures of positive selection in the same genes across divergent migratory clades.

## Materials and Methods

### Whole Genome Sequencing and Species Tree Construction

We generated a whole-genome resequencing dataset for 480 individuals from 101 parulid warbler taxa and one outgroup, the yellow-breasted chat (*Icteria virens*; Supplementary Table S1). This dataset includes nearly all species in the family Parulidae (116 species according to the Clements taxonomy; Clements et al. 2025), and represents 17 of 18 recognized genera. We sampled 1–12 individuals per taxon (mean = 4.7), and 89% of taxa had samples from at least four individuals. DNA was extracted from blood samples collected in the field from breeding birds (*n* = 108) or tissue samples obtained from museum loans (*n* = 373) using Qiagen DNeasy Blood & Tissue kits (sample information can be found in Supporting Information of Bennett et al. 2025). Blood samples were obtained under IACUC protocols approved by Cornell University (#2015-0065) or Penn State University (#201900879) and under a USGS federal master bird banding permit (#24043 to D.P.L.T.), New York State Department of Environmental Conservation banding permit (#156), Pennsylvania Game Commission banding permit (#46121), and Pennsylvania State Forest SFRA permit #2220.

Dual-indexed libraries with P2 150bp paired-end chemistry were sequenced on Illumina HighSeq and NextSeq systems. Sequencing protocols are described for *Setophaga*, *Catharopeza*, *Vermivora cyanoptera*, and *V. chrysoptera* in Baiz et al. (2021), for *V. bachmanii* in Wood et al. (2023), and for all other species in Bennett et al. (2025). We trimmed sequencing reads with AdapterRemoval (Schubert et al. 2016), marked PCR duplicates with PicardTools (Broad Institute 2024), and aligned reads to the *Setophaga coronata* reference genome (Baiz et al. 2021; NCBI Accession #GCA_001746935.2) using Bowtie 2 (Langmead and Salzberg 2012). The average coverage per sample was 3.8X and all samples aligned to the reference at a high alignment rate (mean = 85.4% reads mapped) with no evidence of alignment biases (Baiz et al. 2021; Bennett et al. 2025).

Because we only required a single representative sequence from each species for phylogeny construction, we combined BAM alignment files from all individuals of each taxon into a single consensus sequence. This approach allows us to increase confidence in base calling by combining reads from many individuals sequenced at low coverage into a moderate-coverage (∼20X) species alignment, and has previously been used successfully for phylogenomic analyses of parulid warblers (Baiz et al. 2021; Bennett et al. 2025). To do this, we first combined BAM files from individual samples into a single file for each taxon using samtools *merge* (Li et al. 2009; Danecek et al. 2021) and then generated a VCF file containing genotype likelihoods for variant sites in the alignment using bcftools *mpileup* (Danecek et al. 2021). We called variants using bcftools *call* and used bcftools *filter* to remove variants with low site quality scores (QUAL < 20) and abnormally high read depth (DP > 200), which can arise from misalignment of repetitive regions. Variants were normalized using bcftools *norm*. The parulid warbler species tree (Fig. 1) was constructed using 4,016 concatenated ultra-conserved elements (UCE) and flanking regions, as described in Baiz et al. (2021) and Bennett et al. (2025). We inferred the species tree in IQ-TREE (Minh et al. 2020) using the TVM+F+I+R4 DNA evolution model and ran 1000 replicates of an approximate likelihood ratio test (aLRT) to estimate branch support.

**FIGURE 1.**
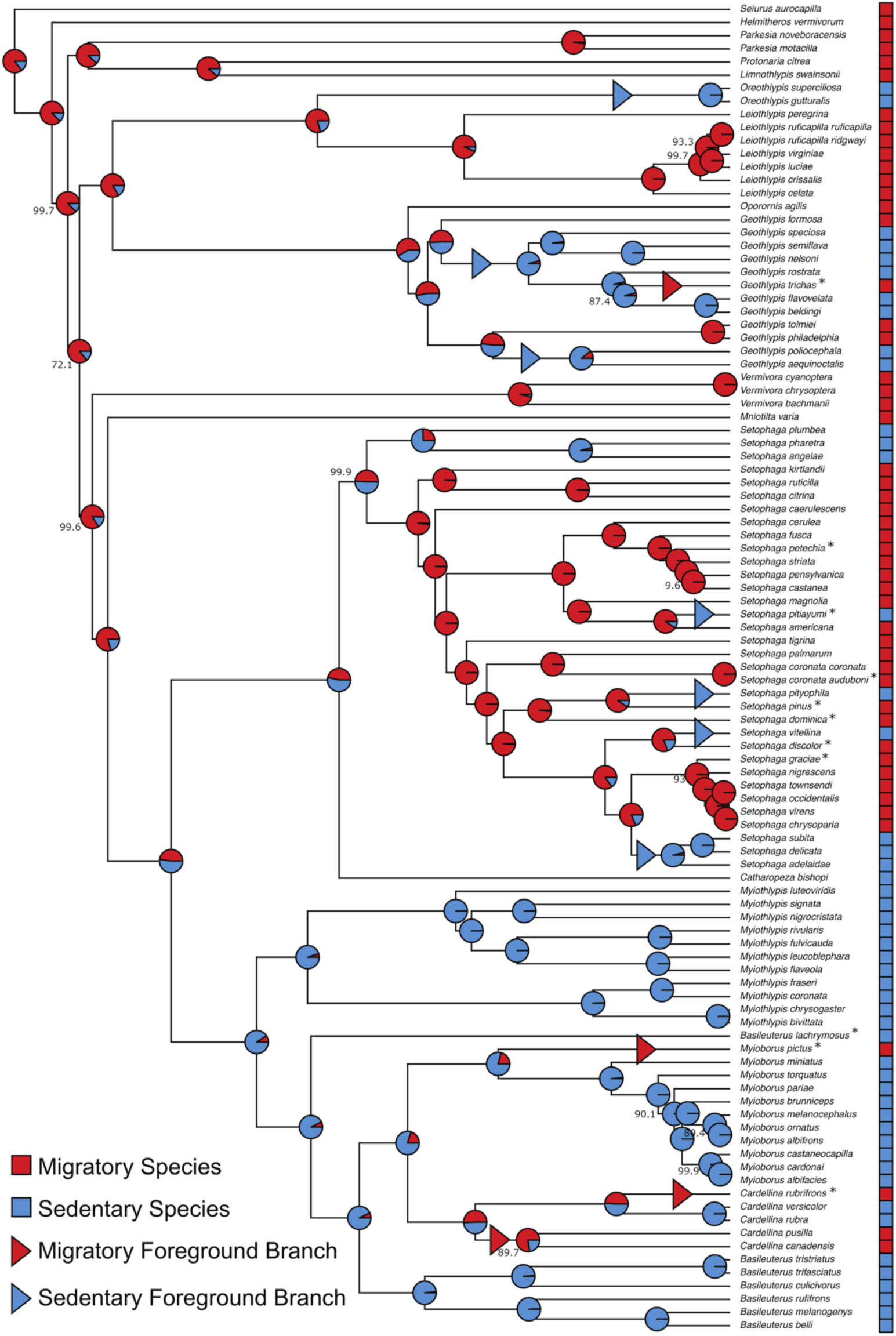
Parulid warbler species tree labeled with migratory propensity of each species (squares to the right of scientific names), results of ancestral character state reconstruction (pie charts on nodes), and foreground branches selected for PAML analyses of coding sequence evolution(triangles on branches). Nodes of the species tree that received <100% support by approximate likelihood ratio test (aLRT) are labeled with their aLRT value. Migratory propensity is indicated by color (red = migratory, blue = sedentary). Pie charts represent the posterior probabilities of the two character states at each node. Asterisks next to species names indicate that a species is partially migratory: a binary character state was assigned for these species based on the migratory propensity of the population from which the samples were collected.

### Coding Migratory Propensity

We classified the migratory propensity of each species based on Birds of the World species accounts (Fig. 1; Supplementary Table S1; Billerman et al. 2022). Species were coded as migratory if all populations and individuals migrate (often described in species accounts as “complete” or “obligate” migrants; *n* = 42) and as sedentary if all or nearly all individuals are permanent residents that do not migrate (*n* = 48). Eleven species were “partially migratory,” with some populations migrating while others remain resident throughout the year. For downstream analyses using discrete binary character states, we coded partially migratory species as migratory or sedentary based on the collection location of the samples. For five partially migratory species, all samples were obtained from migratory regions, and for two species all samples were from sedentary regions. Samples for four partially migratory species were collected from both migratory and resident regions. We categorized these species as migratory, because for each a greater proportion of samples either originated from the migratory portion of the range or was identified as a migratory subspecies.

### Alignments of Candidate Migration Genes

We extracted sequences for 25 genes previously identified as candidates potentially influencing variation in migratory phenotype (Supplementary Table S2). Twenty-four of these candidate genes were evaluated for association with migratory variation among 70 highly divergent species from all avian orders by Lugo Ramos et al. (2017). These were included in the present study to directly compare whether genes that were not found to vary in relation to migratory propensity at a broad taxonomic scale do associate with migratory phenotype at a finer intrafamilial scale. One gene included in the study by Lugo Ramos et al. (2017), *ADCYAP1*, was excluded from our analysis because much of the gene overlapped gaps in the myrtle warbler reference genome, yielding an unreliable alignment. We included one additional gene not examined by Lugo Ramos et al., *VPS13A*, which has been associated with differences in migratory direction and distance and exhibits signatures of selection in *Vermivora* warblers (Toews et al. 2019).

The known functions of these candidate genes (typically characterized in model organisms) encompass control of circadian rhythms, cellular stress response, metabolism, and cell signaling (Supplementary Table S2). Some candidate genes have been associated with exploratory behavior (Garamszegi et al. 2014; Timm et al. 2019), and variants in others lead to movement disorders in model organisms, suggesting a functional role in controlling locomotion (Ishida et al. 2009; Shi et al. 2011). Most of the included candidate genes have been linked to migratory phenotype in birds through studies identifying genetic differentiation associated with variation in migratory behavior or measuring differential gene expression in birds in migratory vs. nonmigratory states (Supplementary Table S2). Five genes were proposed as candidates originally by Ruegg et al. (2014) based on known function, but associations with migratory phenotype have not been shown empirically. These genes were retained in the study for comparability with Lugo Ramos et al. (2017).

We extracted candidate gene sequences (including untranslated regions [UTRs], introns, and coding regions) and generated consensus sequences for each species using the same protocol described for extracting UCE regions. The genomic location of each candidate gene was determined by annotating the *S. coronata* reference genome based on the zebra finch assembly (*Taeniopygia guttata*; NCBI Accession #GCF_003957565.2) using LiftOff with default options (Shumate and Salzberg 2021). In cases where multiple alleles were present at a position in the consensus sequence for a species, IUPAC ambiguity codes were used to represent the bases that were present. Regions containing long stretches of N’s due to gaps in the reference genome assembly were manually removed from the alignment using Geneious Prime (v2025.0.3, https://www.geneious.com).

For analyses of coding sequence evolution in PAML, alignments containing only coding regions (CDS) were constructed in Geneious by extracting CDS regions from the full gene alignment and then concatenating them. CDS regions were identified from the *S. coronata* genome annotation and checked for completeness against the annotated zebra finch genome (bTaeGut1_v1.p) using BLAT in the Ensembl genome browser (Harrison et al. 2024). The CDS alignments of six genes were missing one codon and two genes were missing two codons due to gaps in the *S. coronata* reference genome. Each extracted CDS region was trimmed manually in Geneious to align with the correct reading frame before concatenation. Protein sequence alignments were generated by translating the CDS region alignments in Geneious.

We found that when ambiguity codes were included in CDS alignments, we were not able to run PAML analyses because the number of ambiguous codons became computationally intractable, particularly for the longest or most variable genes. To ameliorate this issue, we used a custom Python script (Supplementary Data) to replace ambiguous nucleotides with the most common allele at that site in the alignment. Although information about within-species polymorphisms is lost using this approach, this analysis was intended to detect fixed differences between species resulting from positive selection.

### Gene Tree Construction and Gene Tree versus Species Tree Comparisons

We inferred maximum likelihood phylogenies for each candidate gene in IQ-TREE using the concatenated full gene alignments. The best supported model of sequence evolution was identified for each gene using ModelFinder (Kalyaanamoorthy et al. 2017; Supplementary Table S3). To estimate branch support, we ran 1000 bootstrap replicates implementing the ultrafast bootstrap approximation (UFBoot; Minh et al. 2013; Hoang et al. 2018).

To visualize discordance between each gene tree and the parulid species tree, we plotted cophylogenies using the “cophylo” function in phytools (Revell 2024). To quantify discordance between the trees, we calculated Jaccard Robinson-Foulds (JRF) distances (Böcker et al. 2013; Smith 2020) using the R package TreeDist (Smith 2019, 2022). The JRF distance is an extension of the classical Robinson-Foulds metric for comparing phylogenetic trees (Robinson and Foulds 1981) that allows for greater resolution by taking into account splits that are similar, but not identical, rather than simply counting splits unique to one of the two trees (Böcker et al. 2013; Smith 2020). We report both the raw JRF distance and the normalized JRF distance, which is rescaled based on the maximum possible value for the set of trees and ranges from 0 (trees identical) to 1 (no splits shared between trees).

We quantified and compared phylogenetic signal in migratory propensity between the parulid warbler species tree and candidate migration gene trees using the *D* statistic of Fritz and Purvis (2010) implemented in the R package caper (Orme et al. 2023). The *D* statistic is a measure of phylogenetic signal in a binary trait derived from the sum of sister-clade differences in a tree (Fritz and Purvis 2010). When the distribution of traits across the tree follows the expectations of a Brownian motion model of evolution, *D* = 0, and when traits are distributed in a phylogenetically random manner, *D* = 1. A *D* statistic value < 0 indicates that the trait has higher phylogenetic signal than expected under a Brownian motion model (i.e., higher “clustering” of the trait), and PhyloD > 1 means the trait is phylogenetically overdispersed. For each gene tree, a randomization test with 10,000 permutations was performed to assess whether the observed phylogenetic tree significantly deviated from either the random topology model or the Brownian motion model. If genetic variants within candidate migration genes are shared between distantly related migratory species, we expect migratory propensity to exhibit greater phylogenetic signal (i.e., lower values of *D*) in the gene trees compared to the parulid species tree.

### Reconstructing Ancestral Character States

To inform the selection of focal branches for analyses of coding sequence evolution in PAML, we performed an ancestral character state reconstruction (ACSR) to identify the most likely migratory state of each node within the parulid warbler species tree and to infer when transitions between migratory and sedentary behavior likely occurred. The ACSR was implemented using the R packages phytools (Revell 2024) and corHMM (Beaulieu et al. 2013). We first rooted the species tree using the outgroup, *Icteria virens*, and then removed the outgroup using the “drop.tip” function in the R package ape (Paradis and Schliep 2019). We converted the phylogram to an ultrametric tree using the “chronos” function in ape, applying the “clock” model of substitution rate variation and lambda = 1. We used the “fitMk” function in phytools to fit four models of discrete character evolution (equal rates [ER], all rates different [ARD], only transitions from migratory to sedentary [M→S], and only transitions from sedentary to migratory [S→M]). We determined that the ER model was the best fitting using the Akaike information criterion (AIC; Supplementary Table S4). We performed a marginal ancestral character state reconstruction using corHMM specifying the ER model, with one rate category and no fixed nodes. We plotted the phylogeny with pie charts representing the probability of each character state at each node using the “plotTree” function in phytools and the “nodelabels” function in ape.

### Modeling Rates of Coding Sequence Evolution

To investigate whether rates of coding sequence evolution in the 25 candidate migration genes were associated with transitions in migratory phenotype, we tested models of molecular evolution using the CODEML program in PAML (Yang 1997, 2007). CODEML estimates dN/dS (denoted as ω), the ratio of the nonsynonymous substitution rate to the synonymous substitution rate, using maximum likelihood codon-based models to detect signatures of selection in protein coding genes (Álvarez-Carretero et al. 2023). A nonsynonymous substitution rate (dN) greater than the synonymous substitution rate (dS; ω > 1) is indicative of positive selection increasing the fixation of adaptive nonsynonymous mutations. A dN/dS ratio less than one (ω < 1) suggests the gene has experienced purifying selection, preventing the fixation of deleterious nonsynonymous variants in the population. When a sequence is evolving neutrally, ω ≈ 1. Although numerous approaches exist for inferring rates of coding sequence evolution, we tested for signatures of positive selection using dN/dS to enable direct comparability with the previous analysis of this set of candidate migration genes across a broader set of avian species (Lugo Ramos et al. 2017).

We first asked whether selective pressure shifted along lineages where transitions in migratory behavior likely occurred. To evaluate this, we fit “branch” models which allow ω to vary between selected focal lineages (“foreground” branches) and the remaining branches of the tree (“background” branches; Supplementary Table S5). For the “migratory branch” model, we set as the foreground four branches where—inferred from the ancestral character state reconstruction—a migratory clade likely evolved from a sedentary “background” (Figure 1; Supplementary Table S6). While we are primarily interested in selection associated with the gain of migration, selective pressures may also accompany its loss, for example through the need to adapt to different environmental conditions or transition to different circadian rhythms (Winger et al. 2019). We therefore also fit “sedentary branch” models that designated seven sedentary lineages as foreground branches (Figure 1; Supplementary Table S6). We used likelihood ratio tests (LRTs) to determine whether the branch models provided a significantly better fit to the data than the null m0 model, which holds ω constant among branches (Supplementary Table S5).

In order to assess whether specific amino acid sites have been subject to positive selection related to transitions in migratory behavior, we fit branch-site models (migratory foreground and sedentary foreground), which—in addition to allowing ω to vary between foreground and background branches—allow ω to vary between sites (Supplementary Table S5). The branch-site model classifies sites into four classes: class 0, where sites are under purifying selection (ω < 1) on all branches; class 1, where sites evolve neutrally (ω = 1) on all branches; and classes 2a and 2b, where sites that are constrained (ω < 1) or neutral (ω = 1) on background branches are under positive selection (ω > 1) on foreground branches. We then used LRTs to determine whether the branch-site model including a positively selected class better fit the data than a null branch-site model in which the foreground ω for classes 2a and 2b was fixed at 1 (Supplementary Table S5). When the branch-site model provided a significantly better fit than the null model, we used Bayes Empirical Bayes (BEB) tests to identify which amino acid sites had a high posterior probability (P > 0.95) of belonging to the positively selected site class (Yang et al. 2005).

Finally, we fit site models, which allow ω to vary among sites but not among branches, to determine whether any candidate genes showed evidence of positive selection independent of transitions in migratory phenotype. We compared a “positive selection” (M2a) site model allowing three site classes (ω < 1, ω = 1, and ω > 1) with a null “nearly neutral” (M1a) site model that did not include a positively selected class (ω < 1, ω = 1) using LRTs. As with the branch-site models, we used BEB tests to identify which amino acid sites had a high posterior probability of belonging to the positively selected site class. For putatively selected sites, we mapped amino acid variants across the relevant gene tree using phytools and performed ancestral character state reconstruction with corHMM as described earlier to infer when amino acid transitions occurred and whether they aligned with transitions in migratory behavior.

For all CODEML models, we set CodonFreq = 7 (mutation-selection model; Yang and Nielsen 2008), estFreq = 0 (estimate codon frequencies from the data), clock = 0 (no clock, independent rates per branch), and cleandata = 0 (do not remove sites with ambiguity codes or gaps). To mitigate the risk of convergence on local likelihood optima, we ran each model twice with different starting ω values (ω = 0.5 and ω = 2). We labeled foreground branches on the parulid warbler species tree in Dendroscope (Huson et al. 2007). For this analysis, we used the species tree rather than individual gene trees because our question concerns trait transitions during speciation, and the gene trees showed topological discordance with the species tree that was likely driven by variation in the number of phylogenetically informative sites rather than true evolutionary signal. To account for multiple testing, we first converted chi-square values from the LRTs to *p*-values and then applied a Bonferroni correction, which has been found to effectively reduce Type I error without dramatically reducing power in a simulation study of CODEML analyses (Anisimova and Yang 2007).

## Results

We constructed maximum likelihood phylogenies for 25 candidate migration genes and compared the topologies of these gene trees to the topology of the parulid warbler species tree by generating cophylogeny plots. The absolute Jaccard Robinson-Foulds (JRF) distance between the gene trees and the species tree ranged from 27.6–68.8 (sd = 12.9), with a median of 39.4 and the normalized JRF ranged from 0.14–0.35, with a median of 0.20 (sd = 0.07; Figure 2A; Supplementary Table S3). The topologies of most candidate gene trees were largely concordant with the topology of the species tree (e.g., *CPNE4*; Supplementary Figure S1). In cases where regions of a gene tree were discordant with the species tree (e.g., *DRD4*; Supplementary Figure S2), we did not visually observe increased phylogenetic clustering of species sharing migratory phenotypes.

**FIGURE 2.**
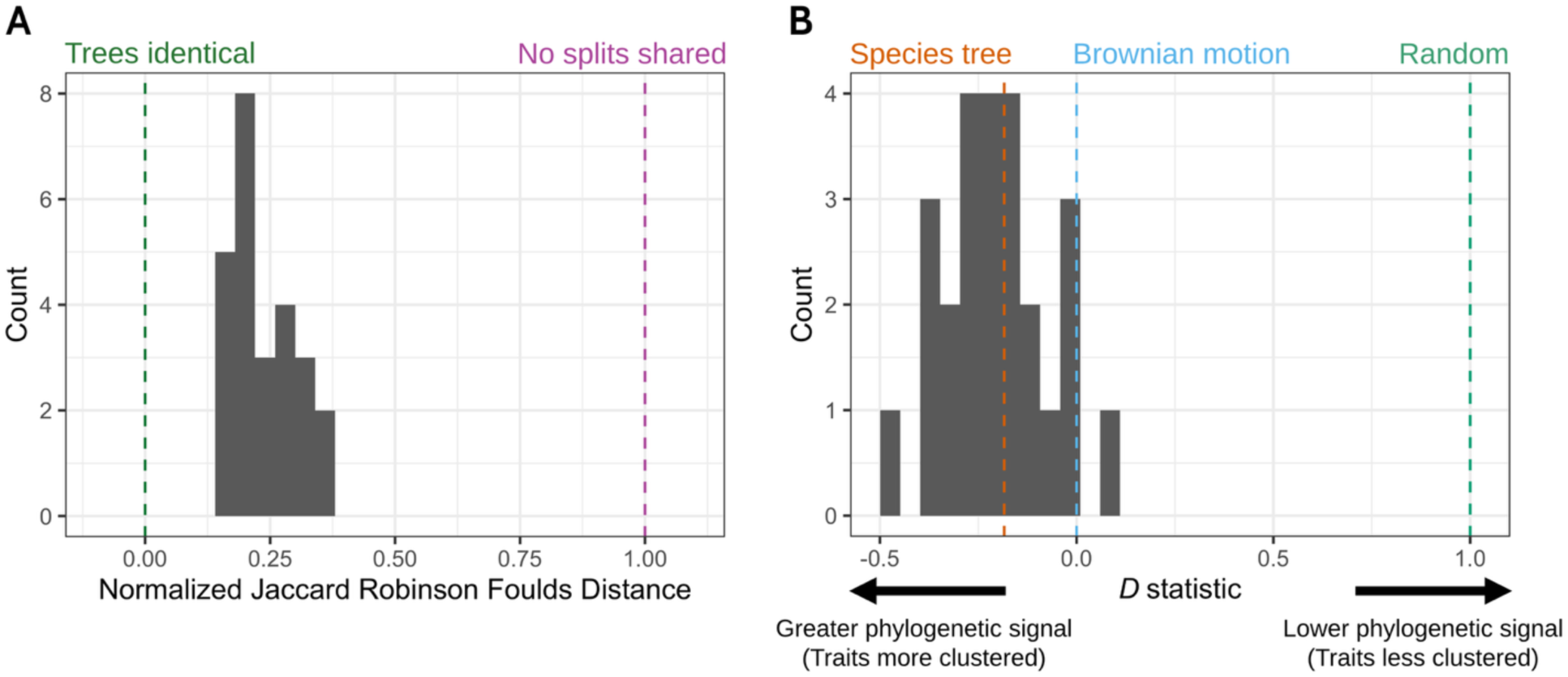
Histograms of topological discordance and phylogenetic signal across 25 candidate migration gene trees relative to the parulid warbler species tree. (A) Normalized Jaccard Robinson-Foulds (JRF) distances between each gene tree and the species tree topology. When two compared phylogenies are identical, JRF = 0 (green dotted line), and when the trees share no identical splits, JRF = 1 (purple dotted line). (B) Histogram of the *D* statistic (Fritz and Purvis 2010) measuring phylogenetic signal of migratory propensity (i.e., sedentary vs. migratory) in each candidate gene tree. When the distribution of traits over the tips of the tree follows a found no obvious increase of clustering of migratory species in the gene trees compared to the species tree. Whereas we found that gene tree discordance was not attributable to increased genetic similarity among migratory species at candidate migration genes, we observed that JRF distance decreased logarithmically with both increasing gene length and the number of parsimony informative sites (Supplementary Figure S3).

To quantify the clustering of species by migratory phenotype in the gene trees compared to the species tree, we calculated the *D* statistic of Fritz and Purvis (2010), a measure of phylogenetic signal in a binary trait. Migratory propensity exhibited significant phylogenetic signal in both the parulid species tree and all candidate gene trees. The trait topology of the species tree (*D* = -0.184) significantly deviated from a random phylogenetic pattern (*P* < 0.0001) and was not significantly different from expectations of the Brownian motion model (*P* = 0.803; Figure 2B). For the candidate gene trees, *D* ranged from -0.474–0.081, with an average of -0.207 (Figure 2B). All gene trees were significantly different from the expectations of the random phylogenetic model, but not the Brownian motion model (Supplementary Table S3). The gene tree *D* values were normally distributed (Shapiro-Wilk *W* = 0.987, *P* = 0.984) and approximately centered around the value of *D* for the species tree (Figure 2B). Though 14 candidate gene trees had lower *D* values than the species tree (i.e., greater phylogenetic signal), visual inspection completely random pattern, *D* = 1 (dotted green line), and when the phylogenetic pattern of the traits fits a Brownian motion model, *D* = 0 (dotted blue line). The *D* value for the parulid warbler species tree is shown by the dotted orange line (*D* = -0.184).

The results of the ancestral character state reconstruction were largely consistent with previous studies (Winger et al. 2012; Mitchell et al. 2019), confirming that the most recent common ancestor of the parulid warblers was most likely migratory (posterior probability of migratory state at the root node = 0.854) and that transitions between migratory and sedentary behavior have occurred many times throughout parulid warbler diversification (Figure 1). The equal rates model was the best fitting model of discrete character evolution (transition rate = 0.813; Supplementary Table S4), suggesting that transitions from migratory to sedentary behavior and vice versa have occurred at roughly equal rates. In several cases, migratory behavior was lost and subsequently regained. For example, migration was likely lost in the ancestor of the clade containing *Myiothlypis*, *Basileuterus*, *Myioborus*, and *Cardellina* (P_sedentary_ = 0.91) or in the directly preceding node (P_sedentary_ = 0.52) but was then regained in the ancestor of *Cardellina pusilla* and *C. canadensis* and certain populations of *C. rubrifrons* and *Myioborus pictus*. Informed by the inferred ancestral states, we selected foreground branches for analyzing coding sequence evolution in CODEML where either a migratory clade arose from a sedentary “background” (i.e., migratory foreground branch) or a sedentary clade arose from a migratory background (sedentary foreground branch; Figure 1, Supplementary Table S5).

Among the coding sequences of the 25 candidate migration genes, the global nonsynonymous substitution rate (dN) ranged from 0.000–0.096 (mean = 0.017, median = 0.011), the synonymous substitution rate (dS) ranged from 0.019–0.652 (mean = 0.314, median = 0.278), and dN/dS ranged from 0.000–0.279 (mean = 0.059, median = 0.029; Table 1). After applying a Bonferroni correction for multiple comparisons, no branch model—regardless of whether migratory or sedentary branches were designated as the foreground—fit the data significantly better than the one-ratio m0 model that assumed a single dN/dS (ω) across all branches (Table 1). Similarly, no migratory or sedentary branch-site model allowing positive selection on foreground branches provided a significantly better fit than the corresponding null model, which constrained ω ≤ 1.

**TABLE 1.**
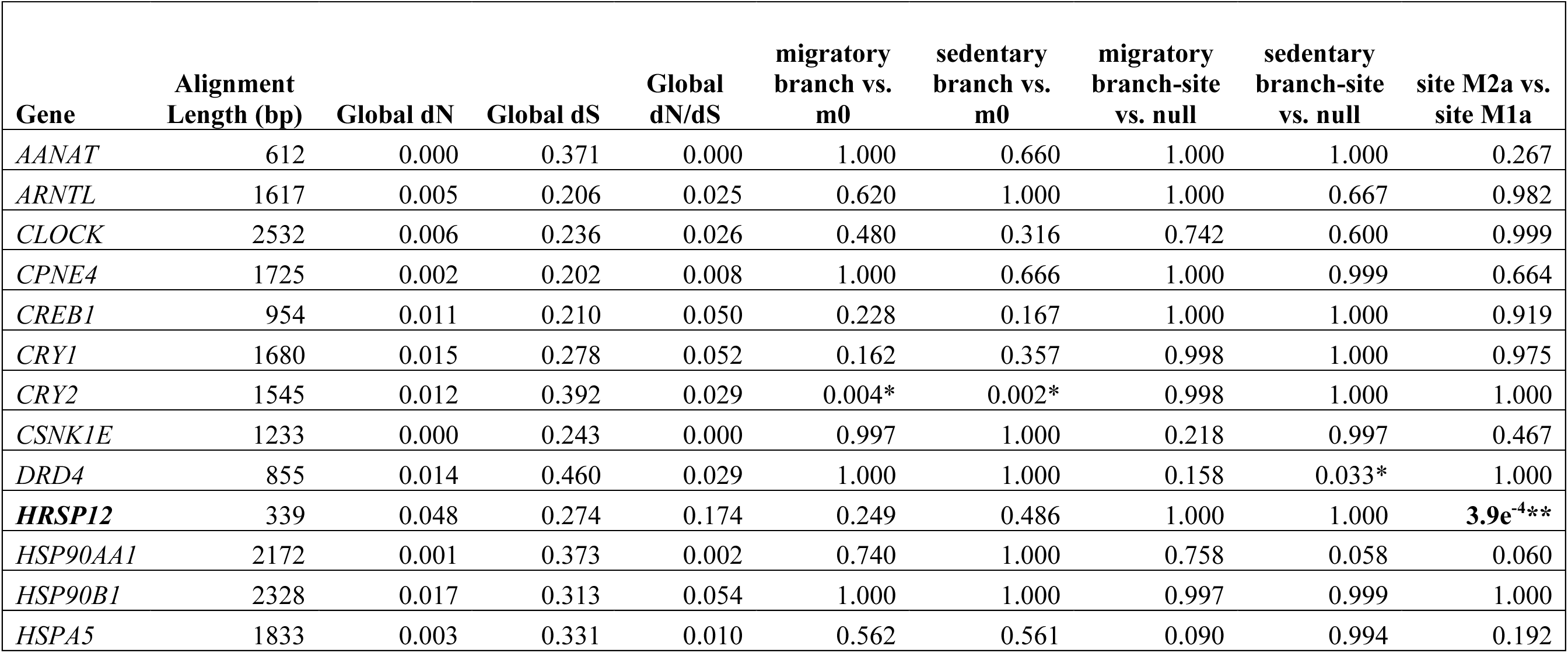
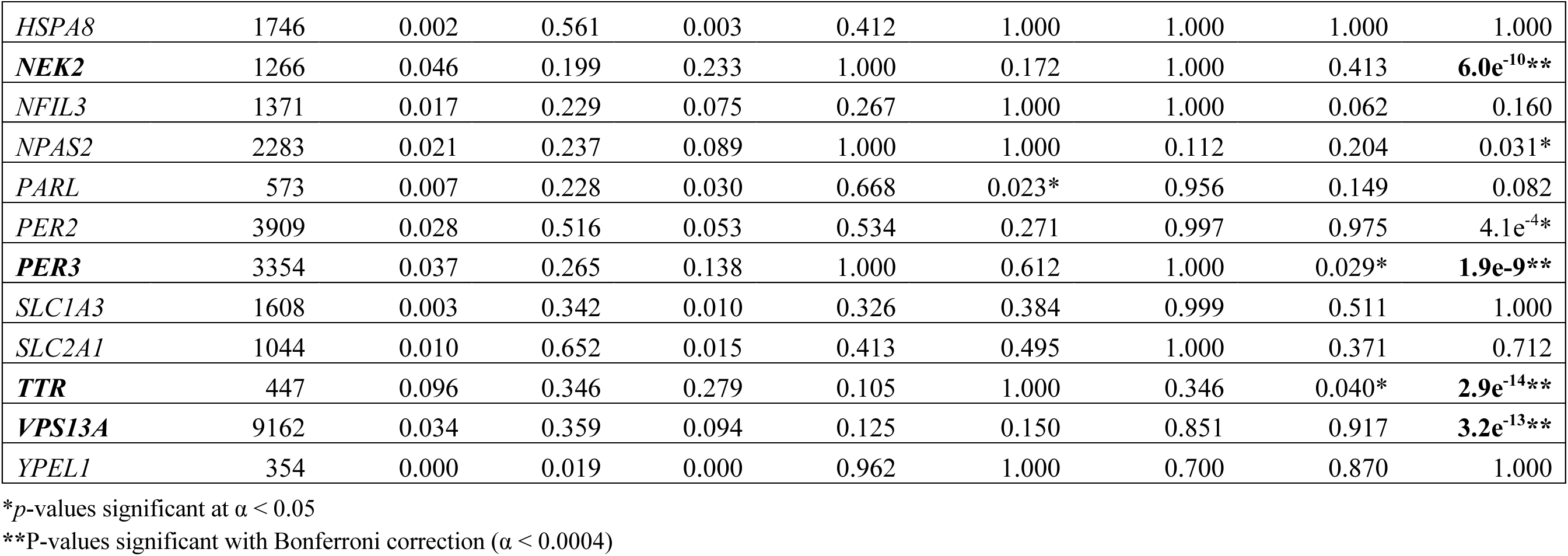
Results of analyses of coding sequence evolution for candidate migration genes across the parulid warbler radiation. Reported for each gene are the length of the alignment (bp), global dN (nonsynonymous substitution rate), dS (synonymous substitution rate) and dN/dS (ω). We report *p*-values for likelihood ratio tests comparing CODEML models of sequence evolution. These comparisons include (1) branch models with either migratory or sedentary foreground branches vs. the one-ratio model (m0), which assumes a single ω across all branches and sites; (2) branch-site models allowing positive selection on foreground branches vs. null models constraining ω ≤ 1; and (3) site models allowing positive selection (M2a) vs. null models constraining ω ≤ 1 (M1a). *P*- values for likelihood ratio tests significant after correcting for multiple testing area highlighted in bold.

| Gene | Alignment Length (bp) | Global dN | Global dS | Global dN/dS | migratory branch vs. m0 | sedentary branch vs. m0 | migratory branch-site vs. null | sedentary branch-site vs. null | site M2a vs. site M1a |
|---|---|---|---|---|---|---|---|---|---|
| *AANAT* | 612 | 0.000 | 0.371 | 0.000 | 1.000 | 0.660 | 1.000 | 1.000 | 0.267 |
| *ARNTL* | 1617 | 0.005 | 0.206 | 0.025 | 0.620 | 1.000 | 1.000 | 0.667 | 0.982 |
| *CLOCK* | 2532 | 0.006 | 0.236 | 0.026 | 0.480 | 0.316 | 0.742 | 0.600 | 0.999 |
| *CPNE4* | 1725 | 0.002 | 0.202 | 0.008 | 1.000 | 0.666 | 1.000 | 0.999 | 0.664 |
| *CREB1* | 954 | 0.011 | 0.210 | 0.050 | 0.228 | 0.167 | 1.000 | 1.000 | 0.919 |
| *CRY1* | 1680 | 0.015 | 0.278 | 0.052 | 0.162 | 0.357 | 0.998 | 1.000 | 0.975 |
| *CRY2* | 1545 | 0.012 | 0.392 | 0.029 | 0.004* | 0.002* | 0.998 | 1.000 | 1.000 |
| *CSNK1E* | 1233 | 0.000 | 0.243 | 0.000 | 0.997 | 1.000 | 0.218 | 0.997 | 0.467 |
| *DRD4* | 855 | 0.014 | 0.460 | 0.029 | 1.000 | 1.000 | 0.158 | 0.033* | 1.000 |
| ***HRSP12*** | 339 | 0.048 | 0.274 | 0.174 | 0.249 | 0.486 | 1.000 | 1.000 | **3.9e<sup>-4</sup>\*\*** |
| *HSP90AA1* | 2172 | 0.001 | 0.373 | 0.002 | 0.740 | 1.000 | 0.758 | 0.058 | 0.060 |
| *HSP90B1* | 2328 | 0.017 | 0.313 | 0.054 | 1.000 | 1.000 | 0.997 | 0.999 | 1.000 |
| *HSPA5* | 1833 | 0.003 | 0.331 | 0.010 | 0.562 | 0.561 | 0.090 | 0.994 | 0.192 |
| *HSPA8* | 1746 | 0.002 | 0.561 | 0.003 | 0.412 | 1.000 | 1.000 | 1.000 | 1.000 |
| ***NEK2*** | 1266 | 0.046 | 0.199 | 0.233 | 1.000 | 0.172 | 1.000 | 0.413 | **6.0e<sup>-10</sup>\*\*** |
| *NFIL3* | 1371 | 0.017 | 0.229 | 0.075 | 0.267 | 1.000 | 1.000 | 0.062 | 0.160 |
| *NPAS2* | 2283 | 0.021 | 0.237 | 0.089 | 1.000 | 1.000 | 0.112 | 0.204 | 0.031* |
| *PARL* | 573 | 0.007 | 0.228 | 0.030 | 0.668 | 0.023* | 0.956 | 0.149 | 0.082 |
| *PER2* | 3909 | 0.028 | 0.516 | 0.053 | 0.534 | 0.271 | 0.997 | 0.975 | 4.1e<sup>-4</sup>* |
| ***PER3*** | 3354 | 0.037 | 0.265 | 0.138 | 1.000 | 0.612 | 1.000 | 0.029* | **1.9e-9\*\*** |
| *SLC1A3* | 1608 | 0.003 | 0.342 | 0.010 | 0.326 | 0.384 | 0.999 | 0.511 | 1.000 |
| *SLC2A1* | 1044 | 0.010 | 0.652 | 0.015 | 0.413 | 0.495 | 1.000 | 0.371 | 0.712 |
| ***TTR*** | 447 | 0.096 | 0.346 | 0.279 | 0.105 | 1.000 | 0.346 | 0.040* | **2.9e<sup>-14</sup>\*\*** |
| ***VPS13A*** | 9162 | 0.034 | 0.359 | 0.094 | 0.125 | 0.150 | 0.851 | 0.917 | **3.2e<sup>-13</sup>\*\*** |
| *YPEL1* | 354 | 0.000 | 0.019 | 0.000 | 0.962 | 1.000 | 0.700 | 0.870 | 1.000 |

\**p*-values significant at α < 0.05
\*\*P-values significant with Bonferroni correction (α < 0.0004)

For five candidate genes (*HRSP12*, *NEK2*, *PER3*, *TTR*, and *VPS13A*), the site model allowing positively selected sites (ω > 1; M2a) provided a significantly better fit than the nearly neutral model (M1a), which restricts all sites to ω ≤ 1 (Table 1). Bayes Empirical Bayes analysis identified 1–8 amino acid sites per gene (mean = 4.2) with posterior probability > 0.95 of belonging to the positively selected site class (ω > 1; Supplementary Table S6). The proportion of sites inferred to be under positive selection ranged from 0.006 to 0.02 (mean = 0.01) and estimated ω values for this positively selected site class ranged from 2.5 to 9.8, with a mean of 5.3 (Supplementary Table S6). Examination of the amino acid variants present at these sites across parulid warbler species revealed no clear association between the variants and migratory propensity, and the sites were often highly polymorphic across the entire phylogeny (Figure 3, Supplementary Figures S27–S47). These results indicate that while positive selection may have acted on specific sites within these five candidate genes, it was not concentrated on branches where transitions in migratory behavior occurred and thus was most likely the product of lineage-specific evolution unrelated to migratory phenotype.

**FIGURE 3.**
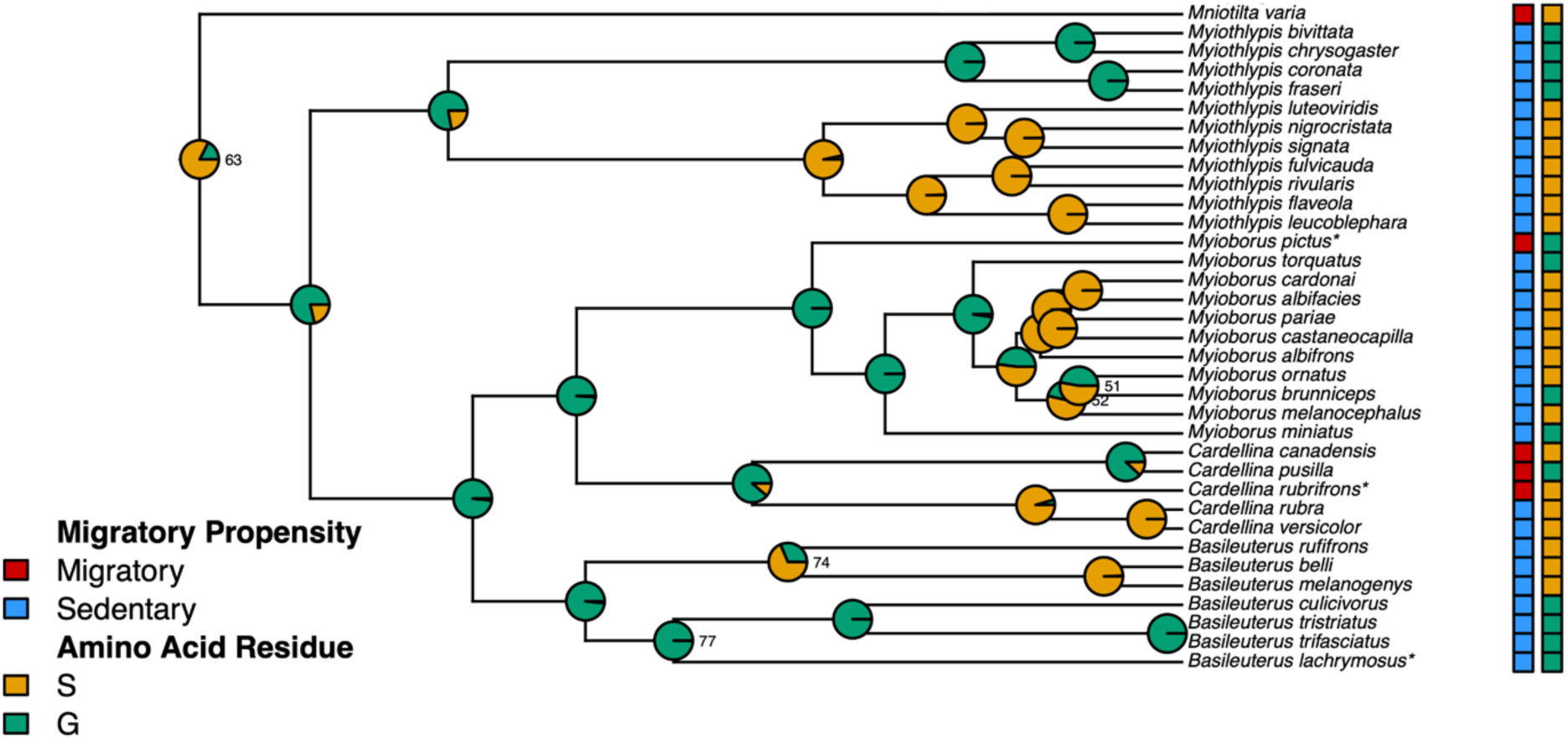
Amino acid variation at *PER3* site 452 mapped across the gene tree for the clade containing *Basileuterus*, *Cardellina*, *Mniotilta*, *Myioborus*, and *Myiothlypis*. Square labels at tips represent migratory propensity (red: migratory, blue: sedentary) and amino acid residue (orange: serine, green: glycine) for each species. Pie charts on nodes show ancestral character state reconstruction posterior probabilities for each amino acid variant. Nodes of the tree that received <80% support by approximate likelihood ratio test (aLRT) are labeled with their aLRT value. Partially migratory species are indicated by asterisks next to their species name. Outside of these lineages, serine was fixed at this site across all remaining parulid warblers. Although this site showed strong evidence of positive selection (significantly better fit of the M2a model over M1a and BEB posterior probability of 0.999), none of the branch or branch-site models testing for rate shifts tied to migratory transitions fit better than the null model.

## Discussion

We characterized variation in 25 candidate migration genes across 101 parulid warbler taxa using whole-genome resequencing data and an updated UCE phylogeny of the family. Comparing candidate migration gene trees to the species tree revealed elevated discordance in some genes that was associated with alignment length but not driven by the clustering of species based on migratory strategy. This pattern suggests that discordance stemmed from limited phylogenetically informative sites in shorter genes, rather than from selection on shared variation among species with similar migratory strategies. Testing models of coding sequence evolution in PAML indicated that most candidate migration genes have been subject to strong purifying selection. We identified sites with significantly elevated dN/dS in five candidate genes, but variation at these sites was not associated with differences in migratory propensity and we did not detect repeated selection occurring across lineages where transitions in migratory behavior likely occurred. Taken together, these results contradict the hypothesis that parallel molecular evolution in these candidate migration genes facilitated the repeated evolution of migratory behavior across the parulid warbler radiation.

### Gene Tree Discordance Does Not Relate to Migratory Strategy

When the same genetic variants drive repeated phenotypic evolution, gene trees constructed from those loci should show taxa that share phenotype clustering together more closely than their phylogenetic relationships would predict. Though this pattern could arise due to repeated selection on shared ancestral polymorphisms, in closely related species that retain reproductive compatibility among lineages this can also result from historical adaptive introgression between species (Jones et al. 2012, 2018). Such clustering has been documented in parulid warblers for the pigmentation genes *BCO2*, *BDH1L*, and *CYP2J19*, indicating that introgression likely facilitated the repeated evolution of yellow and red coloration in this group (Baiz et al. 2021; Bennett et al. 2025). Despite a well-documented history of hybridization and frequent transitions between migratory and sedentary behavior in the parulid warblers, we observed no greater clustering by migratory phenotype than expected based on phylogenetic relationships.

Migratory propensity exhibited strong phylogenetic signal in both the UCE species tree and all candidate gene trees, corroborating previous estimates of phylogenetic signal for migratory behavior in parulid warblers derived using different methods (Winger et al. 2012; Lugo Ramos et al. 2017; Mitchell et al. 2019). However, we did not observe significantly higher phylogenetic signal (i.e. increased clustering by migratory phenotype) in the candidate migration gene trees compared to the species tree, even in cases of strong topological discordance. The observed discordance between gene trees and the species tree may be more readily explained by gene flow unrelated to migratory phenotype or by incomplete lineage sorting, which is expected to be substantial in rapidly diversifying clades like the parulid warblers (Suh et al. 2015; Oliveros et al. 2019). However, our finding that gene tree discordance was correlated with alignment length—with the shortest genes exhibiting the highest Jaccard-Robinson Foulds distances— suggests that this discordance primarily stemmed from uncertainty in gene tree estimation due to limited informative sites.

Our results support the findings of Lugo Ramos et al. (2017) that migratory species across all orders of birds do not share more variation at candidate migration genes than expected based on phylogenetic relationships. By examining the same question in a radiation of closely related species, we show that the result of this previous study was not simply a function of its broad taxonomic sampling. While the highly divergent species included in Lugo Ramos et al. (2017) likely employ different molecular and physiological mechanisms to facilitate migration (Piersma et al. 2005; Zink 2011), it is more likely that parulid warblers rely on a similar migratory program. That we did not observe shared variation between divergent migratory warblers implies that the repeated evolution of migration is unlikely to rely on identical variation at the nucleotide level within these candidate migration genes. Though Lugo Ramos et al. reported increased clustering by migratory phenotype in two genes (*HRSP12* and *HSPA4*) when their analysis was restricted to passerines, we observed no such pattern in these genes in the parulid warblers. This suggests that the pattern observed previously in these genes may have resulted from gene tree discordance clustering migratory and sedentary species by chance rather than because shared variation found in these genes facilitates migratory behavior.

### Elevated Rates of Coding Sequence Evolution Are Independent of Migratory Phenotype

Analyzing models of coding sequence evolution in CODEML revealed signs of strong purifying selection across candidate migration genes, with most sites exhibiting dN/dS (ω) values near zero. This pattern is expected for protein coding genes, since non-synonymous mutations are typically deleterious due to their effect on amino acid sequences (Ohta 1995). However, we found that the site model including positively selected sites (ω > 1) fit the data significantly better than the nearly neutral model for five candidate genes: *HRSP12*, *NEK2*, *PER3*, *TTR*, and *VPS13A*. These genes span a range of functions, including cellular stress response, control of circadian rhythms, cell cycle regulation, and hormone and lipid transport (Supplementary Table S2).

In other species, *HRSP12* and *PER3* have been associated with migratory propensity in common yellowthroats (*Geothlypis trichas*; (Zamudio-Beltrán et al. 2024), *VPS13A* varies with migratory distance and direction in *Vermivora* warblers (Toews et al. 2019), and *HRSP12* and *TTR* shows differential expression across migratory states in white-crowned sparrows (*Zonotrichia leucophrys*; (Jones et al. 2008). In contrast, *NEK2* remains a putative candidate migration gene without direct evidence of functional association with migratory behavior (Ruegg et al. 2014). Lugo Ramos et al. (2017) also found evidence of positive selection in *NEK2*, but not in *HRSP12*, *PER3*, or *TTR* (*VPS13A* was not included in their analysis).

Although these genes each contained sites that exhibited elevated dN/dS, the patterns of variation at these sites showed no obvious association with migratory phenotype. Reflecting this, none of the branch or branch-site models testing for elevated rates of sequence evolution on migratory lineages arising from sedentary backgrounds (or vice versa) fit the data significantly better than the single rate model. Furthermore, examination of variants at putatively selected sites revealed no clear association with migratory propensity across taxa. Instead, most of the sites were highly polymorphic across the entire phylogeny. This suggests that elevated dN/dS at these sites may result from relaxed purifying selection or balancing selection rather than repeated positive selection across the phylogeny.

We performed dN/dS analysis for comparability with Lugo Ramos et al. (2017), though this approach is conservative and has inherent limitations. First, dN/dS is calculated from fixed substitutions, so recent or weak selection on variants that have not reached fixation may go undetected, especially in recently diverged taxa (Kryazhimskiy and Plotkin 2008). Secondly, because models of coding sequence evolution average dN/dS across either branches or sites, a signal of positive selection can be suppressed by a strong background of purifying selection. However, we performed dN/dS analysis despite these limitations because using a different methodological approach could obscure whether discrepancies between studies stemmed from genuine biological differences in our chosen focal taxa or from differences in analytical power.

Future work could leverage methods that not only are less conservative than dN/dS but also are applicable to non-coding regions. The lack of coding sequence variation associated with transitions in migratory behavior that we observed in this study is consistent with the hypothesis that the evolution of migration is largely driven by variation in non-coding regions affecting gene regulation (Caballero-Lopez and Bensch 2024). Variation in non-coding regions has been associated with the repeated evolution of ecotypes in three-spined sticklebacks (Jones et al. 2012; O’Brown et al. 2015) and pigmentation in parulid warblers (Toews et al. 2016; Baiz et al. 2021), and some previously identified variation associated with intraspecific migratory differences is located within non-coding regions (Delmore et al. 2020b; Caballero-López et al. 2022; Sokolovskis et al. 2023). Approaches allowing for more flexible models of sequence evolution could detect variation in regulatory regions that may affect migratory phenotype (Kowalczyk et al. 2019; Yan et al. 2023).

## Conclusion

We found that while migratory behavior has been lost and regained repeatedly across the parulid warbler radiation, there is no evidence that this was facilitated by the same genetic variants, either through repeated selection acting on shared variants, or through adaptive introgression from interspecific gene flow. These results support the findings of Lugo Ramos et al. (2017), who found no associations between variation at candidate migration genes in 70 species from all avian orders. By analyzing these same genes at a much narrower taxonomic scale, we show that the negative result of Lugo Ramos et al. (2017) likely was not simply the result of broad taxonomic sampling. Instead, the repeated evolution of migratory behavior is unlikely to rely on the same genetic changes within migration genes. An alternative explanation may be that variation in the same genes—or the same molecular pathways—facilitates migratory behavior, but different genetic variants have similar effects. Another hypothesis, which has been proposed due to the rapidity of changes in migratory behavior, is that changes in regulatory regions play the primary role in controlling migratory propensity (reviewed in Caballero-Lopez and Bensch 2024). Despite limited work on gene regulation in non-model birds, connecting regulatory variation to migratory phenotype is a promising future avenue for understanding the genetic architecture of seasonal migration.

## Acknowledgments

We thank Jill Hamilton, George Perry, and Zachary Szpiech for feedback and helpful discussion. Many thanks to Andrew Wood for obtaining and sequencing samples. We are grateful to Alan Brelsford for providing the *Setophaga coronata* genome annotation used in this study. We thank the American Museum of Natural History, Cornell University Museum of Vertebrates, Field Museum of Natural History, University of Kansas Natural History Museum, Louisiana State University of Natural Science, Museum of Southwestern Biology, Berkeley Museum of Vertebrate Zoology, Smithsonian Migratory Bird Center, Smithsonian Tropical Research Institute, University of California Riverside, and University of Washington Burke Museum for blood and tissue loans. Computations for this project were performed on Pennsylvania State University’s Institute for Computational and Data Sciences’ Roar Collab computing cluster (RRID:SCR_025154) and the original genomic data used in the analysis was generated at the Pennsylvania State University Huck Institutes of the Life Sciences Genomics Core Facility (RRID:SCR_023645) and the Biotechnology Resource Center Genomics Facility at the Cornell Institute of Biotechnology (RRID:SCR_021727).

## Data Availability Statement

The raw whole-genome sequencing reads used in this study are available under NCBI BioProject PRJNA630247. The *Setophaga coronata* reference genome is available under NCBI BioProject PRJNA325157. All data and code from this study, including candidate gene alignments and gene trees, are published on the Dryad Digital Repository: http://datadryad.org/share/LINK_NOT_FOR_PUBLICATION/hX-MyGanIl9EfufUgAFkMB_XzKEEcF5NJWNEhmp2dLY.

