## Supplementary Material for "Repeated Transitions in Migratory Behavior Across a Songbird Radiation Are Not Driven by Selection on Shared Variation in Candidate Genes"

**Table S1:** Parulid warbler taxa included in study, with sample size per taxon and assigned migratory status .49–53

**Table S2:** Candidate genes included in study, with genomic position, functional category, and citations .......54–55

**Table S3:** Results from comparisons of candidate migration gene tree topologies vs. the species tree............56–57

### SUPPLEMENTARY FIGURES

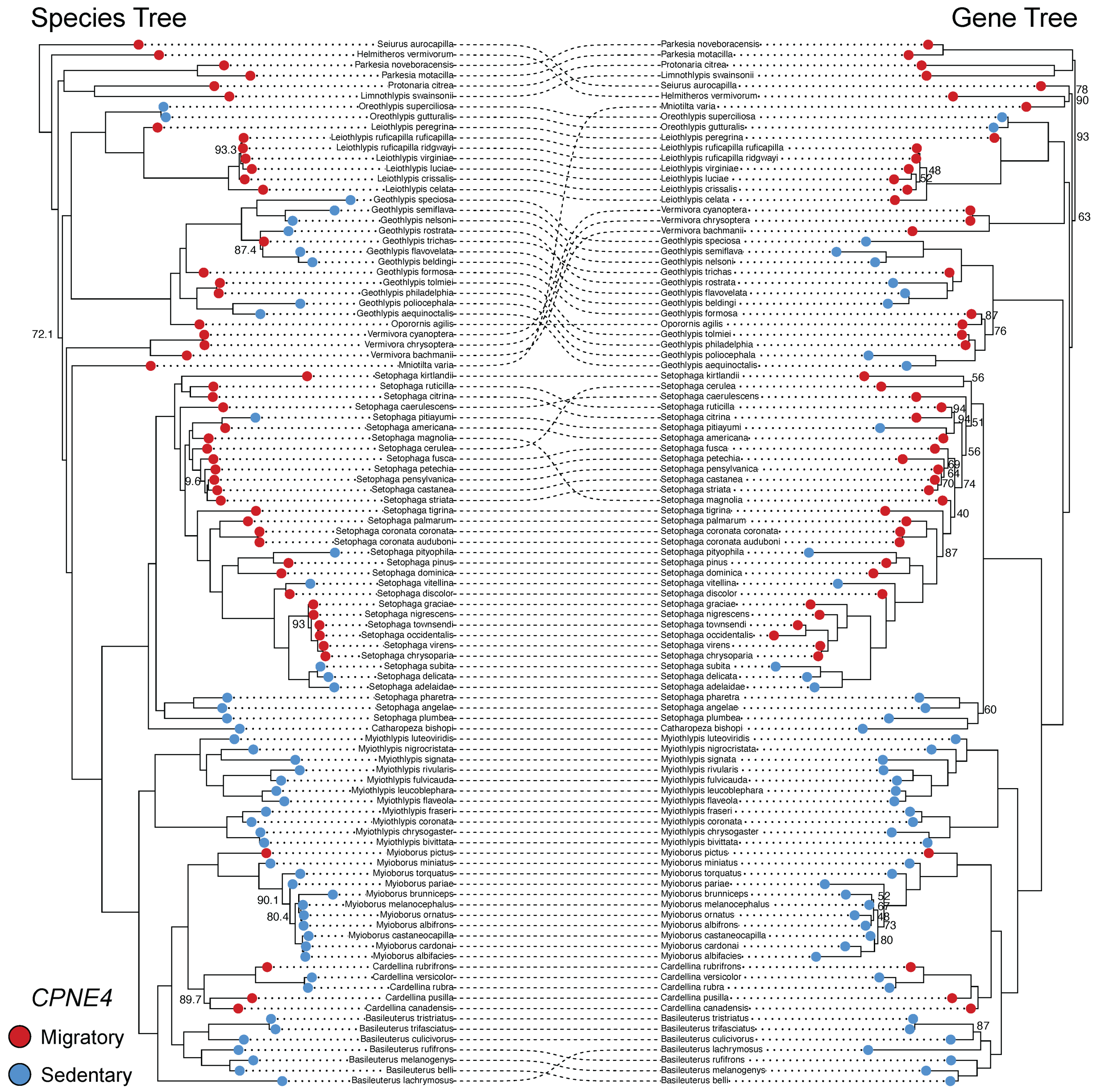

**FIGURE S1. Cophylogeny visualizing regions of concordance and discordance between the topology of a candidate migration gene tree (*CPNE4*; right) and the parulid warbler species tree (left).** *CPNE4* exhibited the least discordance of any candidate gene quantified by the normalized Jaccard-Robinsons Foulds distance (nJRF = 0.14). Dotted lines connect species between the two trees: straight or parallel lines indicate identical placement in both trees, while curved intersecting lines indicate regions of discordance. Red circles at the tips of each tree represent migratory species and blue circles represent sedentary species. Bootstrap values <95 are shown at nodes.

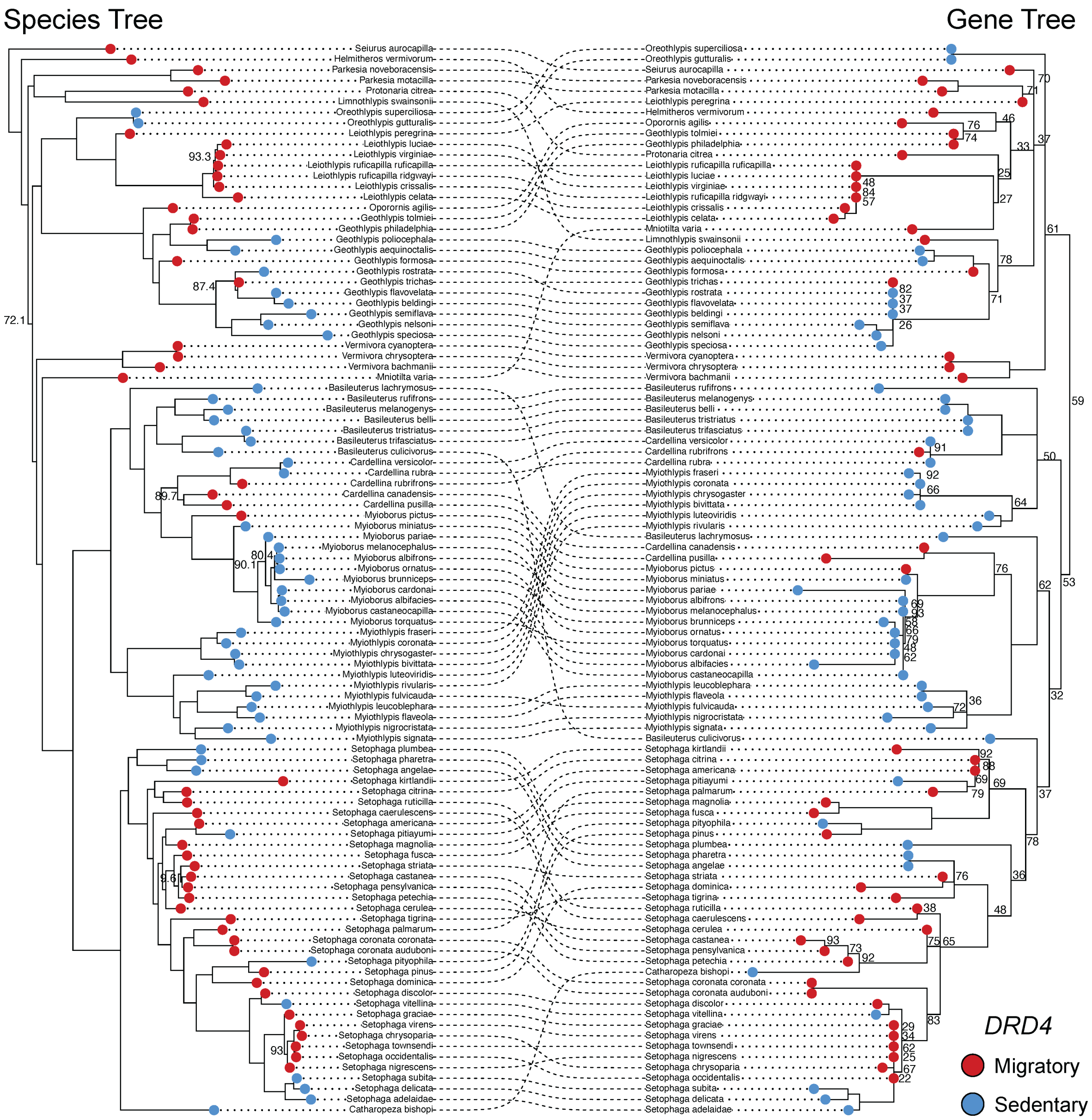

**FIGURE S2. Cophylogeny visualizing regions of concordance and discordance between the topology of a candidate migration gene tree (*DRD4*; right) and the parulid warbler species tree (left).** *DRD4* exhibited the highest discordance of any candidate gene quantified by the normalized Jaccard-Robinsons Foulds distance (nJRF = 0.35). Dotted lines connect species between the two trees: parallel lines indicate identical placement in both trees, while curved intersecting lines indicate regions of discordance. Red circles at the tips of each tree represent migratory species and blue circles represent sedentary species. Bootstrap values <95 are shown at nodes.

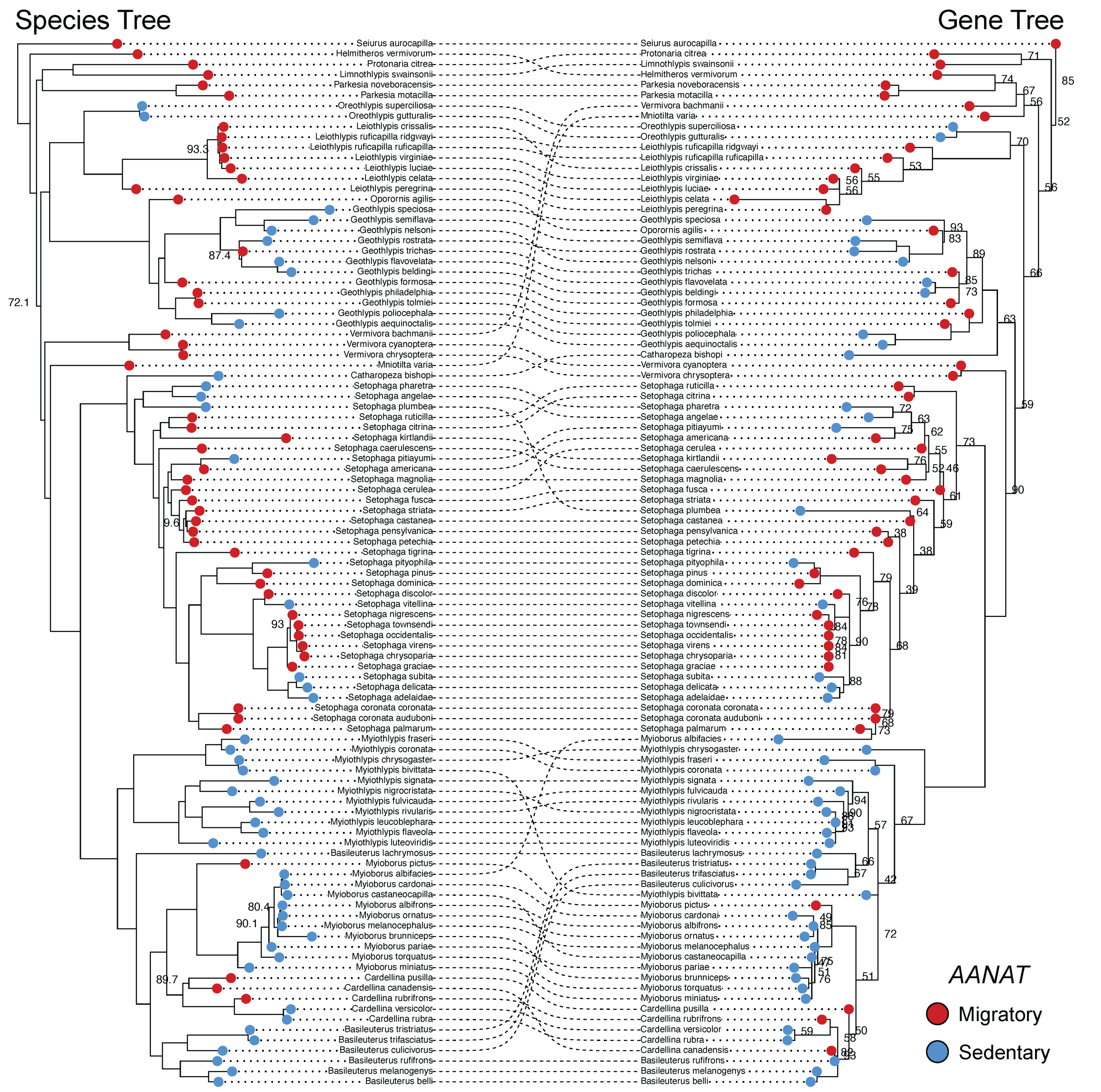

**FIGURE S3. Cophylogeny visualizing regions of concordance and discordance between the topology of a candidate migration gene tree (*AANAT*; right) and the parulid warbler species tree (left).** Dotted lines connect species between the two trees: parallel lines indicate identical placement in both trees, while curved intersecting lines indicate regions of discordance. Red circles at the tips of each tree represent migratory species and blue circles represent sedentary species. Bootstrap values <95 are shown at nodes.

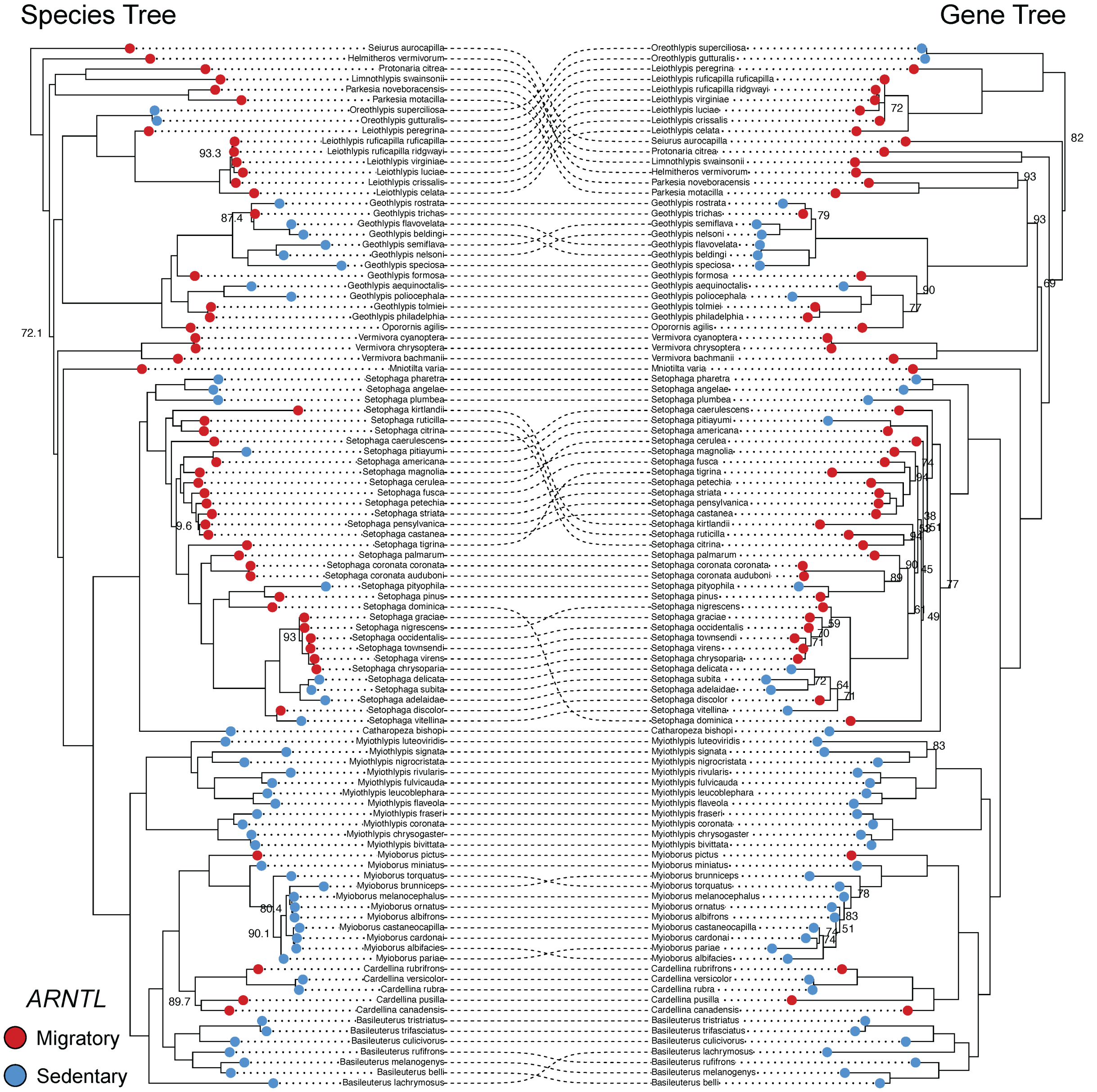

**FIGURE S4. Cophylogeny visualizing regions of concordance and discordance between the topology of a candidate migration gene tree (*ARNTL*; right) and the parulid warbler species tree (left).** Dotted lines connect species between the two trees: parallel lines indicate identical placement in both trees, while curved intersecting lines indicate regions of discordance. Red circles at the tips of each tree represent migratory species and blue circles represent sedentary species. Bootstrap values <95 are shown at nodes.

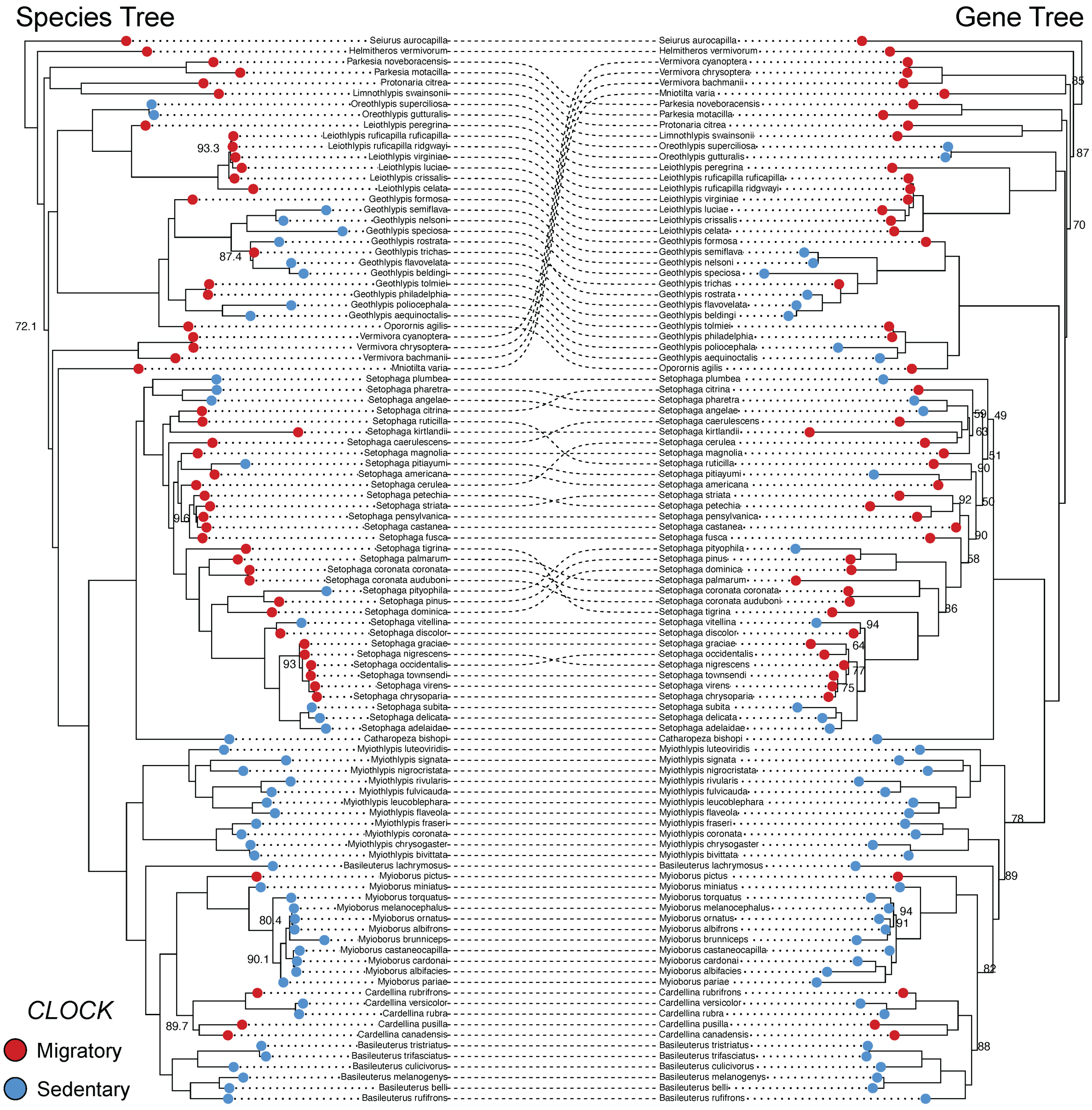

**FIGURE S5. Cophylogeny visualizing regions of concordance and discordance between the topology of a candidate migration gene tree (*CLOCK*; right) and the parulid warbler species tree (left).** Dotted lines connect species between the two trees: parallel lines indicate identical placement in both trees, while curved intersecting lines indicate regions of discordance. Red circles at the tips of each tree represent migratory species and blue circles represent sedentary species. Bootstrap values <95 are shown at nodes.

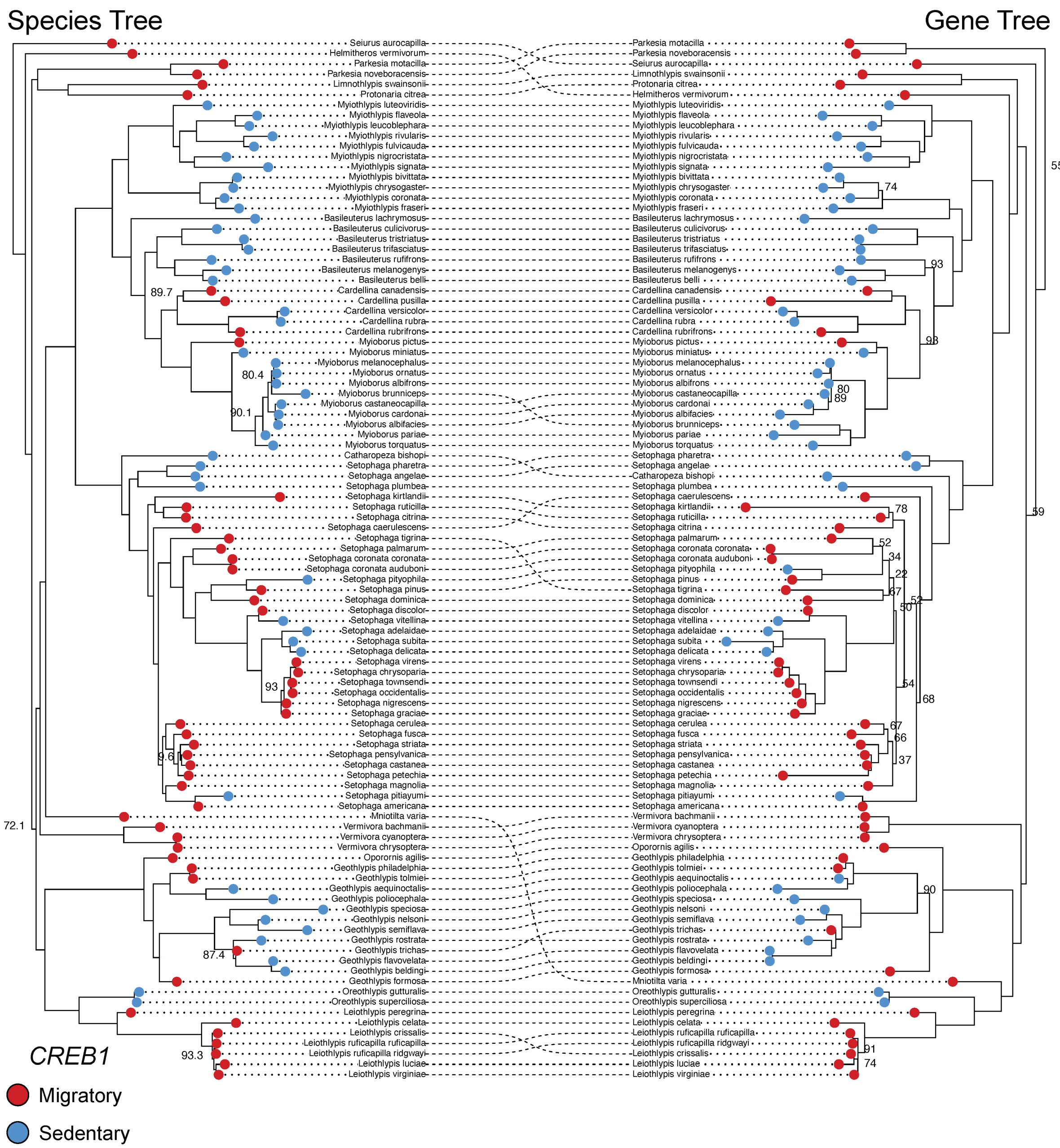

**FIGURE S6. Cophylogeny visualizing regions of concordance and discordance between the topology of a candidate migration gene tree (*CREB1*; right) and the parulid warbler species tree (left).** Dotted lines connect species between the two trees: parallel lines indicate identical placement in both trees, while curved intersecting lines indicate regions of discordance. Red circles at the tips of each tree represent migratory species and blue circles represent sedentary species. Bootstrap values <95 are shown at nodes.

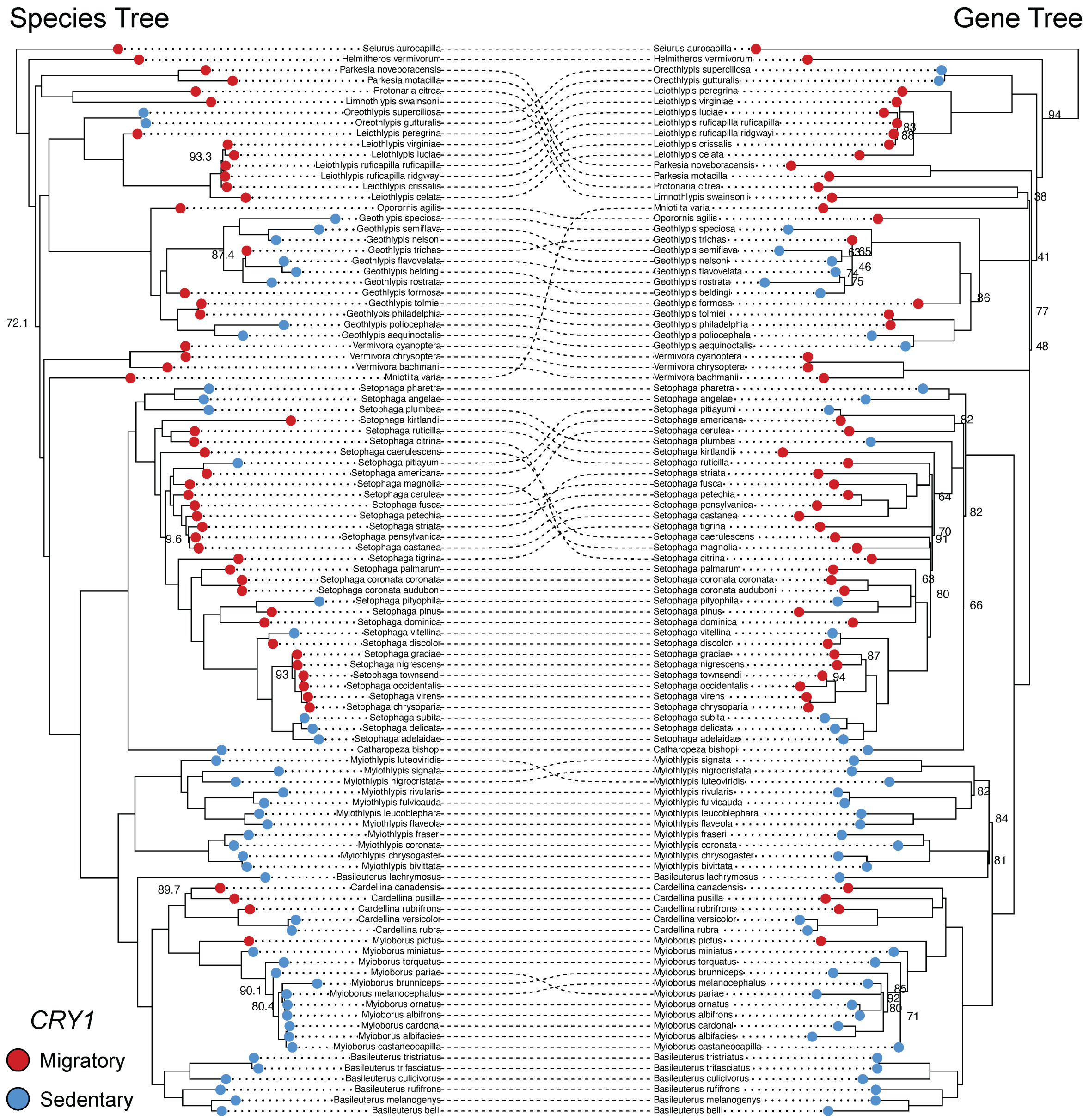

**FIGURE S7. Cophylogeny visualizing regions of concordance and discordance between the topology of a candidate migration gene tree (*CRY1*; right) and the parulid warbler species tree (left).** Dotted lines connect species between the two trees: parallel lines indicate identical placement in both trees, while curved intersecting lines indicate regions of discordance. Red circles at the tips of each tree represent migratory species and blue circles represent sedentary species. Bootstrap values <95 are shown at nodes.

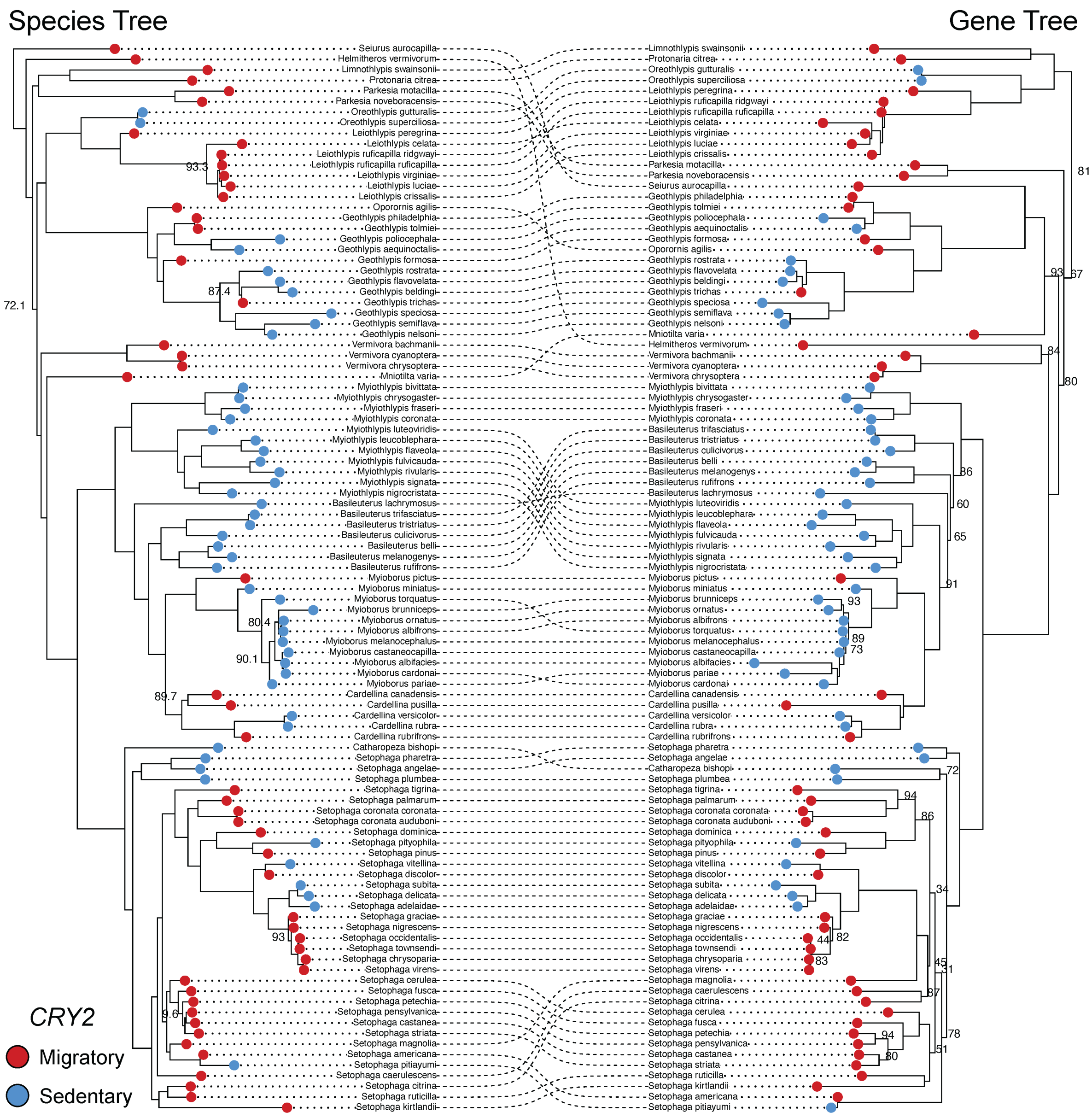

**FIGURE S8. Cophylogeny visualizing regions of concordance and discordance between the topology of a candidate migration gene tree (*CRY2*; right) and the parulid warbler species tree (left).** Dotted lines connect species between the two trees: parallel lines indicate identical placement in both trees, while curved intersecting lines indicate regions of discordance. Red circles at the tips of each tree represent migratory species and blue circles represent sedentary species. Bootstrap values <95 are shown at nodes.

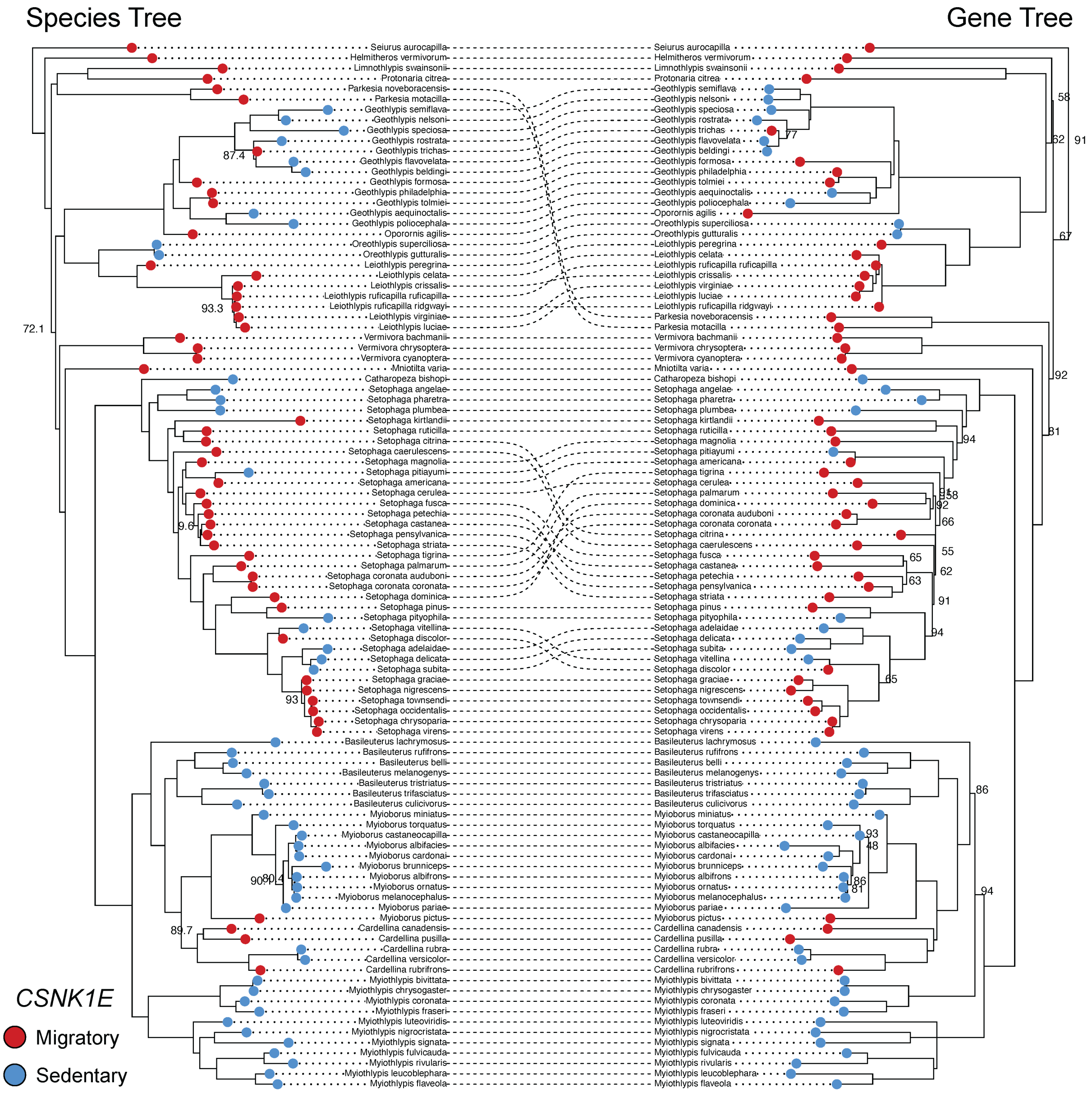

**FIGURE S9. Cophylogeny visualizing regions of concordance and discordance between the topology of a candidate migration gene tree (*CSNK1E*; right) and the parulid warbler species tree (left).** Dotted lines connect species between the two trees: parallel lines indicate identical placement in both trees, while curved intersecting lines indicate regions of discordance. Red circles at the tips of each tree represent migratory species and blue circles represent sedentary species. Bootstrap values <95 are shown at nodes.

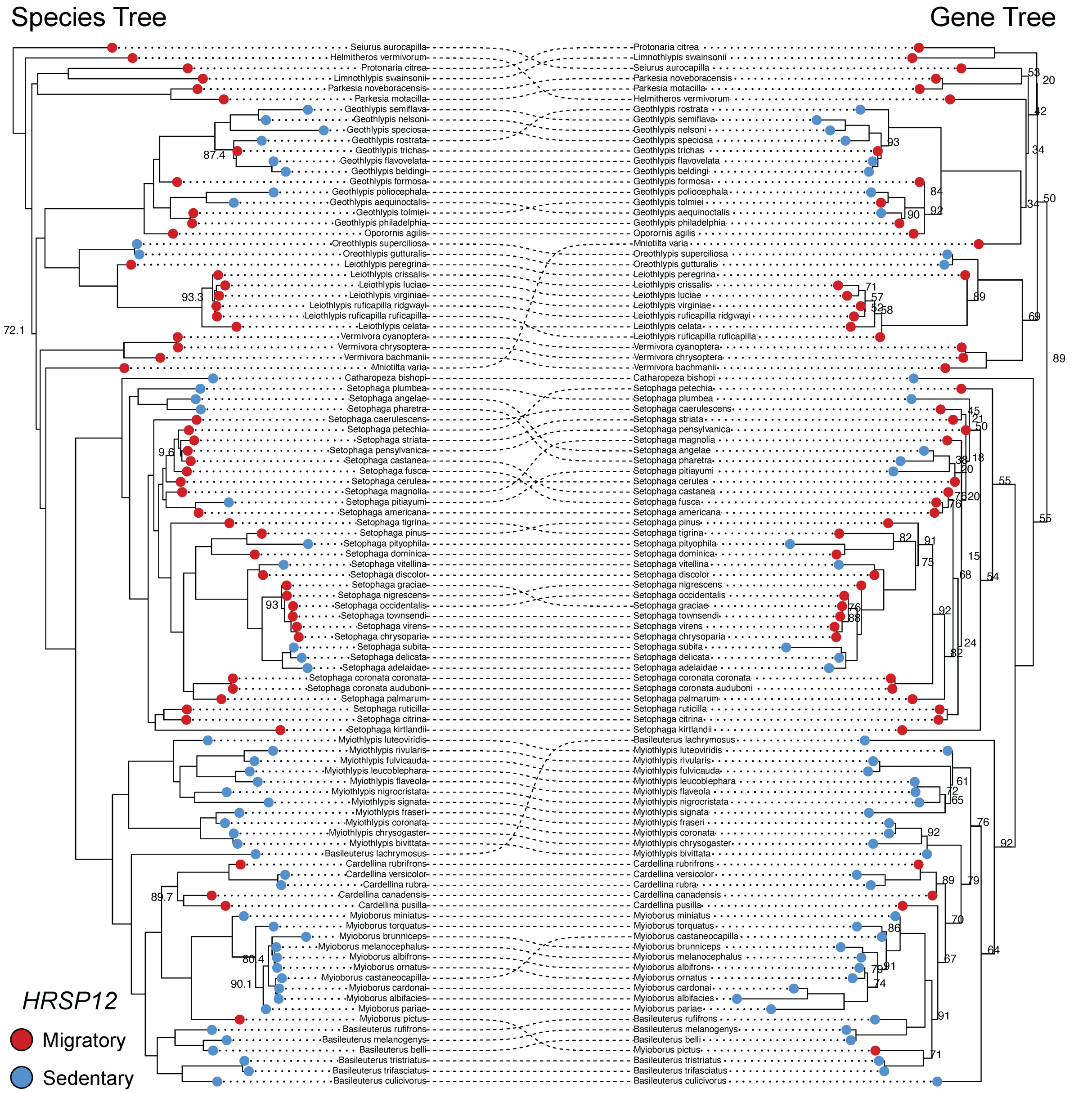

**FIGURE S10. Cophylogeny visualizing regions of concordance and discordance between the topology of a candidate migration gene tree (*HRSP12*; right) and the parulid warbler species tree (left).** Dotted lines connect species between the two trees: parallel lines indicate identical placement in both trees, while curved intersecting lines indicate regions of discordance. Red circles at the tips of each tree represent migratory species and blue circles represent sedentary species. Bootstrap values <95 are shown at nodes.

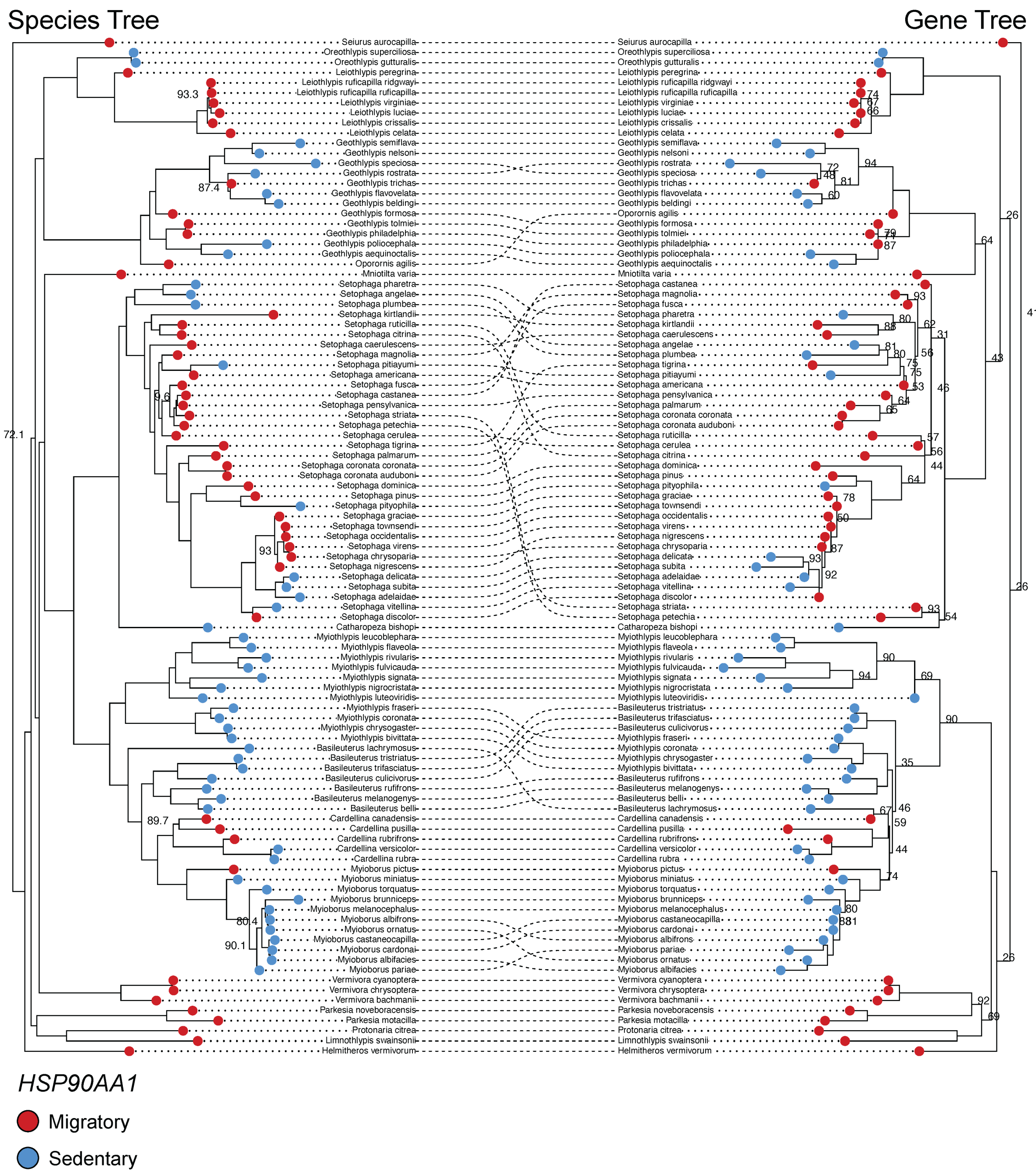

**FIGURE S11. Cophylogeny visualizing regions of concordance and discordance between the topology of a candidate migration gene tree (*HSP90AA1*; right) and the parulid warbler species tree (left).** Dotted lines connect species between the two trees: parallel lines indicate identical placement in both trees, while curved intersecting lines indicate regions of discordance. Red circles at the tips of each tree represent migratory species and blue circles represent sedentary species. Bootstrap values <95 are shown at nodes.

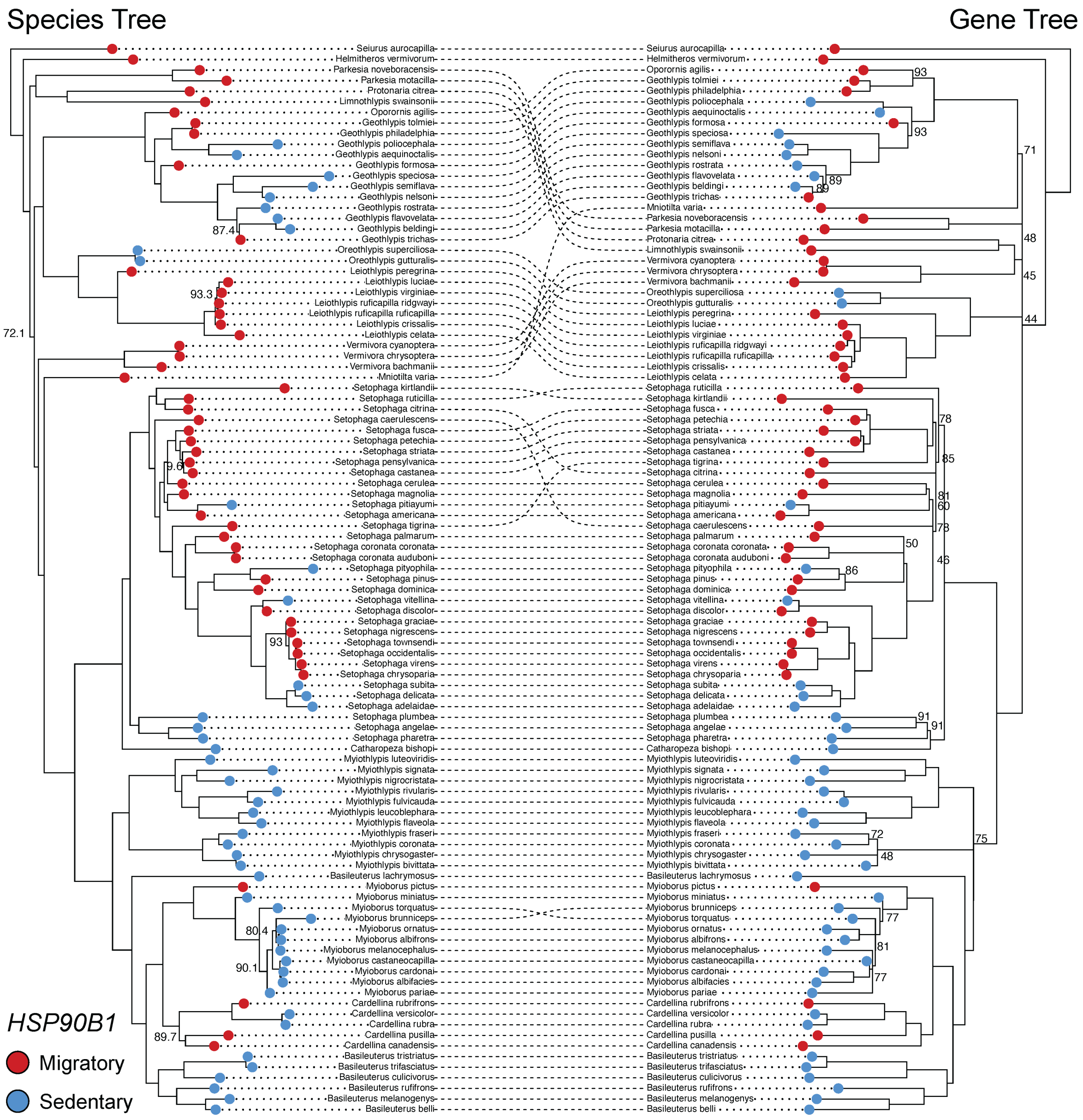

**FIGURE S12. Cophylogeny visualizing regions of concordance and discordance between the topology of a candidate migration gene tree (*HSP90B1*; right) and the parulid warbler species tree (left).** Dotted lines connect species between the two trees: parallel lines indicate identical placement in both trees, while curved intersecting lines indicate regions of discordance. Red circles at the tips of each tree represent migratory species and blue circles represent sedentary species. Bootstrap values <95 are shown at nodes.

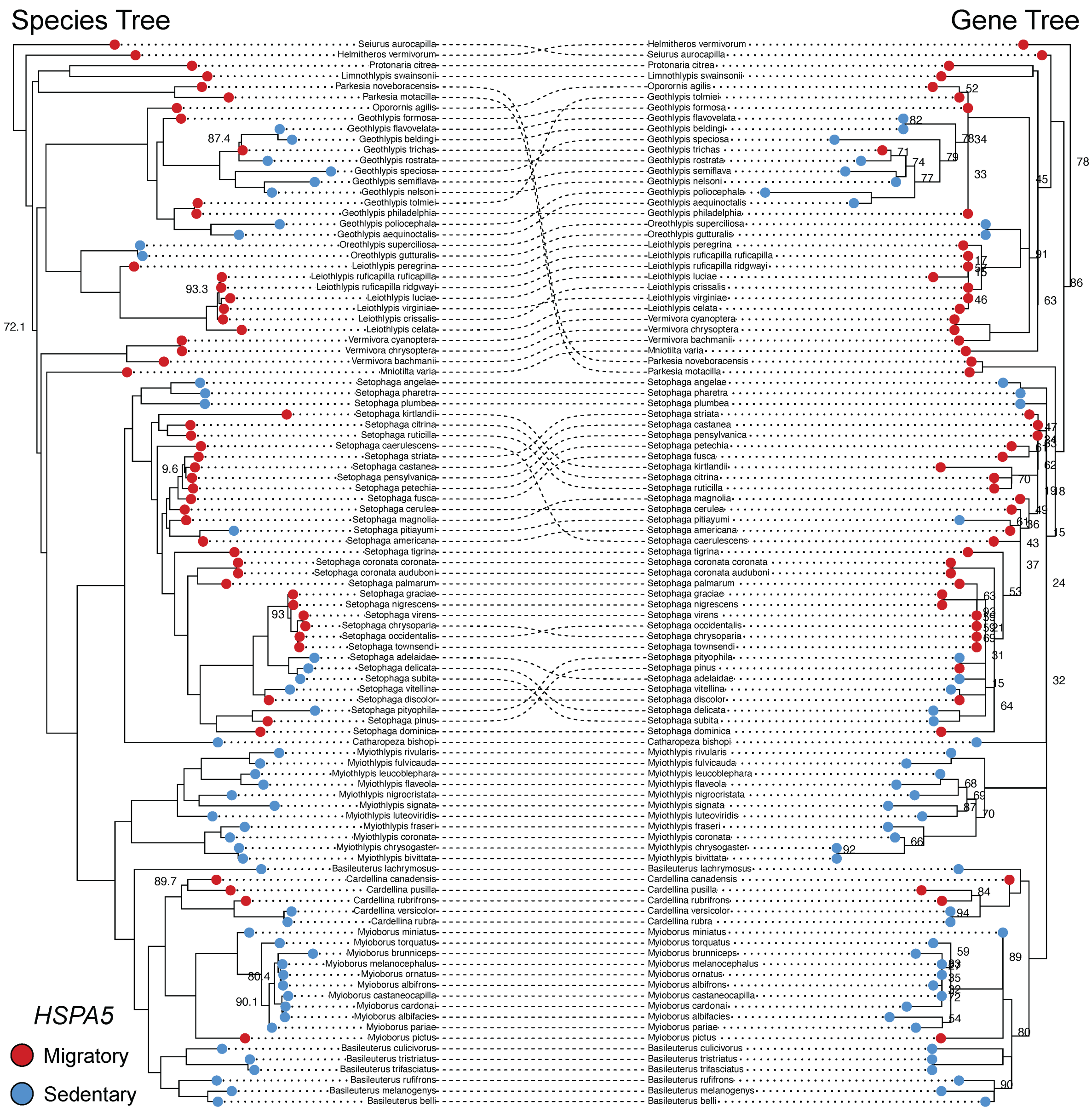

**FIGURE S13. Cophylogeny visualizing regions of concordance and discordance between the topology of a candidate migration gene tree (*HSPA5*; right) and the parulid warbler species tree (left).** Dotted lines connect species between the two trees: parallel lines indicate identical placement in both trees, while curved intersecting lines indicate regions of discordance. Red circles at the tips of each tree represent migratory species and blue circles represent sedentary species. Bootstrap values <95 are shown at nodes.

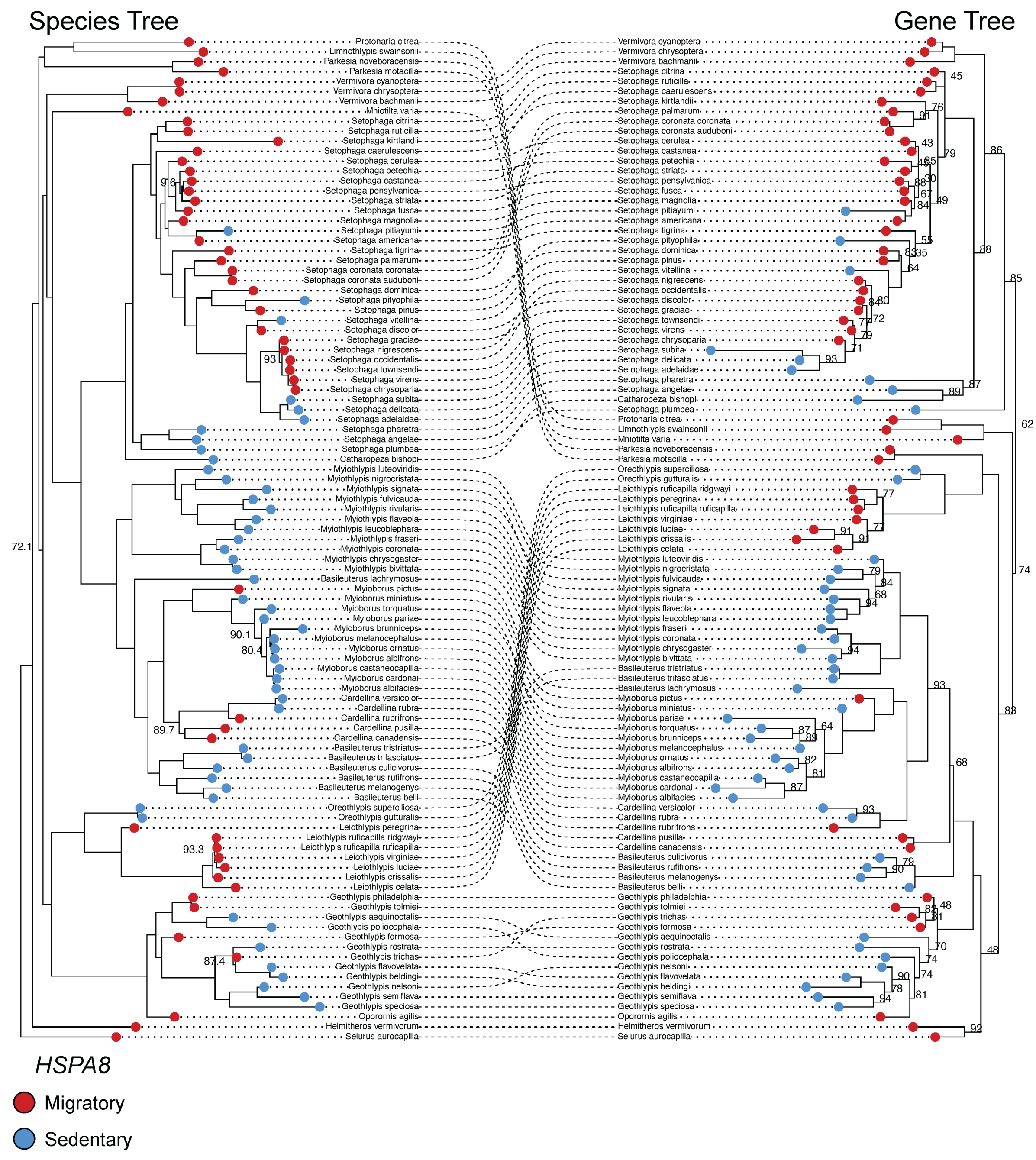

**FIGURE S14. Cophylogeny visualizing regions of concordance and discordance between the topology of a candidate migration gene tree (*HSPA8*; right) and the parulid warbler species tree (left).** Dotted lines connect species between the two trees: parallel lines indicate identical placement in both trees, while curved intersecting lines indicate regions of discordance. Red circles at the tips of each tree represent migratory species and blue circles represent sedentary species. Bootstrap values <95 are shown at nodes.

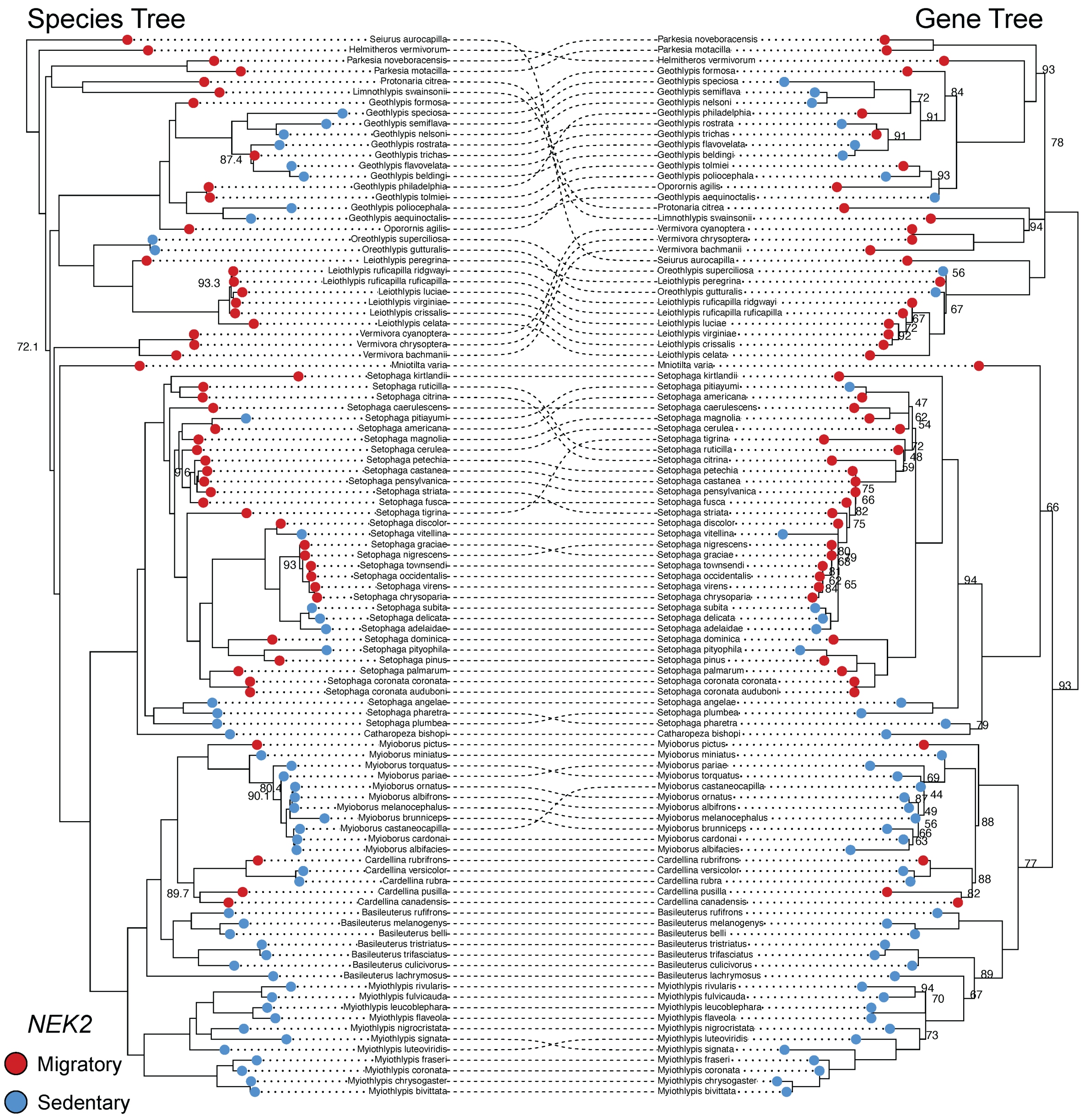

**FIGURE S15. Cophylogeny visualizing regions of concordance and discordance between the topology of a candidate migration gene tree (*NEK2*; right) and the parulid warbler species tree (left).** Dotted lines connect species between the two trees: parallel lines indicate identical placement in both trees, while curved intersecting lines indicate regions of discordance. Red circles at the tips of each tree represent migratory species and blue circles represent sedentary species. Bootstrap values <95 are shown at nodes.

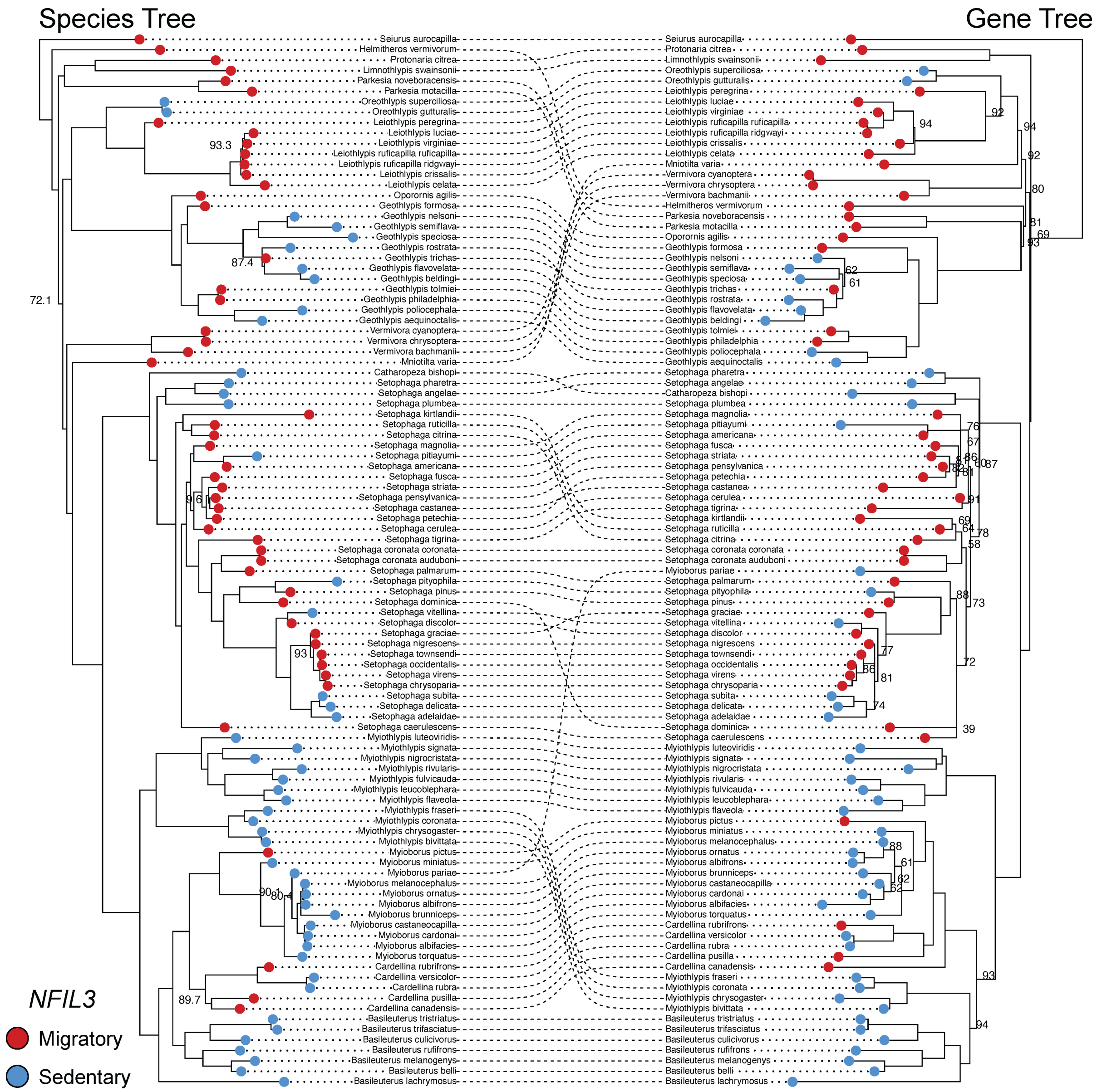

**FIGURE S16. Cophylogeny visualizing regions of concordance and discordance between the topology of a candidate migration gene tree (*NFIL3*; right) and the parulid warbler species tree (left).** Dotted lines connect species between the two trees: parallel lines indicate identical placement in both trees, while curved intersecting lines indicate regions of discordance. Red circles at the tips of each tree represent migratory species and blue circles represent sedentary species. Bootstrap values <95 are shown at nodes.

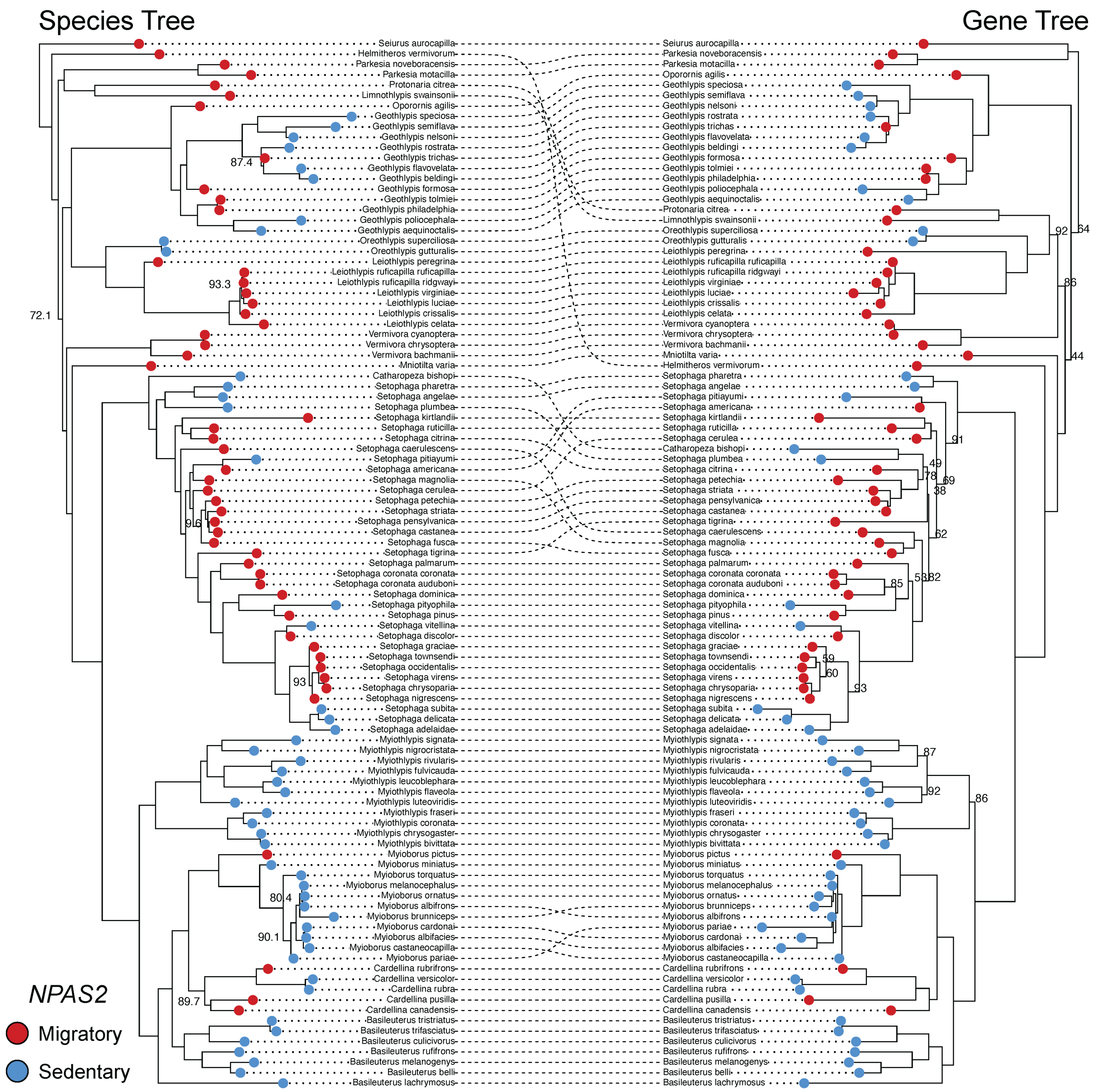

**FIGURE S17. Cophylogeny visualizing regions of concordance and discordance between the topology of a candidate migration gene tree (*NPAS2*; right) and the parulid warbler species tree (left).** Dotted lines connect species between the two trees: parallel lines indicate identical placement in both trees, while curved intersecting lines indicate regions of discordance. Red circles at the tips of each tree represent migratory species and blue circles represent sedentary species. Bootstrap values <95 are shown at nodes.

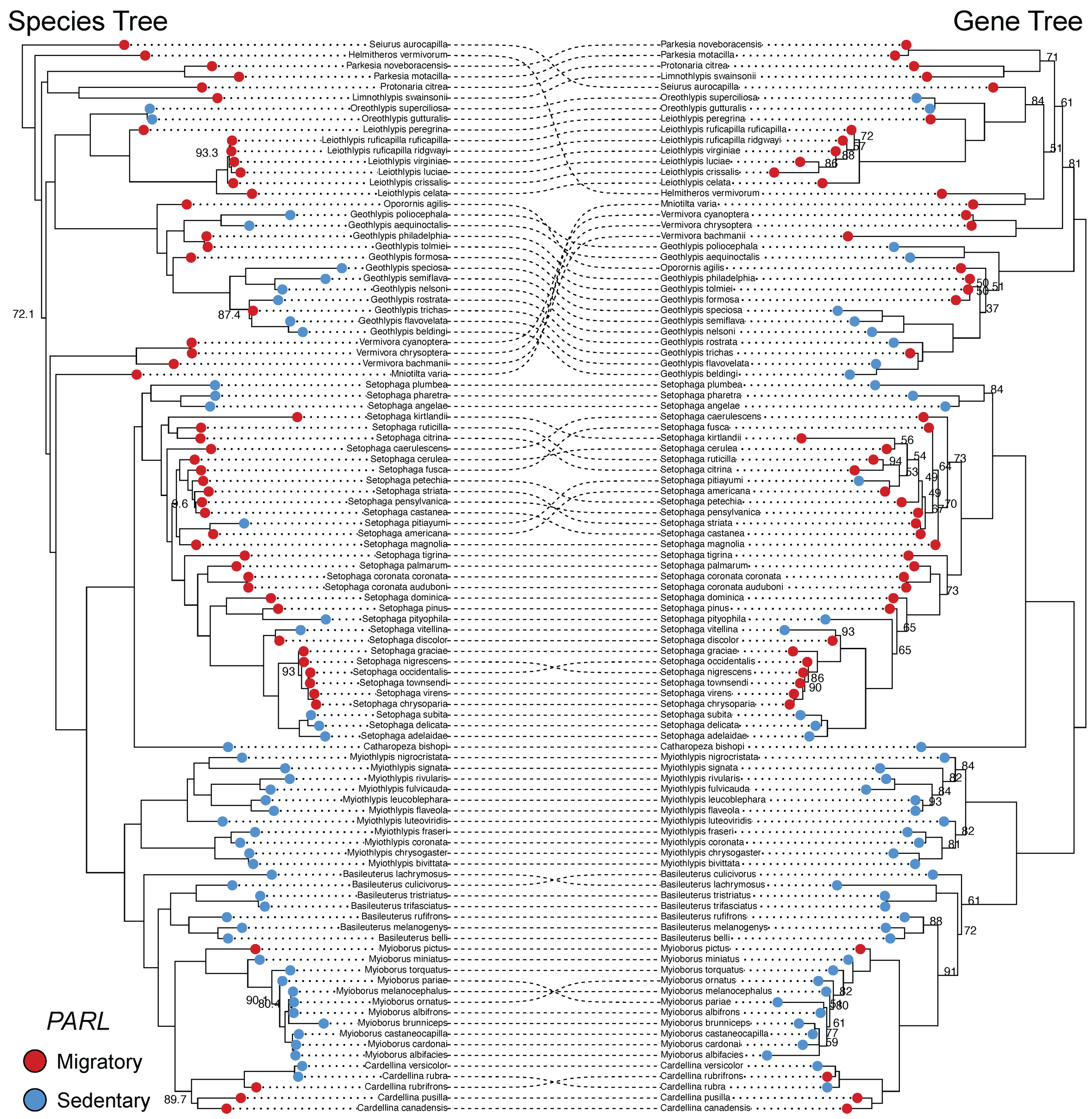

**FIGURE S18. Cophylogeny visualizing regions of concordance and discordance between the topology of a candidate migration gene tree (*PARL*; right) and the parulid warbler species tree (left).** Dotted lines connect species between the two trees: parallel lines indicate identical placement in both trees, while curved intersecting lines indicate regions of discordance. Red circles at the tips of each tree represent migratory species and blue circles represent sedentary species. Bootstrap values <95 are shown at nodes.

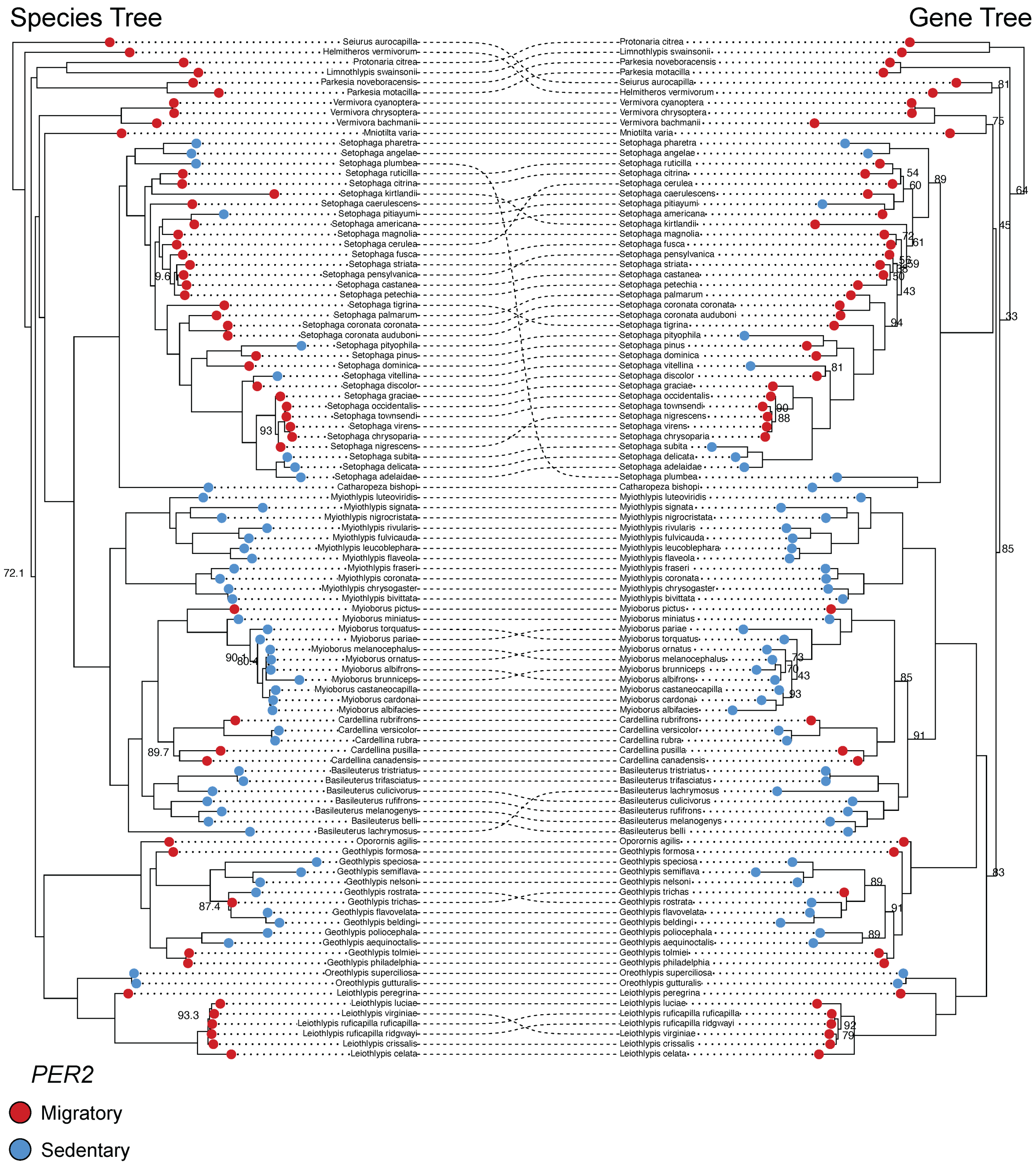

**FIGURE S19. Cophylogeny visualizing regions of concordance and discordance between the topology of a candidate migration gene tree (*PER2*; right) and the parulid warbler species tree (left).** Dotted lines connect species between the two trees: parallel lines indicate identical placement in both trees, while curved intersecting lines indicate regions of discordance. Red circles at the tips of each tree represent migratory species and blue circles represent sedentary species. Bootstrap values <95 are shown at nodes.

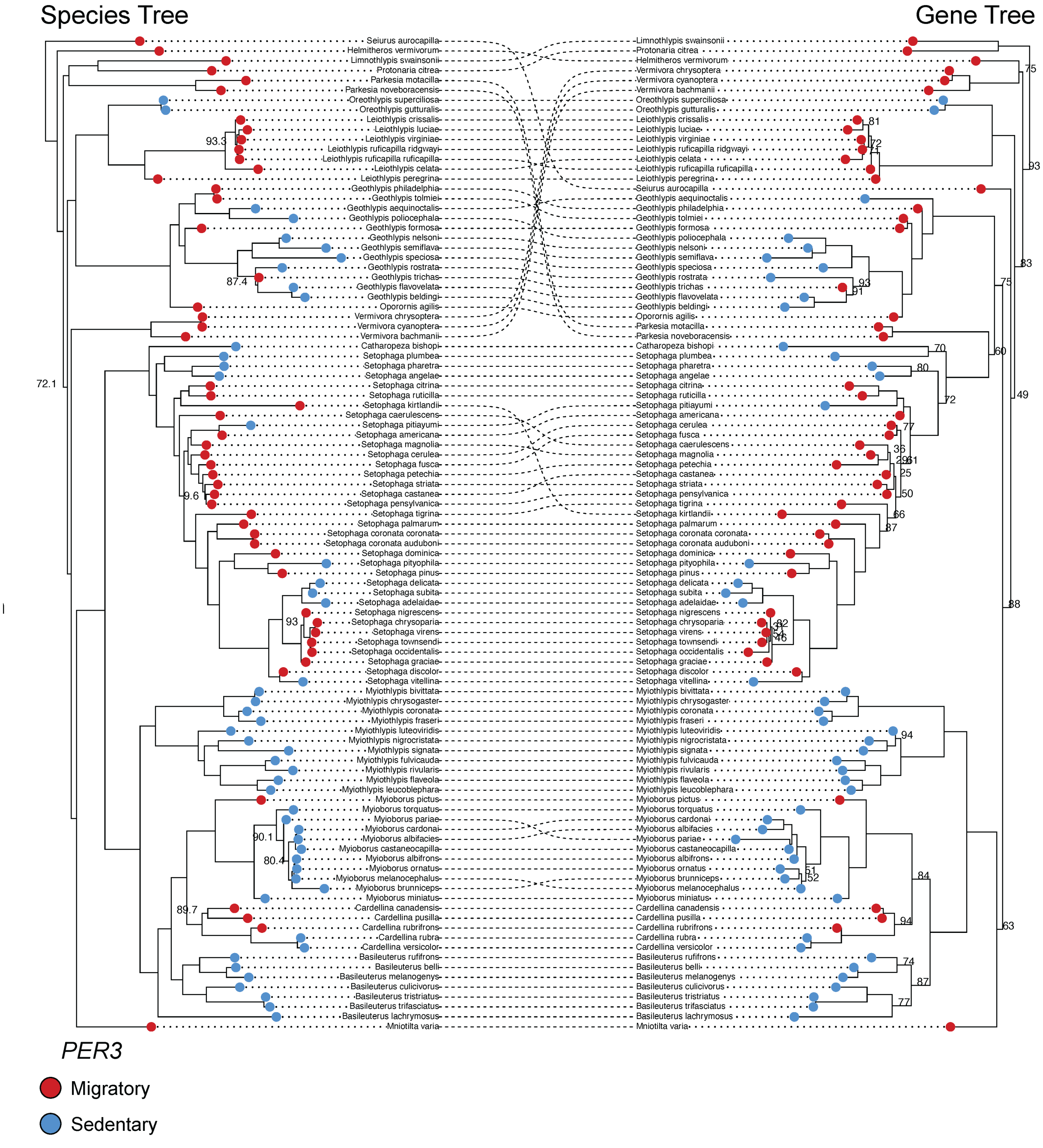

**FIGURE S20. Cophylogeny visualizing regions of concordance and discordance between the topology of a candidate migration gene tree (*PER3*; right) and the parulid warbler species tree (left).** Dotted lines connect species between the two trees: parallel lines indicate identical placement in both trees, while curved intersecting lines indicate regions of discordance. Red circles at the tips of each tree represent migratory species and blue circles represent sedentary species. Bootstrap values <95 are shown at nodes.

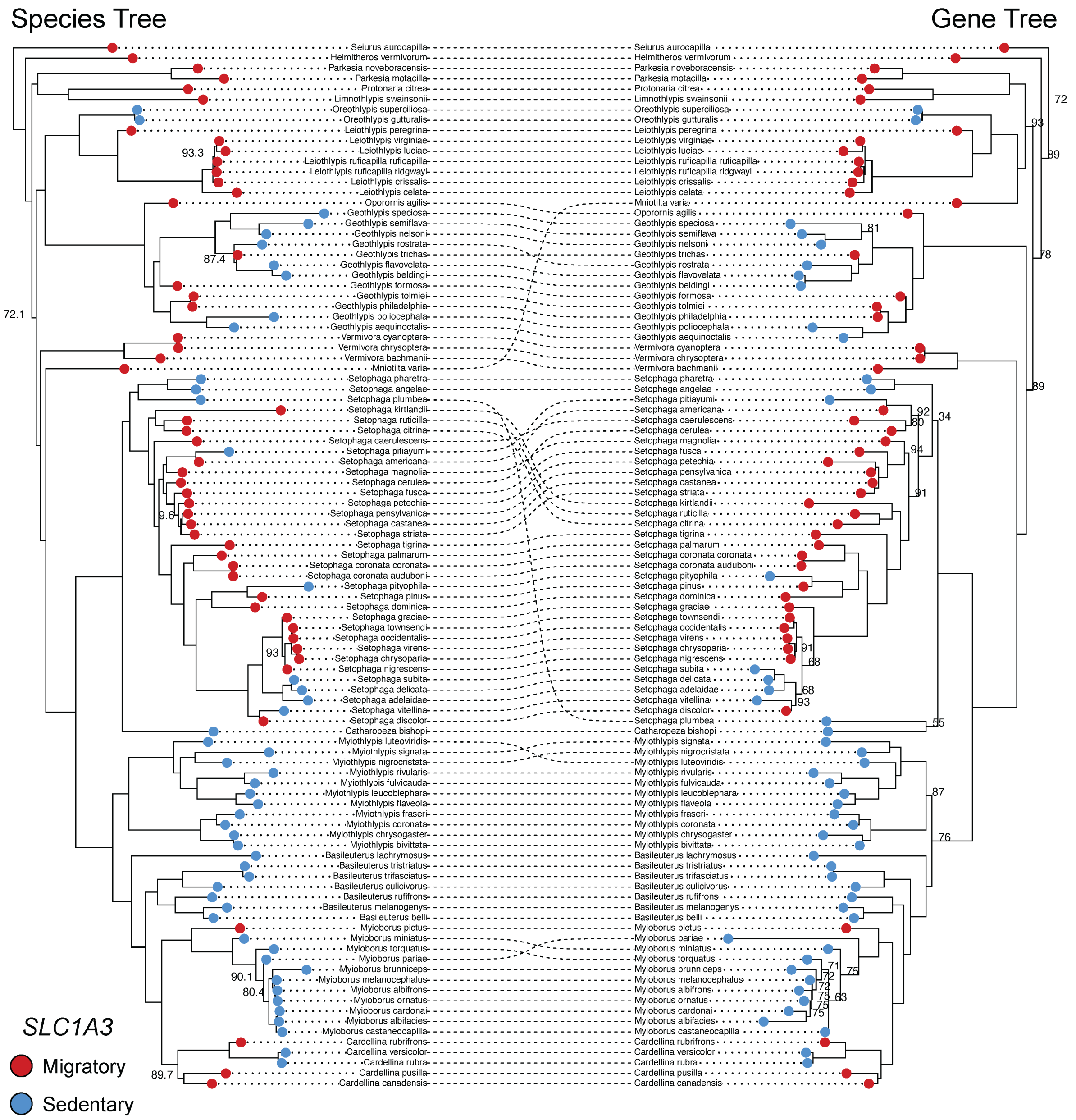

**FIGURE S21. Cophylogeny visualizing regions of concordance and discordance between the topology of a candidate migration gene tree (*SLC1A3*; right) and the parulid warbler species tree (left).** Dotted lines connect species between the two trees: parallel lines indicate identical placement in both trees, while curved intersecting lines indicate regions of discordance. Red circles at the tips of each tree represent migratory species and blue circles represent sedentary species. Bootstrap values <95 are shown at nodes.

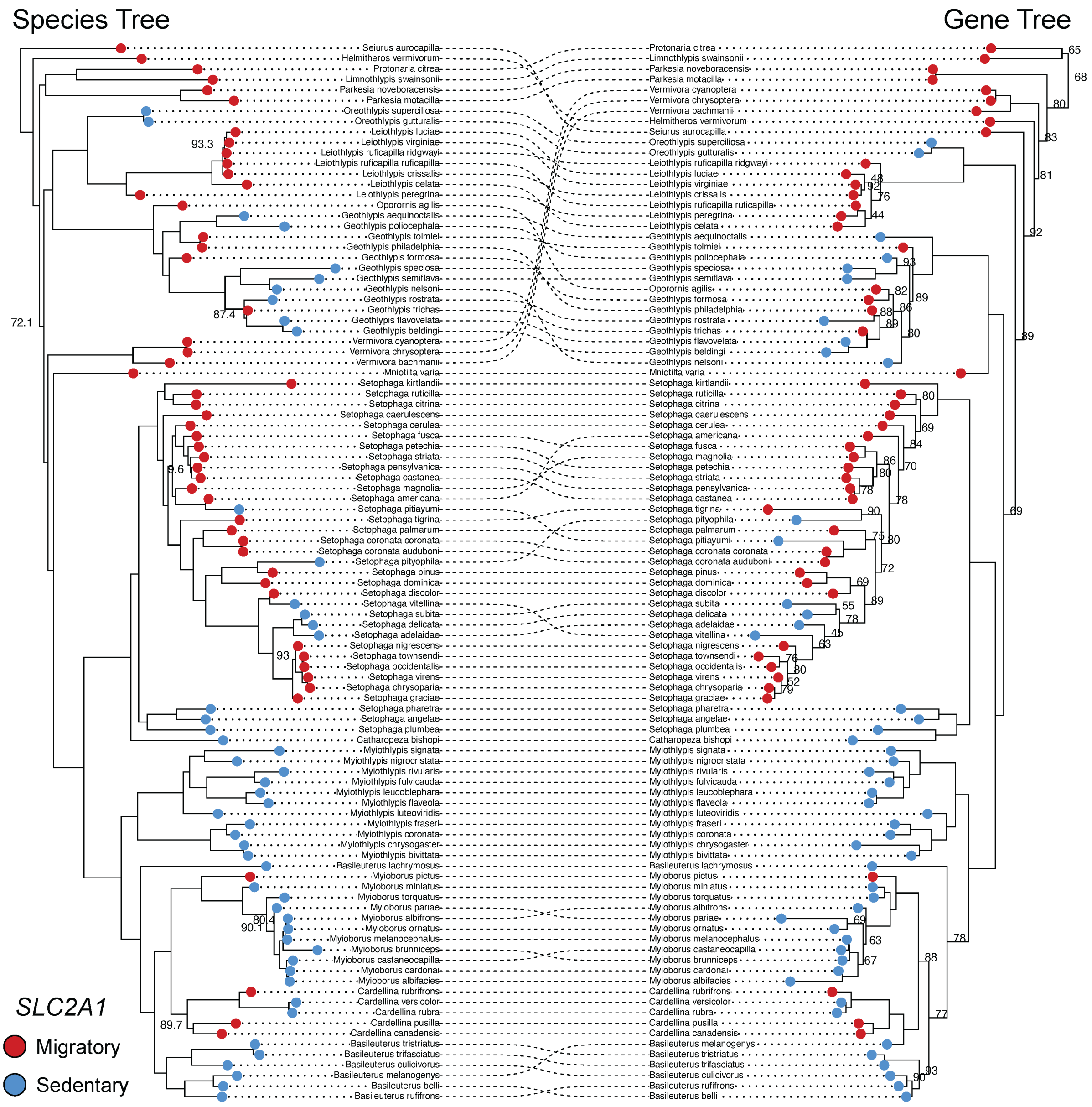

**FIGURE S22. Cophylogeny visualizing regions of concordance and discordance between the topology of a candidate migration gene tree (*SLC2A1*; right) and the parulid warbler species tree (left).** Dotted lines connect species between the two trees: parallel lines indicate identical placement in both trees, while curved intersecting lines indicate regions of discordance. Red circles at the tips of each tree represent migratory species and blue circles represent sedentary species. Bootstrap values <95 are shown at nodes.

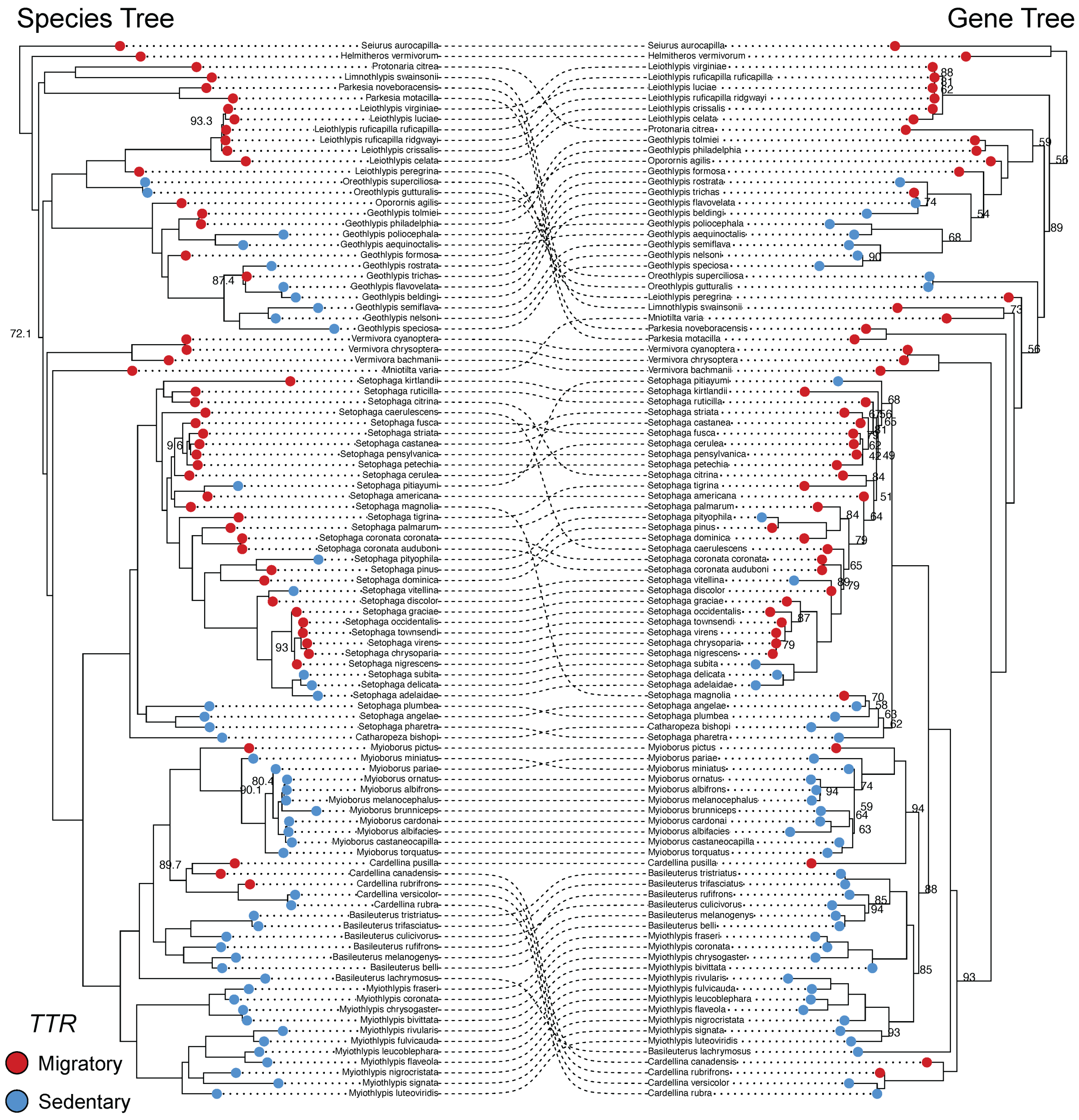

**FIGURE S23. Cophylogeny visualizing regions of concordance and discordance between the topology of a candidate migration gene tree (*TTR*; right) and the parulid warbler species tree (left).** Dotted lines connect species between the two trees: parallel lines indicate identical placement in both trees, while curved intersecting lines indicate regions of discordance. Red circles at the tips of each tree represent migratory species and blue circles represent sedentary species. Bootstrap values <95 are shown at nodes.

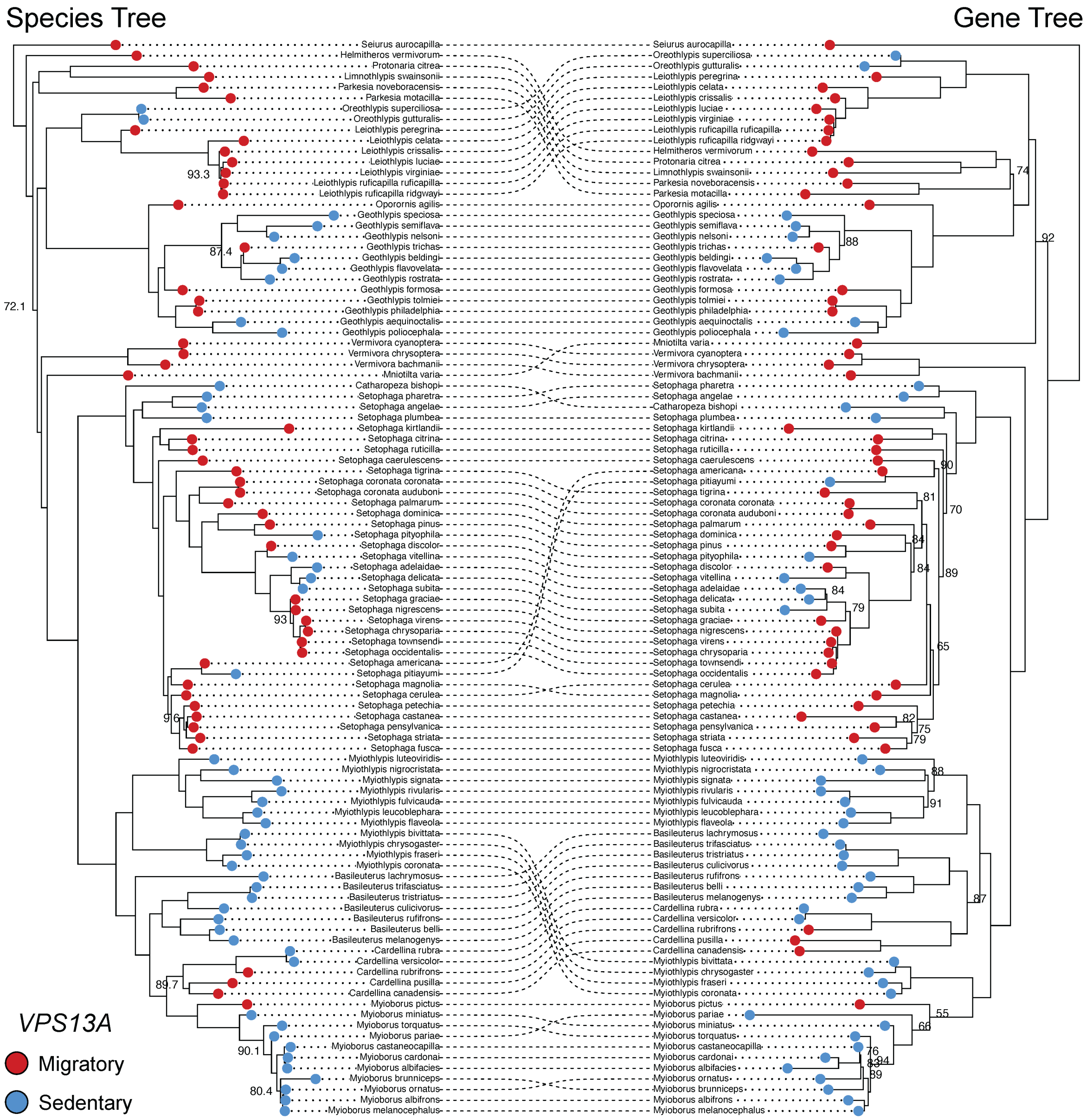

**FIGURE S24. Cophylogeny visualizing regions of concordance and discordance between the topology of a candidate migration gene tree (*VPS13A*; right) and the parulid warbler species tree (left).** Dotted lines connect species between the two trees: parallel lines indicate identical placement in both trees, while curved intersecting lines indicate regions of discordance. Red circles at the tips of each tree represent migratory species and blue circles represent sedentary species. Bootstrap values <95 are shown at nodes.

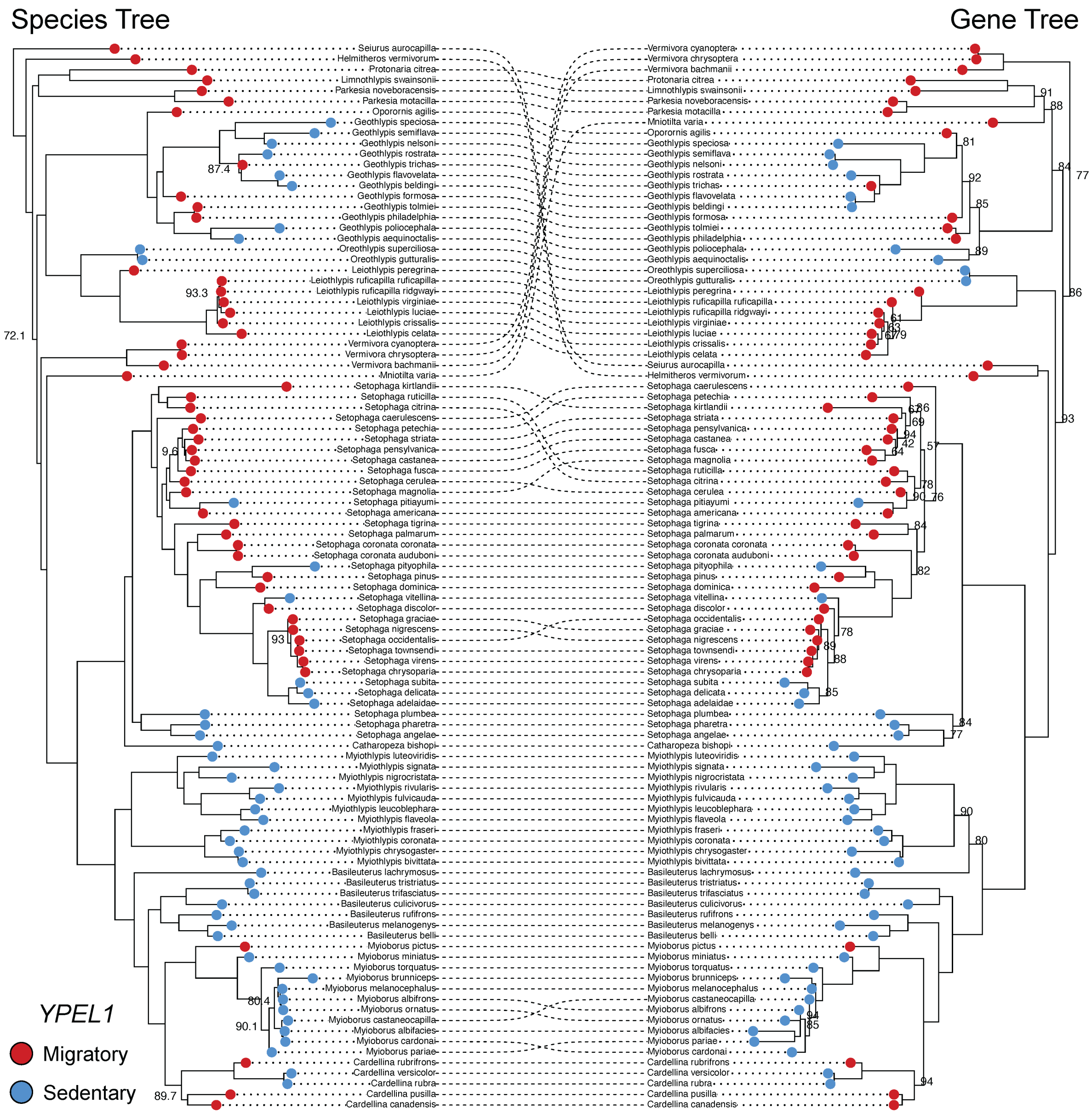

**FIGURE S25. Cophylogeny visualizing regions of concordance and discordance between the topology of a candidate migration gene tree (*YPEL1*; right) and the parulid warbler species tree (left).** Dotted lines connect species between the two trees: parallel lines indicate identical placement in both trees, while curved intersecting lines indicate regions of discordance. Red circles at the tips of each tree represent migratory species and blue circles represent sedentary species. Bootstrap values <95 are shown at nodes.

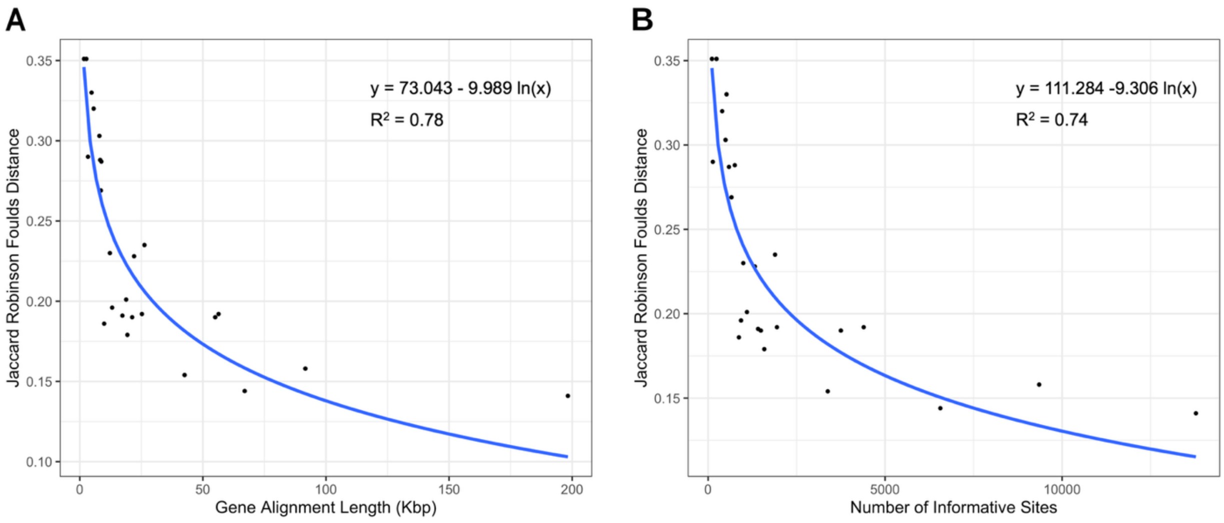

**FIGURE S26. Relationship between normalized Jaccard Robinson-Foulds (JRF) distance and (A) alignment length or (B) number of parsimony informative sites for 25 candidate migration genes.** The logarithmic best fit line is plotted in blue for each comparison. JRF distance was significantly related to both alignment length ($P = 4.6e^{-9}$) and the number of informative sites ($P = 1.8e^{-8}$).

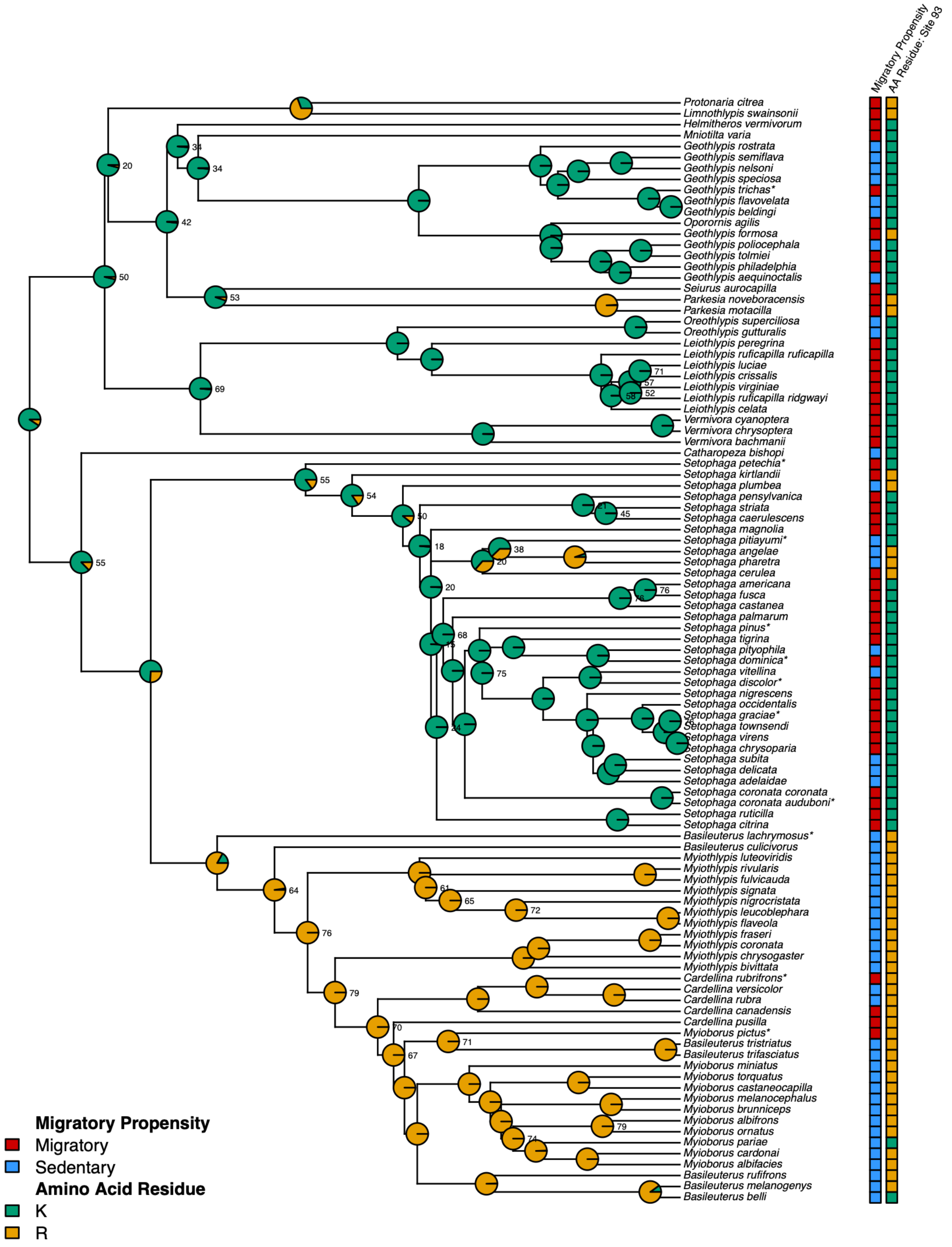

**FIGURE S27. Amino acid variation at *HRSP12* site 93 mapped across the *HRSP12* gene tree for all parulid warblers.** Square labels at tips represent migratory propensity (red: migratory, blue: sedentary) and amino acid residue (orange: arginine, green: lysine) for each species. Pie charts on nodes show ancestral character state reconstruction posterior probabilities for each amino acid variant. Nodes of the tree that received <80% support by approximate likelihood ratio test (aLRT) are labeled with their aLRT value. Partially migratory species are indicated by asterisks next to their species name.

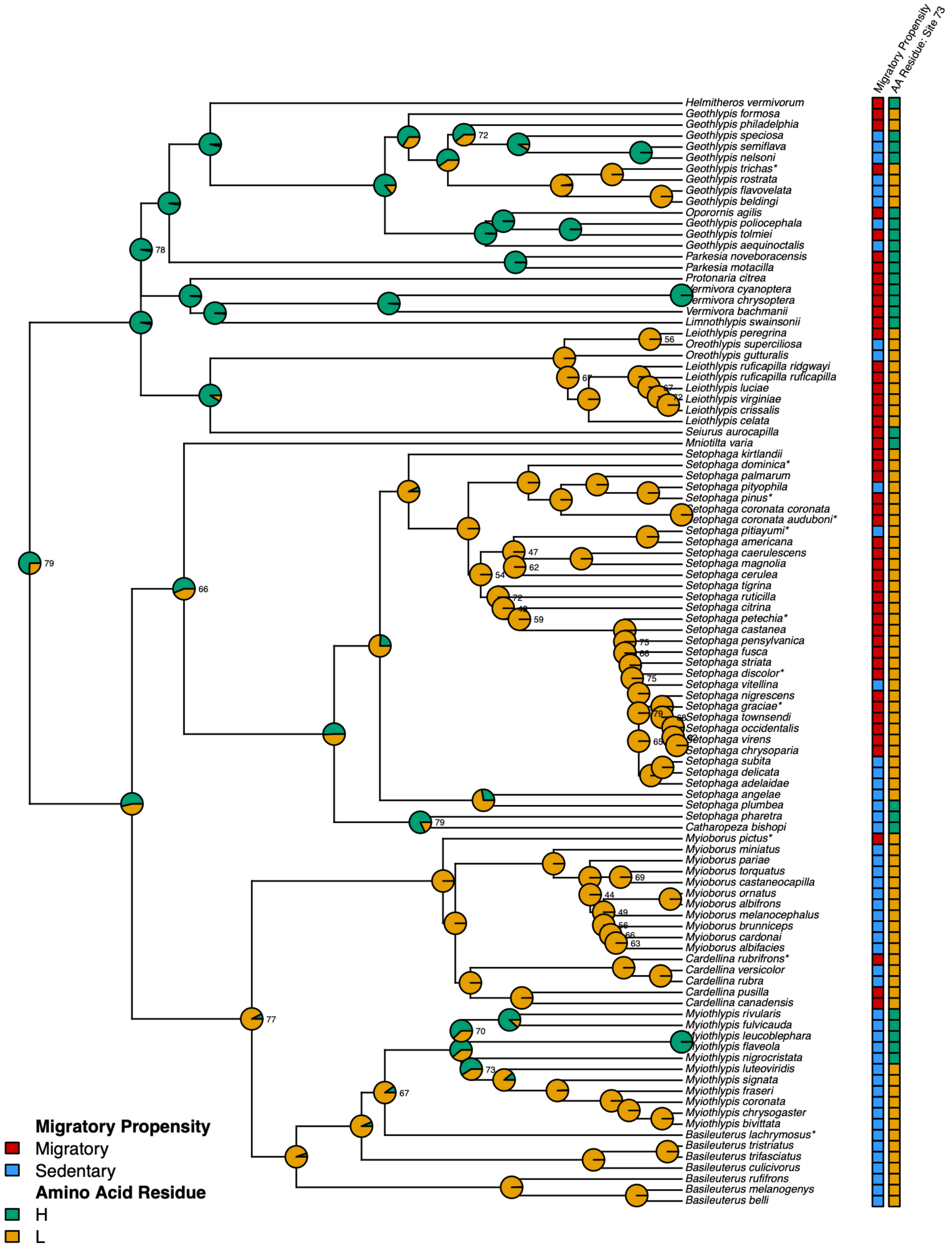

**FIGURE S28. Amino acid variation at *NEK2* site 73 mapped across the *NEK2* gene tree for all parulid warblers.** Square labels at tips represent migratory propensity (red: migratory, blue: sedentary) and amino acid residue (orange: leucine, green: histidine) for each species. Pie charts on nodes show ancestral character state reconstruction posterior probabilities for each amino acid variant. Nodes of the tree that received <80% support by approximate likelihood ratio test (aLRT) are labeled with their aLRT value. Partially migratory species are indicated by asterisks next to their species name.

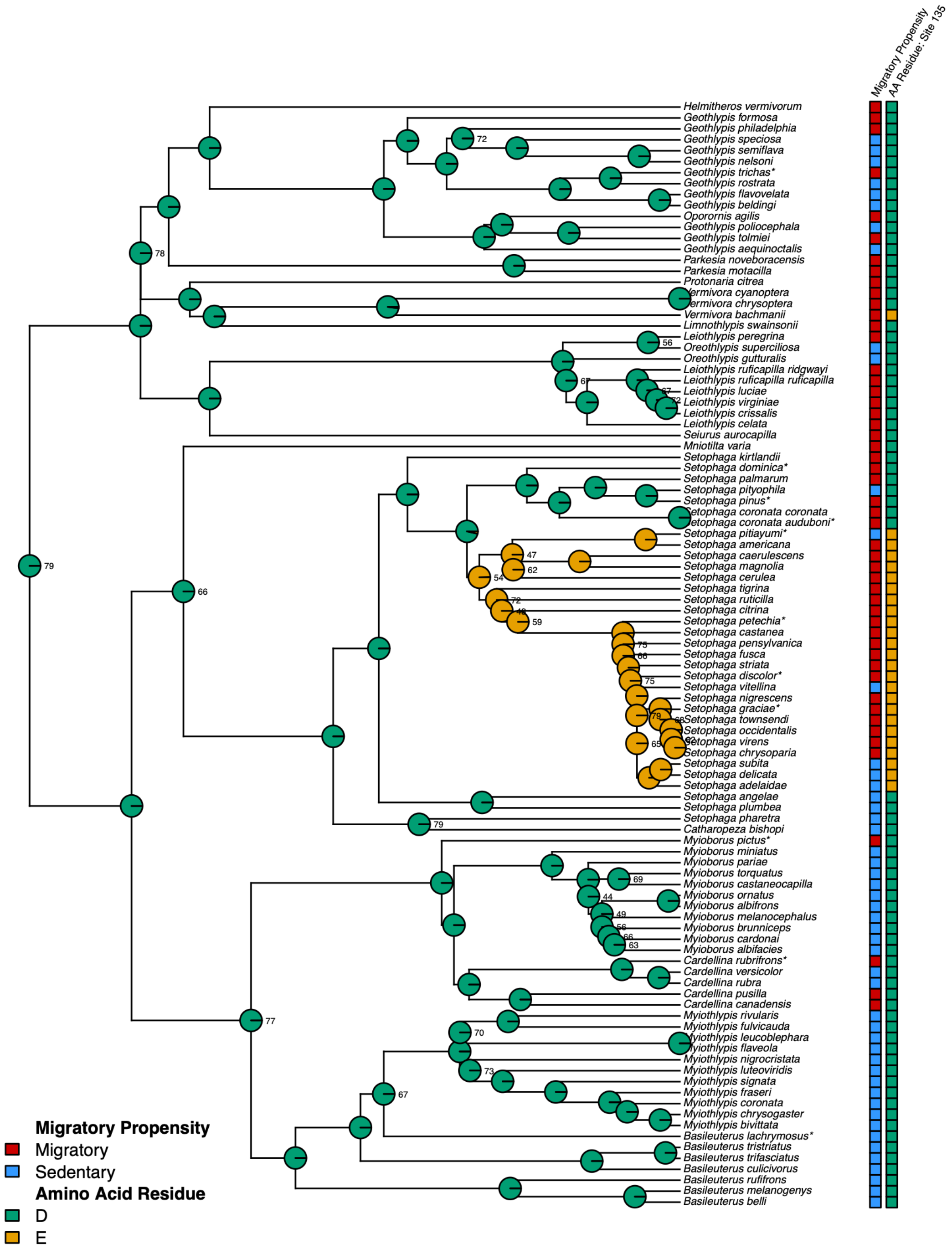

**FIGURE S29. Amino acid variation at *NEK2* site 135 mapped across the *NEK2* gene tree for all parulid warblers.** Square labels at tips represent migratory propensity (red: migratory, blue: sedentary) and amino acid residue (orange: glutamic acid, green: aspartic acid) for each species. Pie charts on nodes show ancestral character state reconstruction posterior probabilities for each amino acid variant. Nodes of the tree that received <80% support by approximate likelihood ratio test (aLRT) are labeled with their aLRT value. Partially migratory species are indicated by asterisks next to their species name.

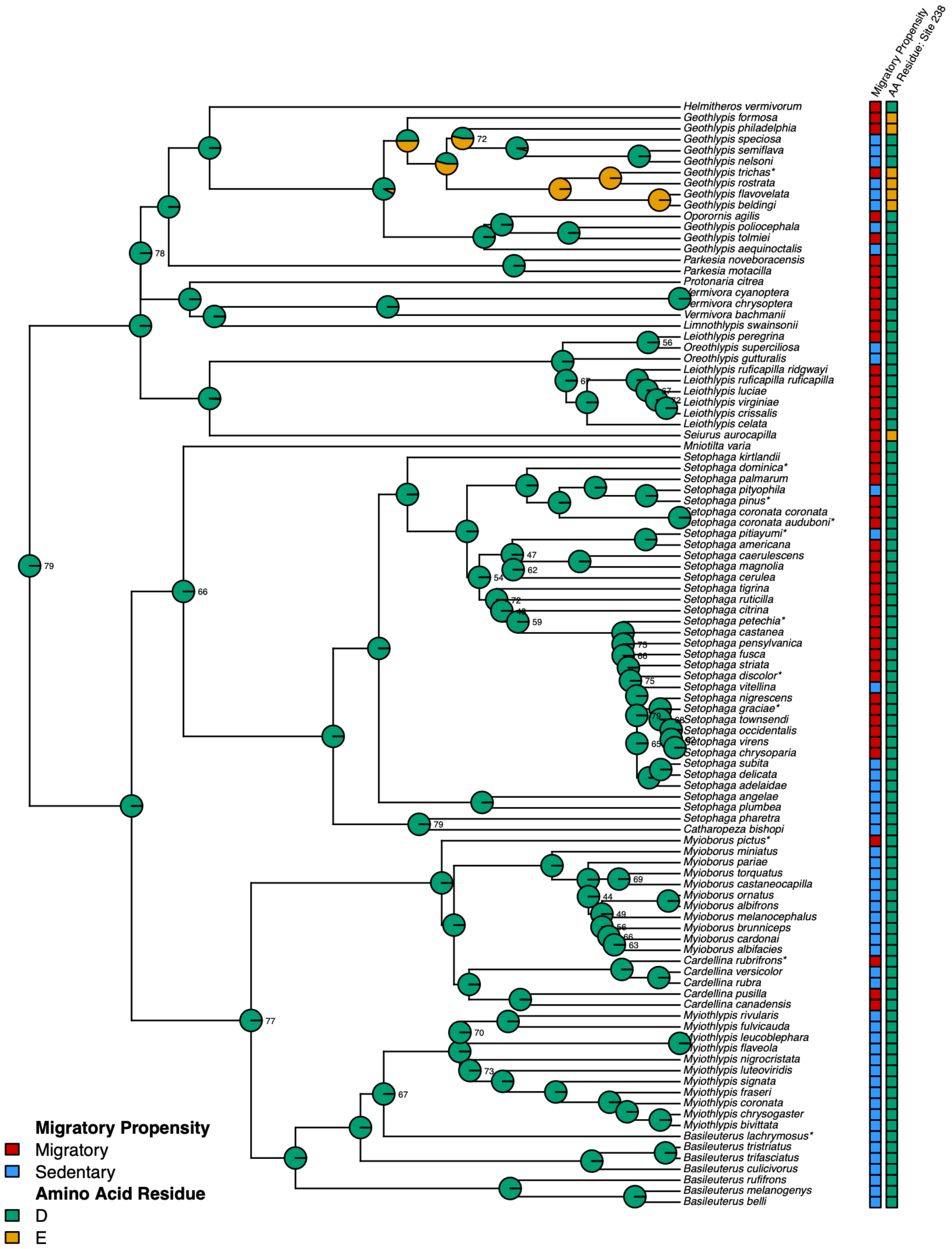

**FIGURE S30. Amino acid variation at *NEK2* site 238 mapped across the *NEK2* gene tree for all parulid warblers.** Square labels at tips represent migratory propensity (red: migratory, blue: sedentary) and amino acid residue (orange: glutamic acid, green: aspartic acid) for each species. Pie charts on nodes show ancestral character state reconstruction posterior probabilities for each amino acid variant. Nodes of the tree that received <80% support by approximate likelihood ratio test (aLRT) are labeled with their aLRT value. Partially migratory species are indicated by asterisks next to their species name.

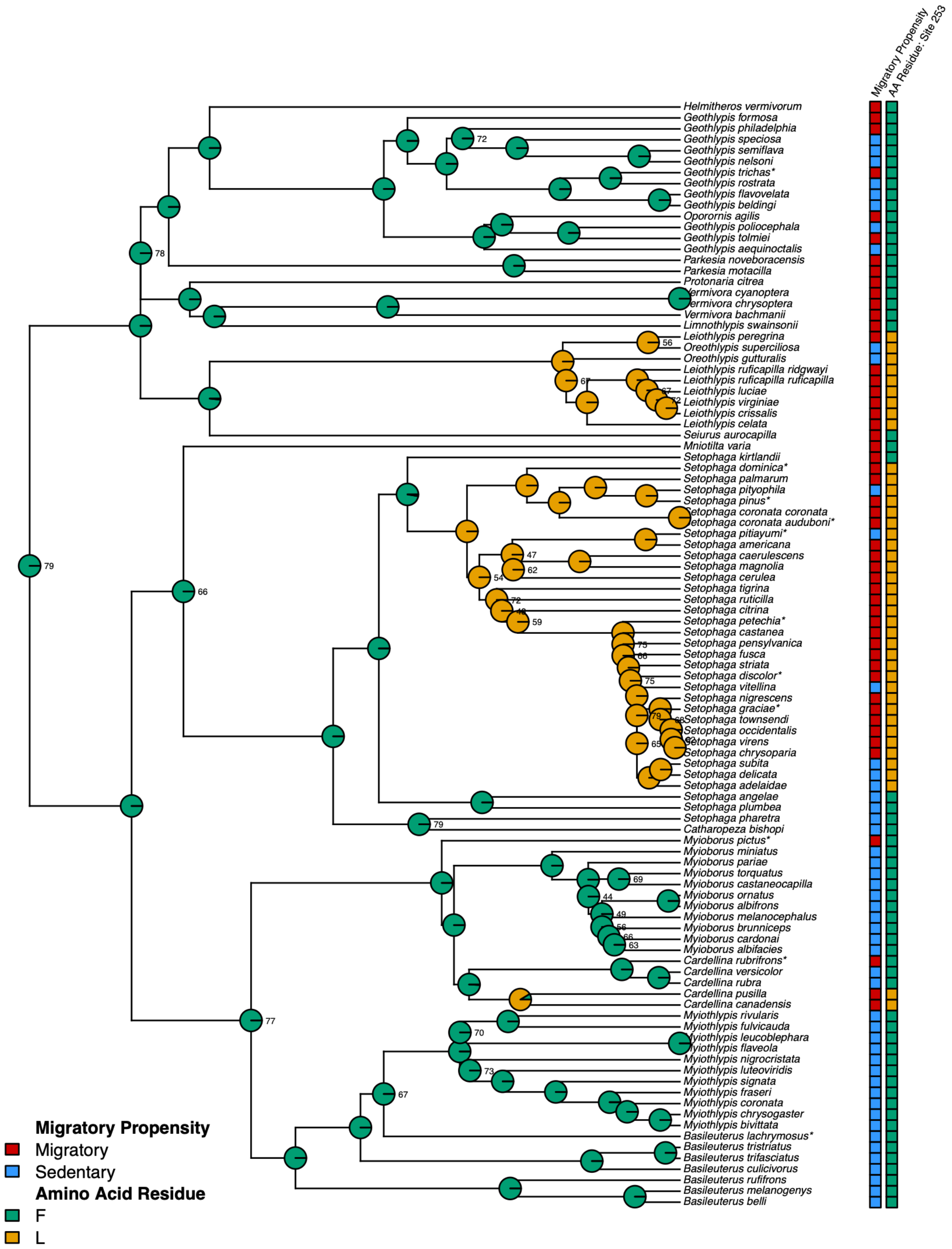

**FIGURE S31. Amino acid variation at *NEK2* site 253 mapped across the *NEK2* gene tree for all parulid warblers.** Square labels at tips represent migratory propensity (red: migratory, blue: sedentary) and amino acid residue (orange: leucine, green: phenylalanine) for each species. Pie charts on nodes show ancestral character state reconstruction posterior probabilities for each amino acid variant. Nodes of the tree that received <80% support by approximate likelihood ratio test (aLRT) are labeled with their aLRT value. Partially migratory species are indicated by asterisks next to their species name.

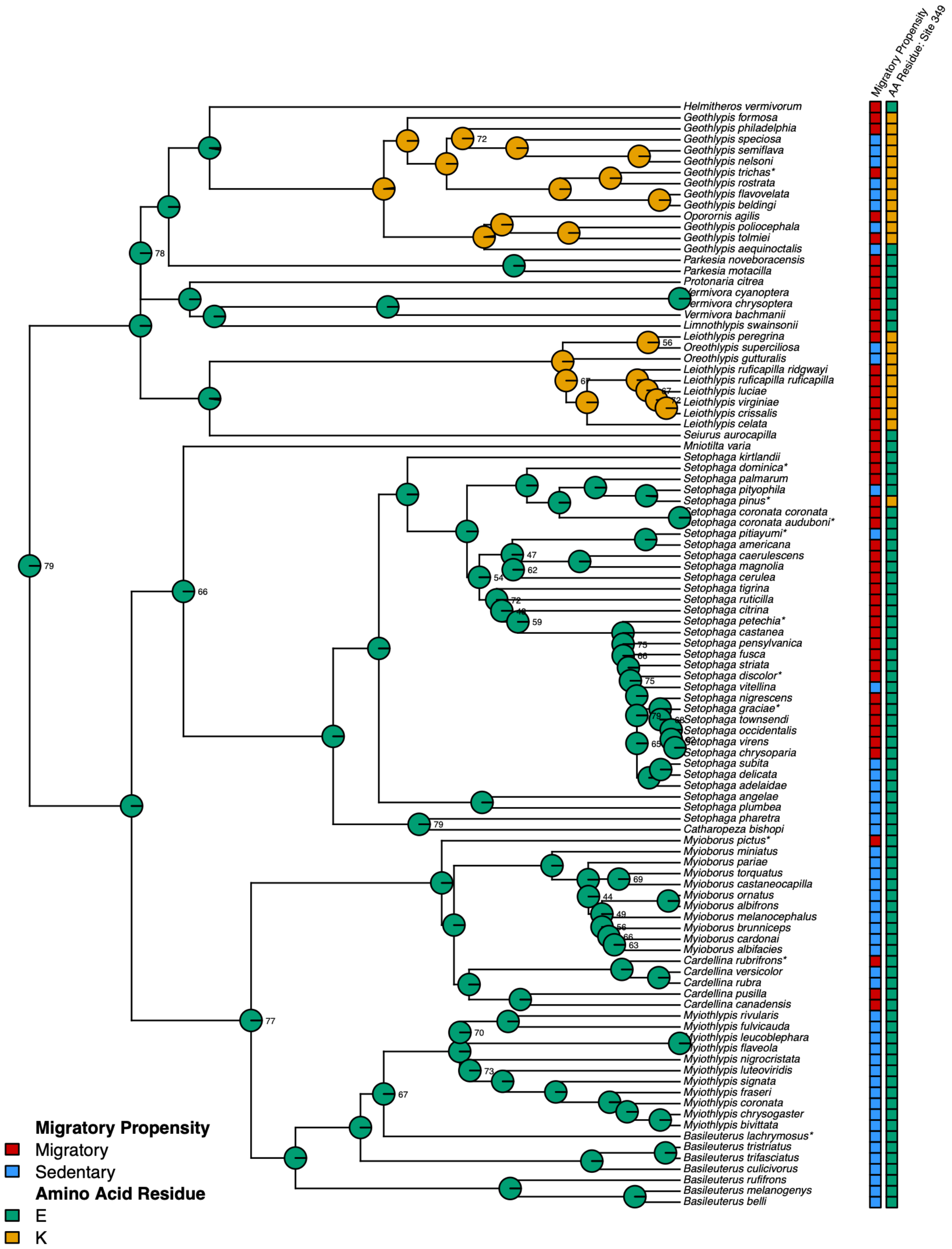

**FIGURE S32. Amino acid variation at *NEK2* site 349 mapped across the *NEK2* gene tree for all parulid warblers.** Square labels at tips represent migratory propensity (red: migratory, blue: sedentary) and amino acid residue (orange: lysine, green: glutamic acid) for each species. Pie charts on nodes show ancestral character state reconstruction posterior probabilities for each amino acid variant. Nodes of the tree that received <80% support by approximate likelihood ratio test (aLRT) are labeled with their aLRT value. Partially migratory species are indicated by asterisks next to their species name.

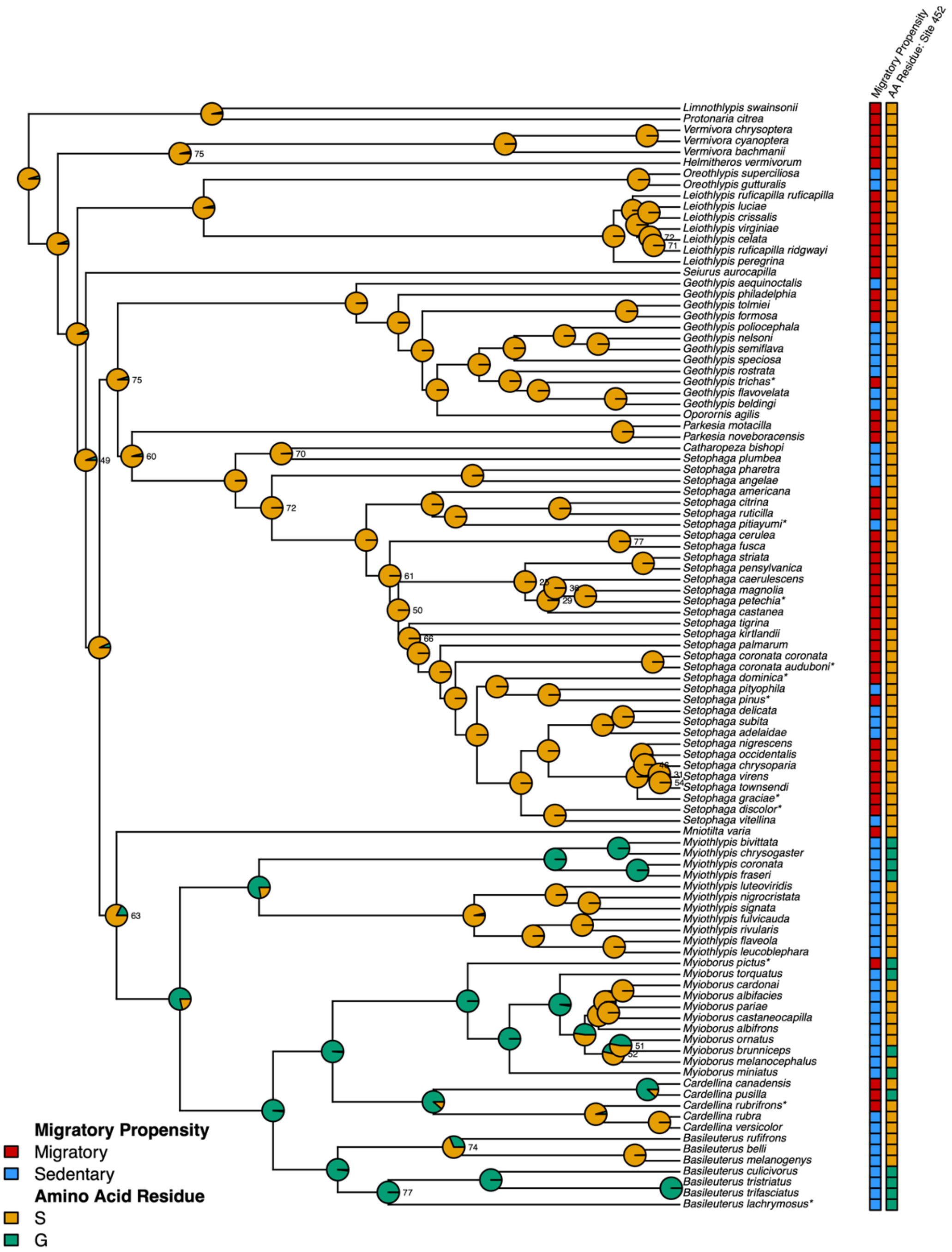

**FIGURE S33. Amino acid variation at *PER3* site 452 mapped across the *PER3* gene tree for all parulid warblers.** Square labels at tips represent migratory propensity (red: migratory, blue: sedentary) and amino acid residue (orange: serine, green: glycine) for each species. Pie charts on nodes show ancestral character state reconstruction posterior probabilities for each amino acid variant. Nodes of the tree that received <80% support by approximate likelihood ratio test (aLRT) are labeled with their aLRT value. Partially migratory species are indicated by asterisks next to their species name.

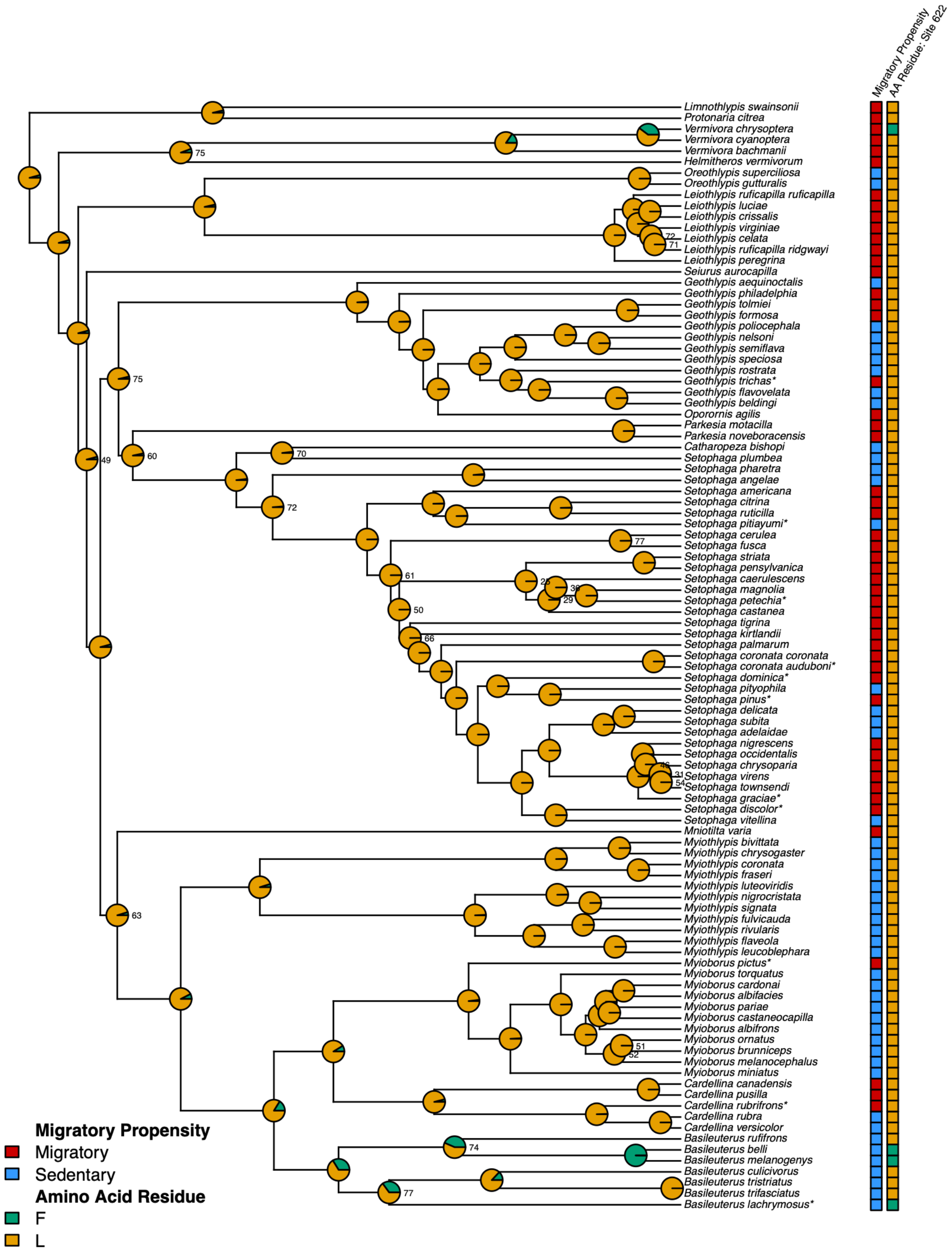

**FIGURE S34. Amino acid variation at *PER3* site 622 mapped across the *PER3* gene tree for all parulid warblers.** Square labels at tips represent migratory propensity (red: migratory, blue: sedentary) and amino acid residue (orange: leucine, green: phenylalanine) for each species. Pie charts on nodes show ancestral character state reconstruction posterior probabilities for each amino acid variant. Nodes of the tree that received <80% support by approximate likelihood ratio test (aLRT) are labeled with their aLRT value. Partially migratory species are indicated by asterisks next to their species name.

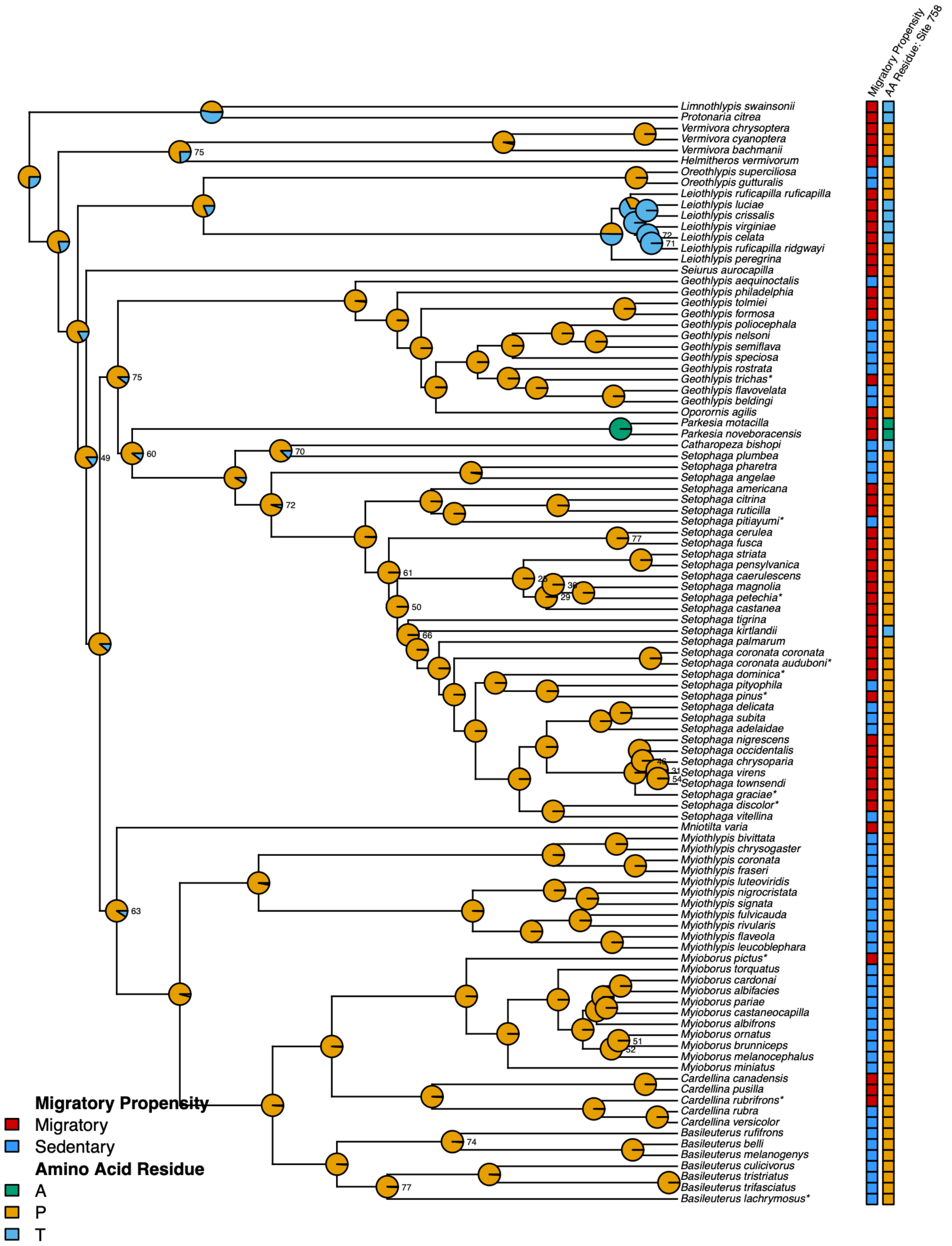

**FIGURE S35. Amino acid variation at *PER3* site 758 mapped across the *PER3* gene tree for all parulid warblers.** Square labels at tips represent migratory propensity (red: migratory, blue: sedentary) and amino acid residue (orange: proline, green: alanine, blue: threonine) for each species. Pie charts on nodes show ancestral character state reconstruction posterior probabilities for each amino acid variant. Nodes of the tree that received <80% support by approximate likelihood ratio test (aLRT) are labeled with their aLRT value. Partially migratory species are indicated by asterisks next to their species name.

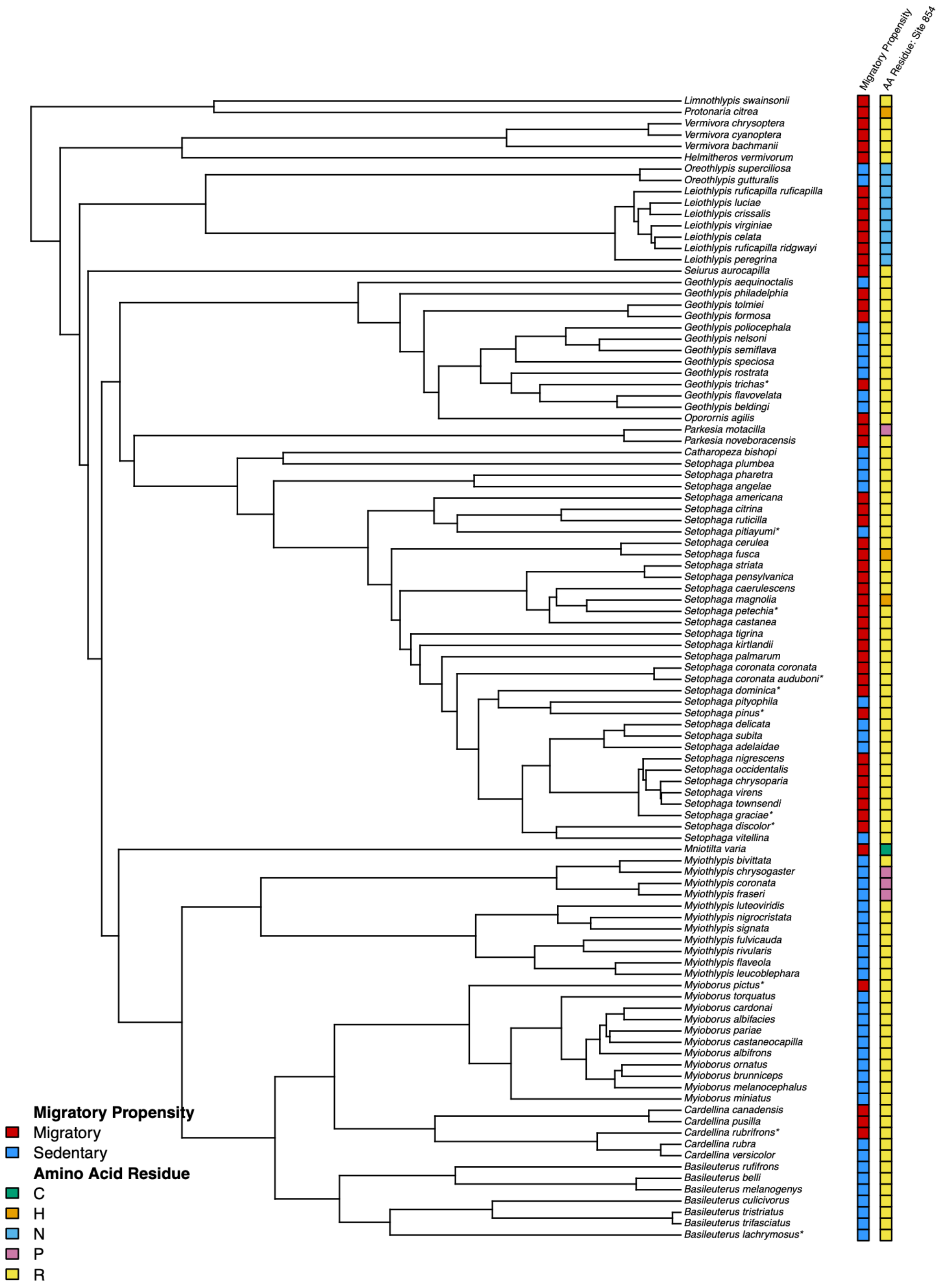

**FIGURE S36. Amino acid variation at *PER3* site 854 mapped across the *PER3* gene tree for all parulid warblers.** Square labels at tips represent migratory propensity (red: migratory, blue: sedentary) and amino acid residue (green: cysteine, orange: histidine, blue: asparagine, pink: proline, yellow: arginine) for each species. Partially migratory species are indicated by asterisks next to their species name.

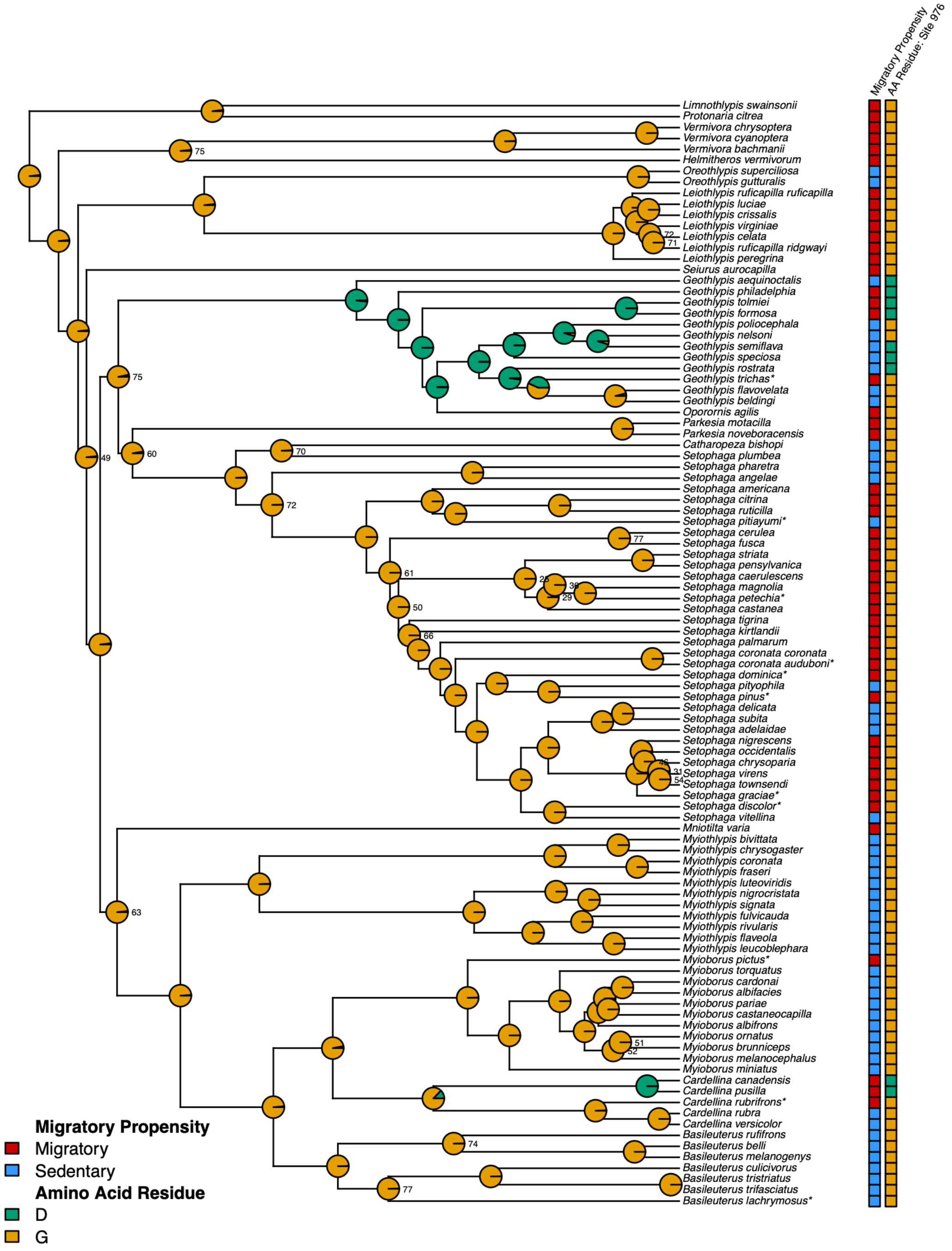

**Figure S37. Amino acid variation at *PER3* site 976 mapped across the *PER3* gene tree for all parulid warblers.** Square labels at tips represent migratory propensity (red: migratory, blue: sedentary) and amino acid residue (orange: glycine, green: aspartic acid) for each species. Pie charts on nodes show ancestral character state reconstruction posterior probabilities for each amino acid variant. Nodes of the tree that received <80% support by approximate likelihood ratio test (aLRT) are labeled with their aLRT value. Partially migratory species are indicated by asterisks next to their species name.

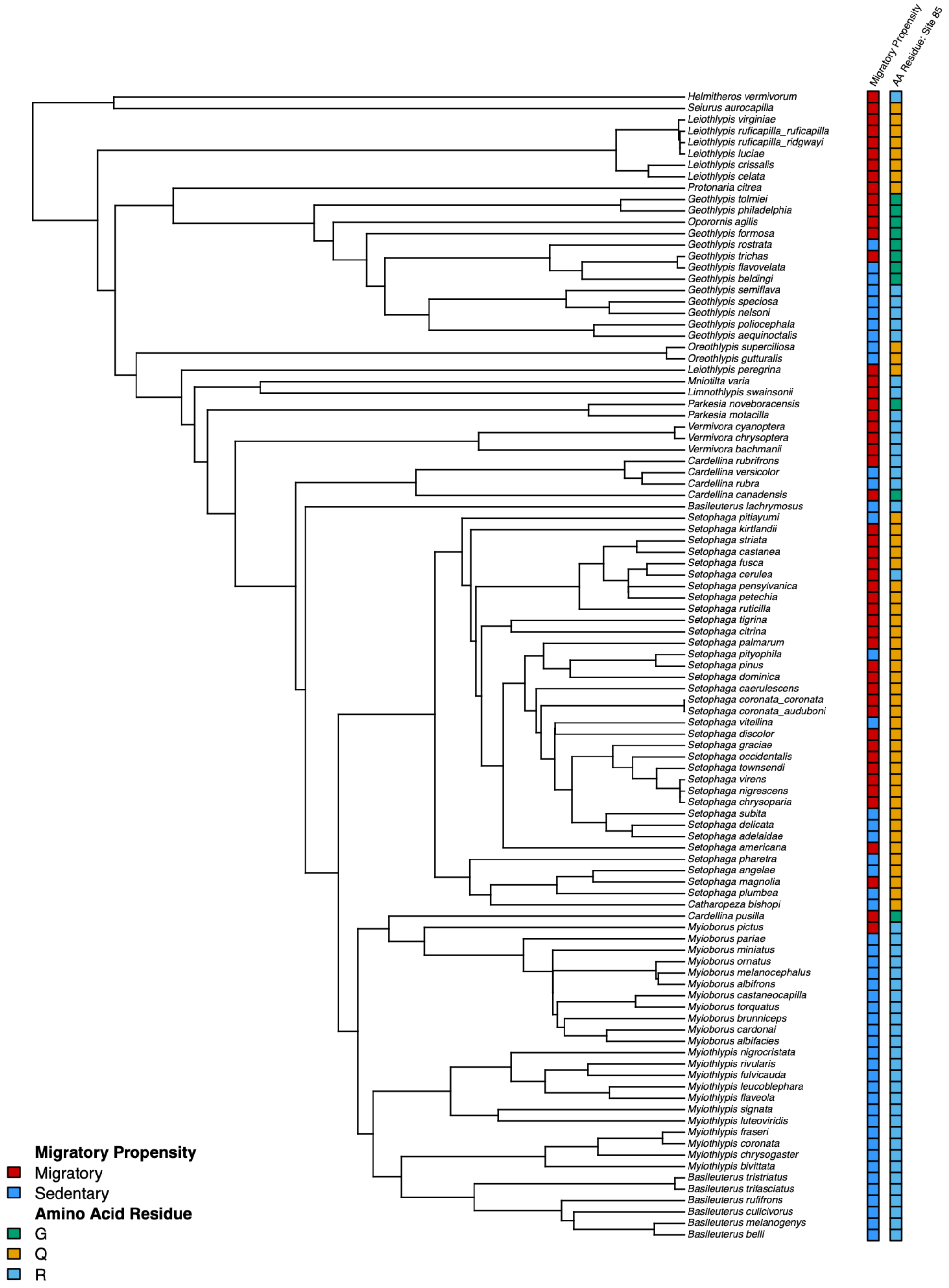

**FIGURE S38. Amino acid variation at *TTR* site 85 mapped across the *TTR* gene tree for all parulid warblers.** Square labels at tips represent migratory propensity (red: migratory, blue: sedentary) and amino acid residue (green: glycine, orange: glutamine, blue: arginine) for each species. Partially migratory species are indicated by asterisks next to their species name.

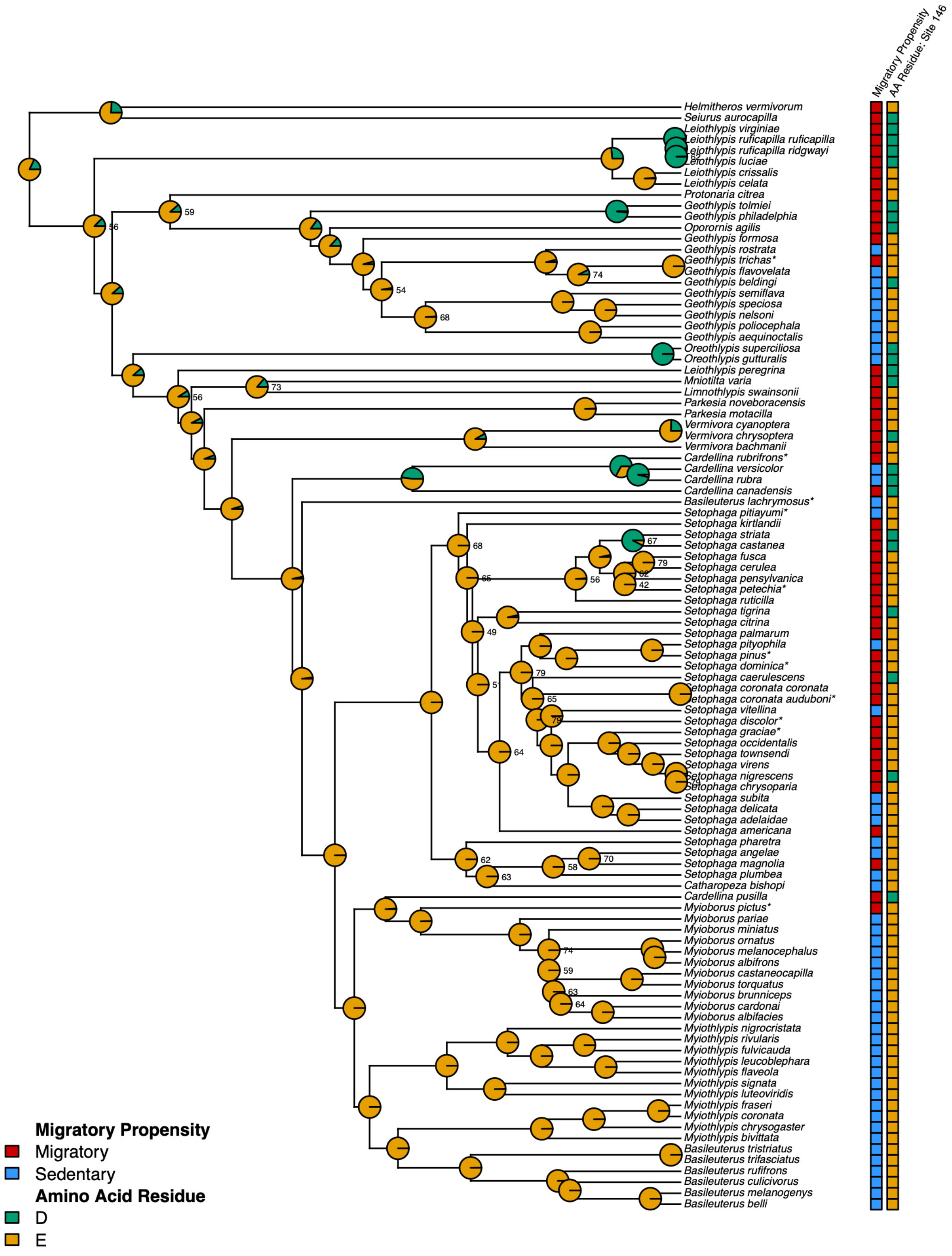

**FIGURE S39. Amino acid variation at *TTR* site 146 mapped across the *TTR* gene tree for all parulid warblers.** Square labels at tips represent migratory propensity (red: migratory, blue: sedentary) and amino acid residue (orange: glutamic acid, green: aspartic acid) for each species. Pie charts on nodes show ancestral character state reconstruction posterior probabilities for each amino acid variant. Nodes of the tree that received <80% support by approximate likelihood ratio test (aLRT) are labeled with their aLRT value. Partially migratory species are indicated by asterisks next to their species name.

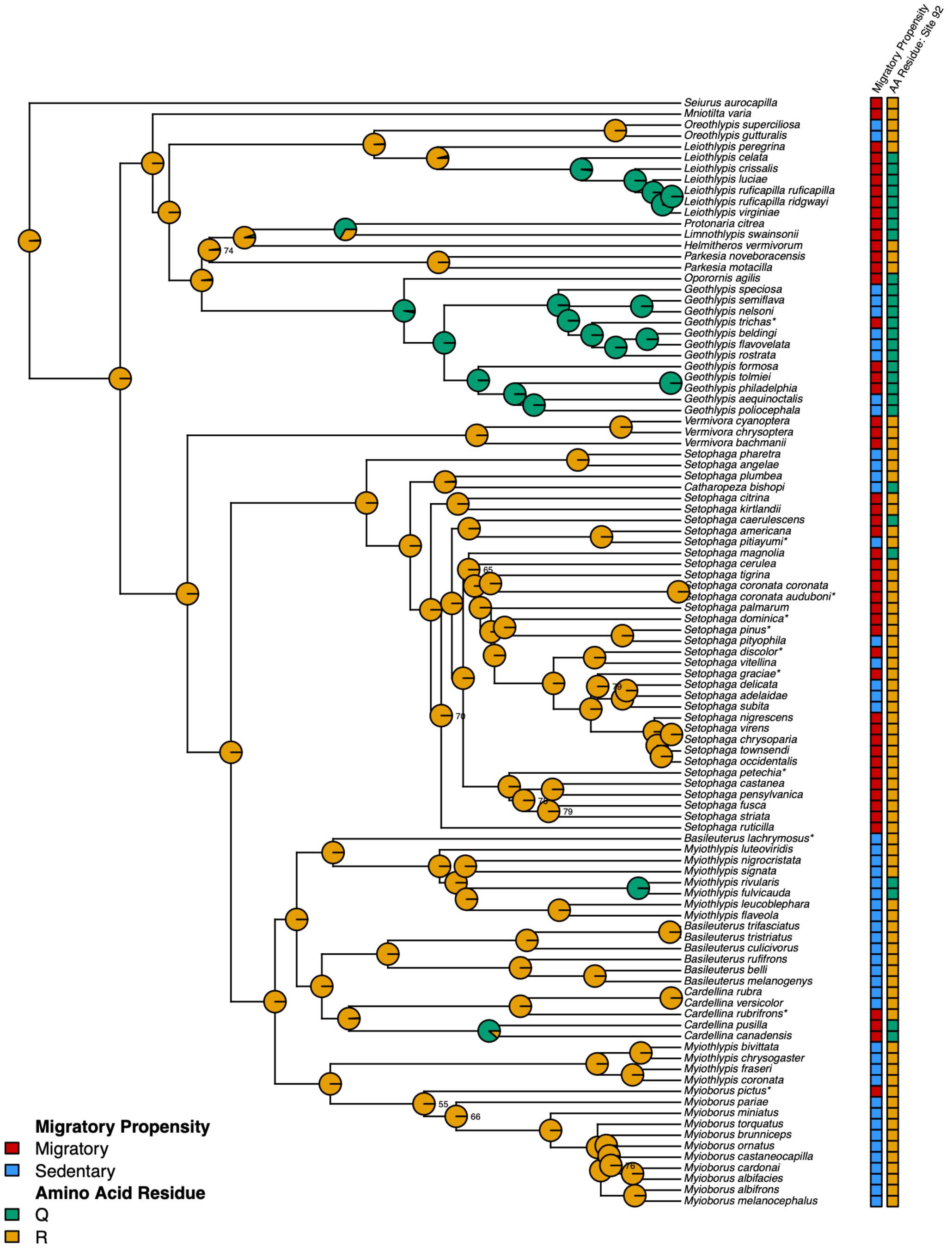

**FIGURE S40. Amino acid variation at *VPS13A* site 92 mapped across the *VPS13A* gene tree for all parulid warblers.** Square labels at tips represent migratory propensity (red: migratory, blue: sedentary) and amino acid residue (orange: arginine, green: glutamine) for each species. Pie charts on nodes show ancestral character state reconstruction posterior probabilities for each amino acid variant. Nodes of the tree that received <80% support by approximate likelihood ratio test (aLRT) are labeled with their aLRT value. Partially migratory species are indicated by asterisks next to their species name.

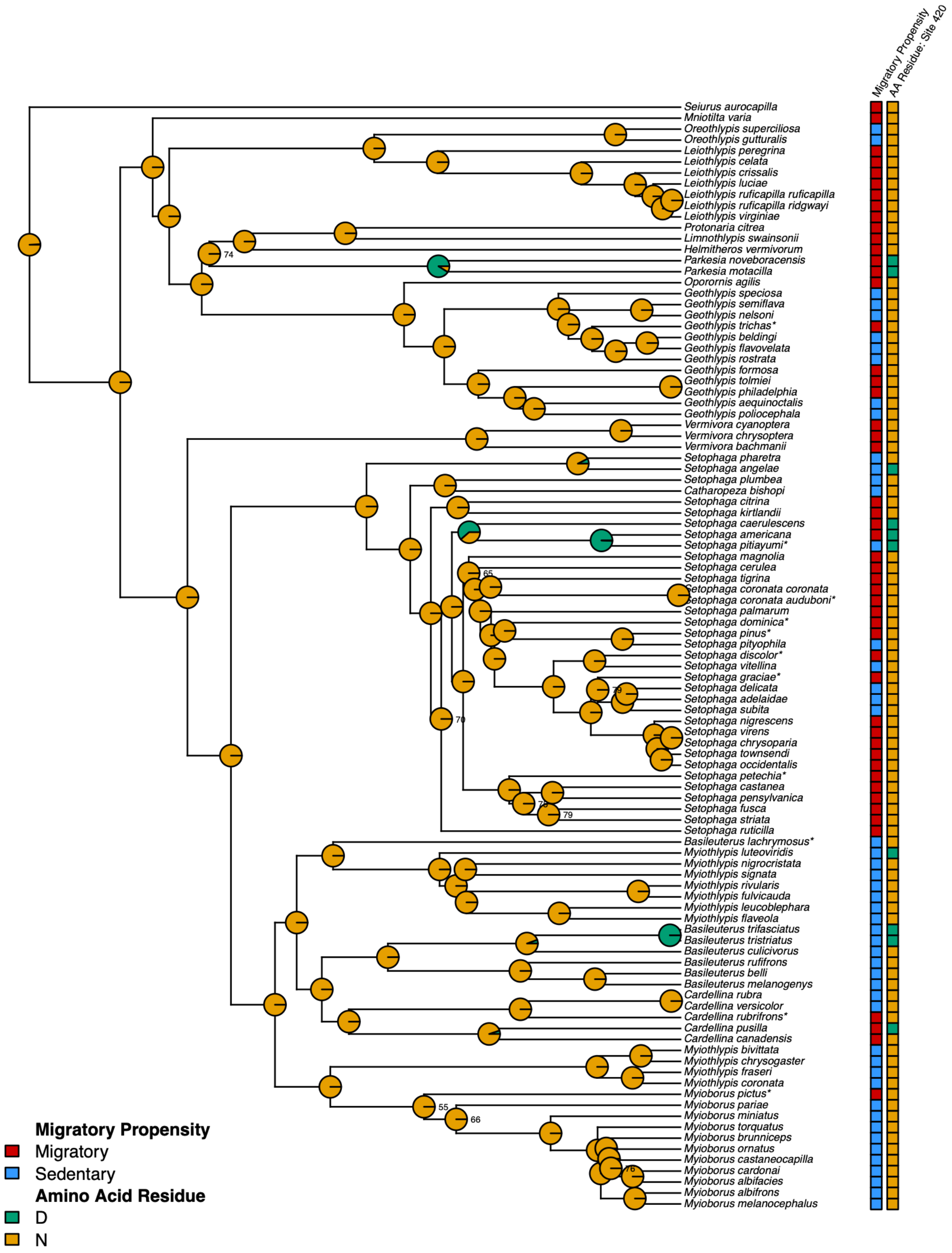

**FIGURE S41. Amino acid variation at *VPS13A* site 420 mapped across the *VPS13A* gene tree for all parulid warblers.** Square labels at tips represent migratory propensity (red: migratory, blue: sedentary) and amino acid residue (orange: asparagine, green: aspartic acid) for each species. Pie charts on nodes show ancestral character state reconstruction posterior probabilities for each amino acid variant. Nodes of the tree that received <80% support by approximate likelihood ratio test (aLRT) are labeled with their aLRT value. Partially migratory species are indicated by asterisks next to their species name.

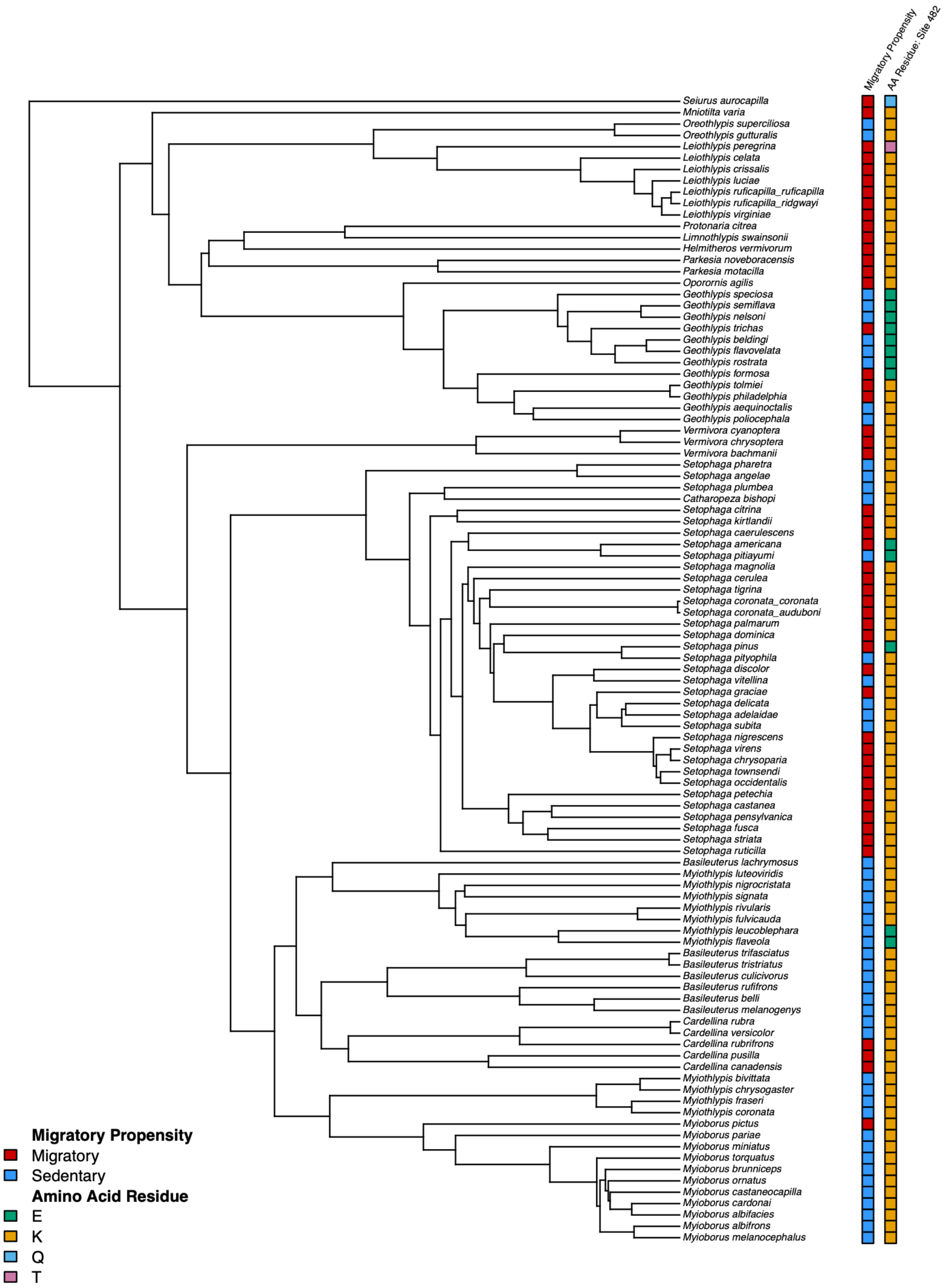

**FIGURE S42. Amino acid variation at *VPS13A* site 482 mapped across the *VPS13A* gene tree for all parulid warblers.** Square labels at tips represent migratory propensity (red: migratory, blue: sedentary) and amino acid residue (green: glutamic acid, orange: lysine, blue: glutamine, pink: threonine) for each species. Partially migratory species are indicated by asterisks next to their species name.

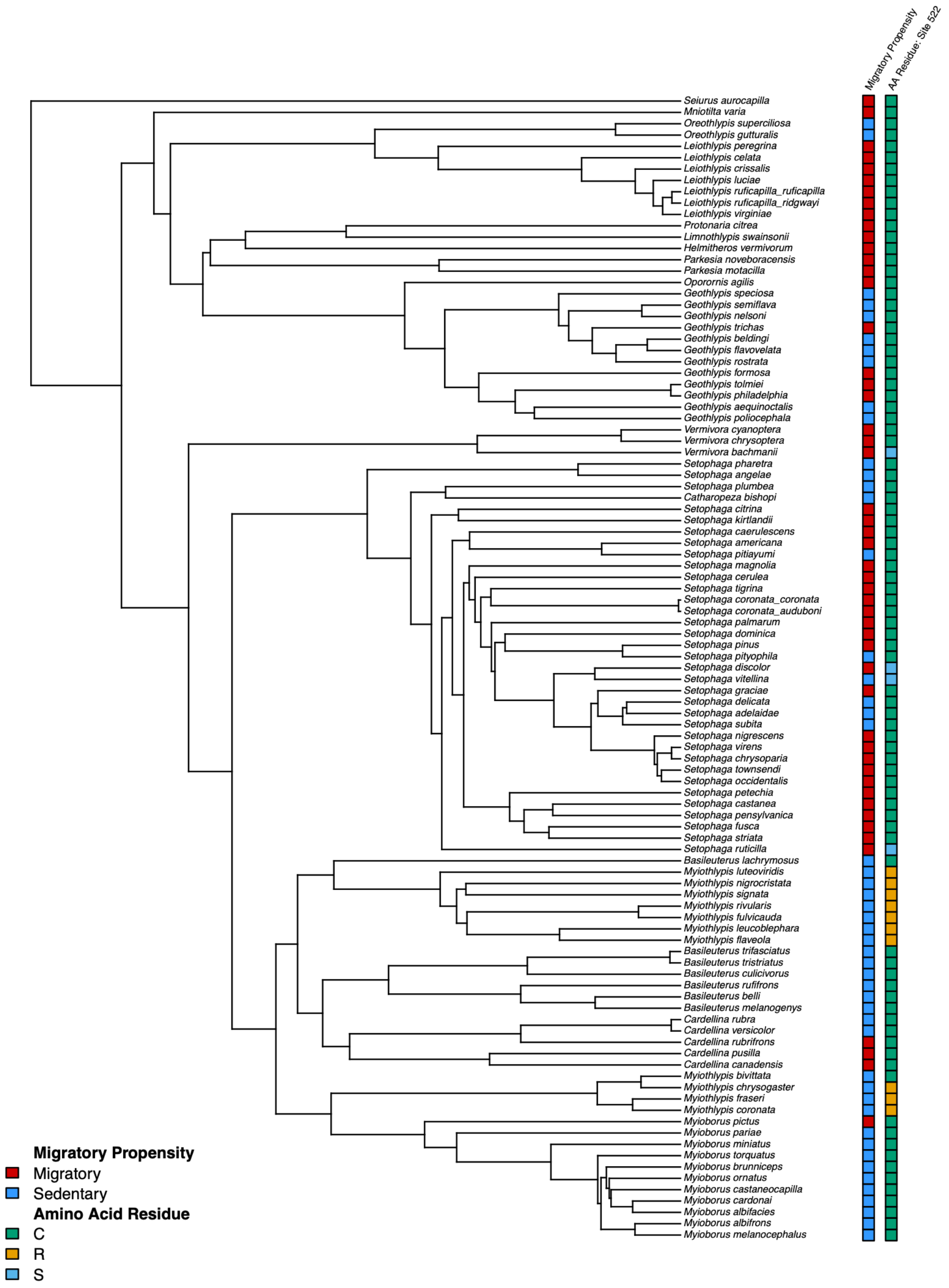

**FIGURE S43. Amino acid variation at *VPS13A* site 522 mapped across the *VPS13A* gene tree for all parulid warblers.** Square labels at tips represent migratory propensity (red: migratory, blue: sedentary) and amino acid residue (green: cysteine, orange: arginine, blue: serine) for each species. Partially migratory species are indicated by asterisks next to their species name.

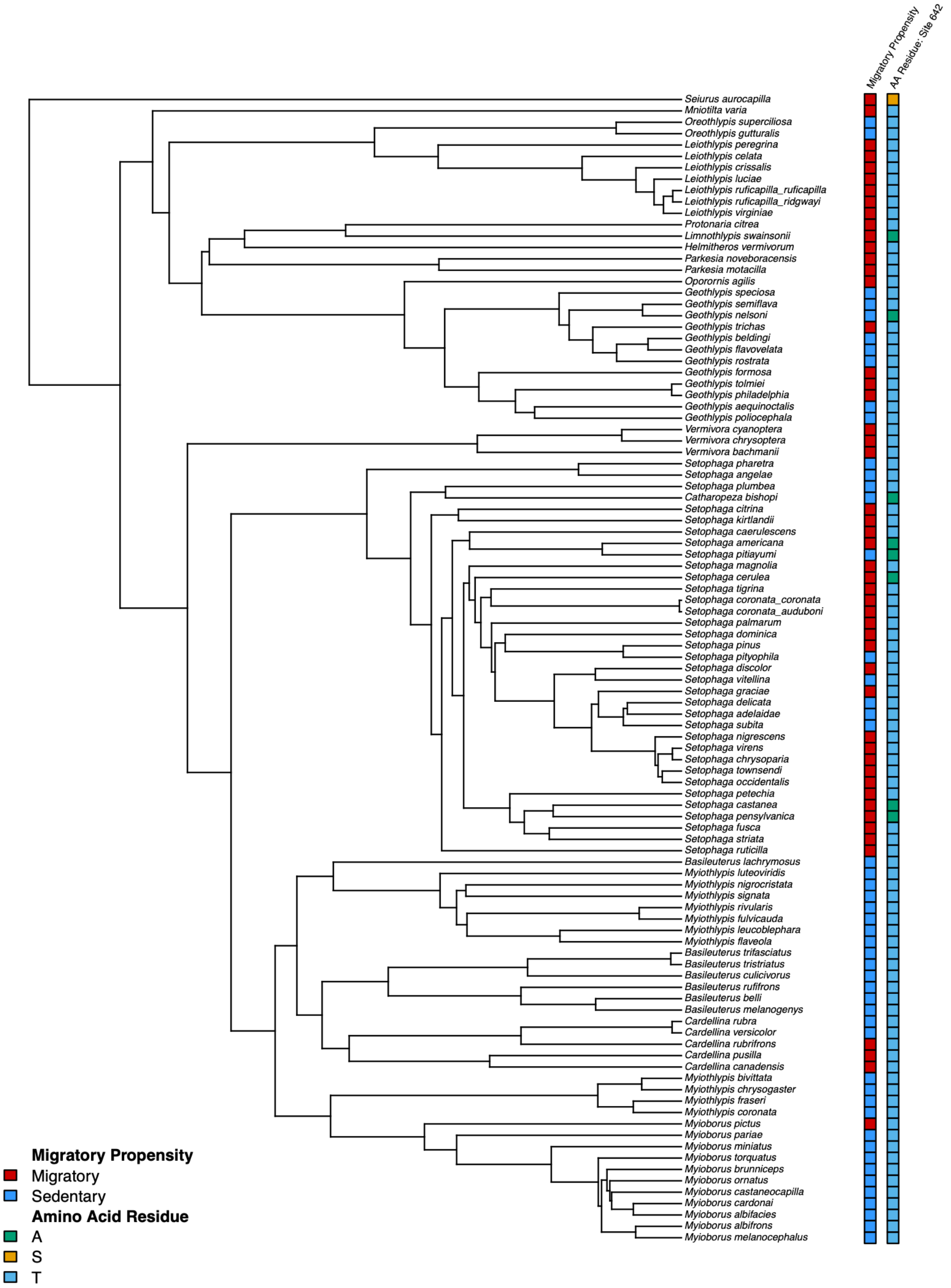

**FIGURE S44. Amino acid variation at *VPS13A* site 642 mapped across the *VPS13A* gene tree for all parulid warblers.** Square labels at tips represent migratory propensity (red: migratory, blue: sedentary) and amino acid residue (green: alanine, orange: serine, blue: threonine) for each species. Partially migratory species are indicated by asterisks next to their species name.

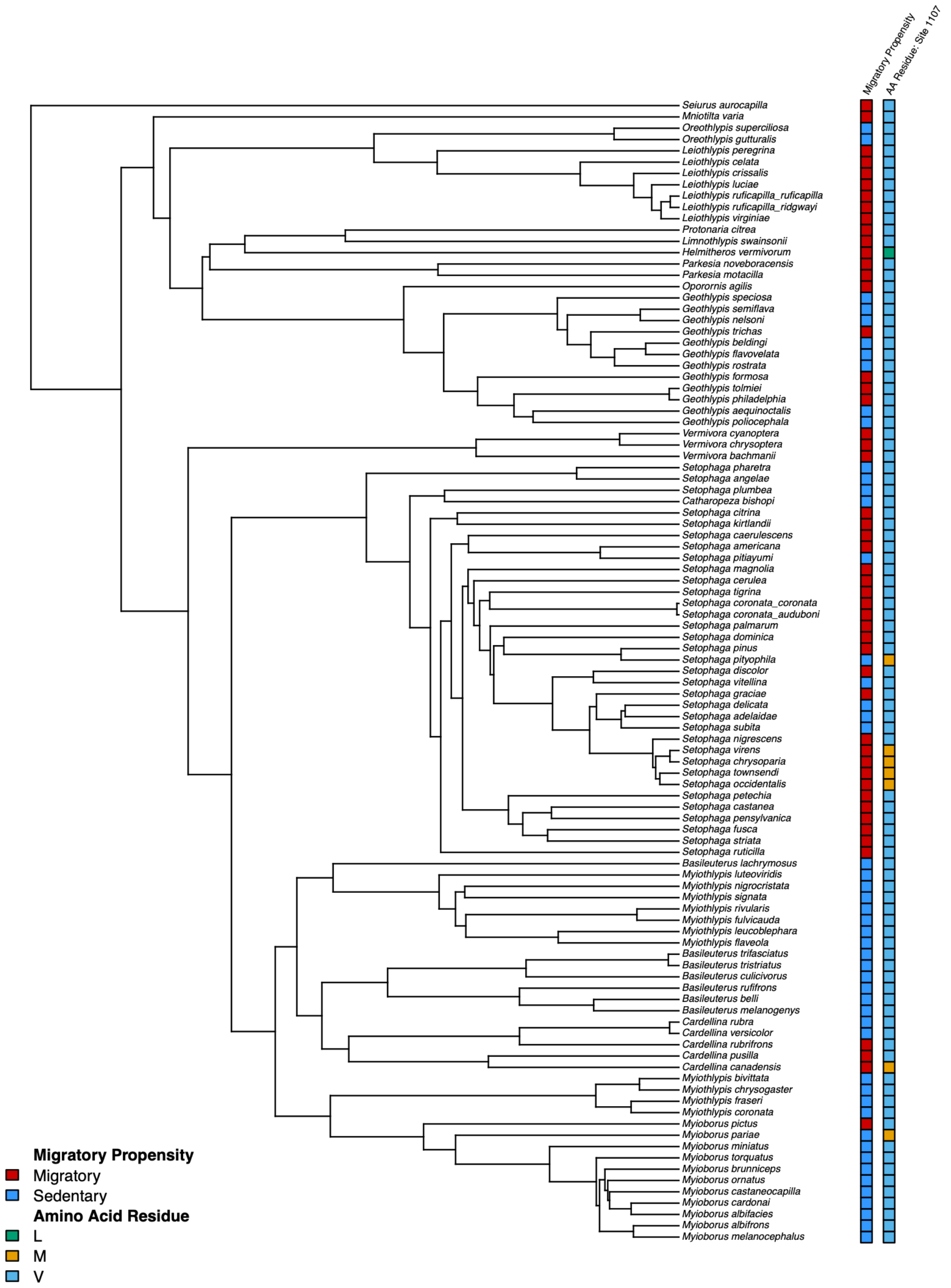

**FIGURE S45. Amino acid variation at *VPS13A* site 1107 mapped across the *VPS13A* gene tree for all parulid warblers.** Square labels at tips represent migratory propensity (red: migratory, blue: sedentary) and amino acid residue (green: leucine, orange: methionine, blue: valine) for each species. Partially migratory species are indicated by asterisks next to their species name.

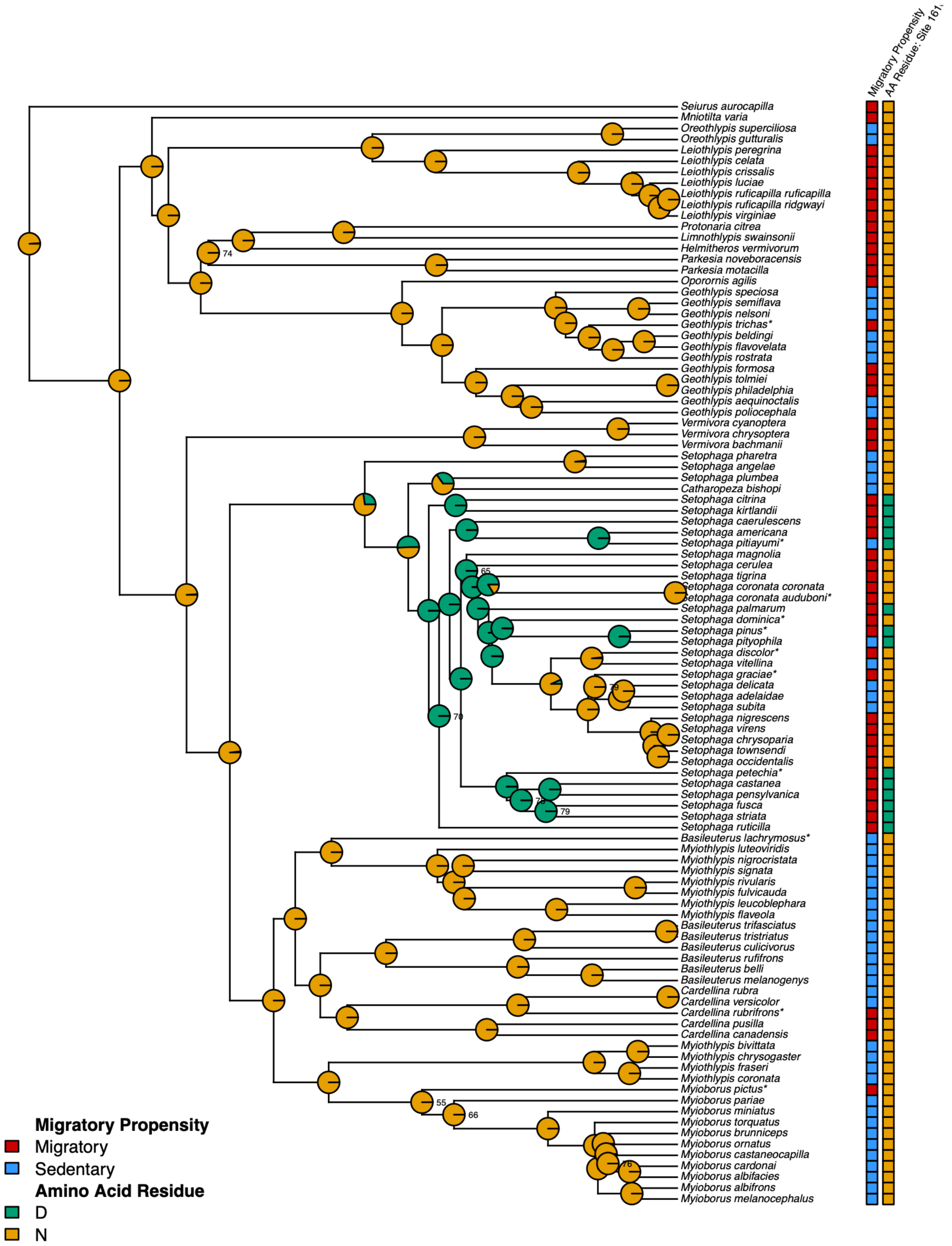

**FIGURE S46. Amino acid variation at *VPS13A* site 1610 mapped across the *VPS13A* gene tree for all parulid warblers.** Square labels at tips represent migratory propensity (red: migratory, blue: sedentary) and amino acid residue (orange: asparagine, green: aspartic acid) for each species. Pie charts on nodes show ancestral character state reconstruction posterior probabilities for each amino acid variant. Nodes of the tree that received <80% support by approximate likelihood ratio test (aLRT) are labeled with their aLRT value. Partially migratory species are indicated by asterisks next to their species name.

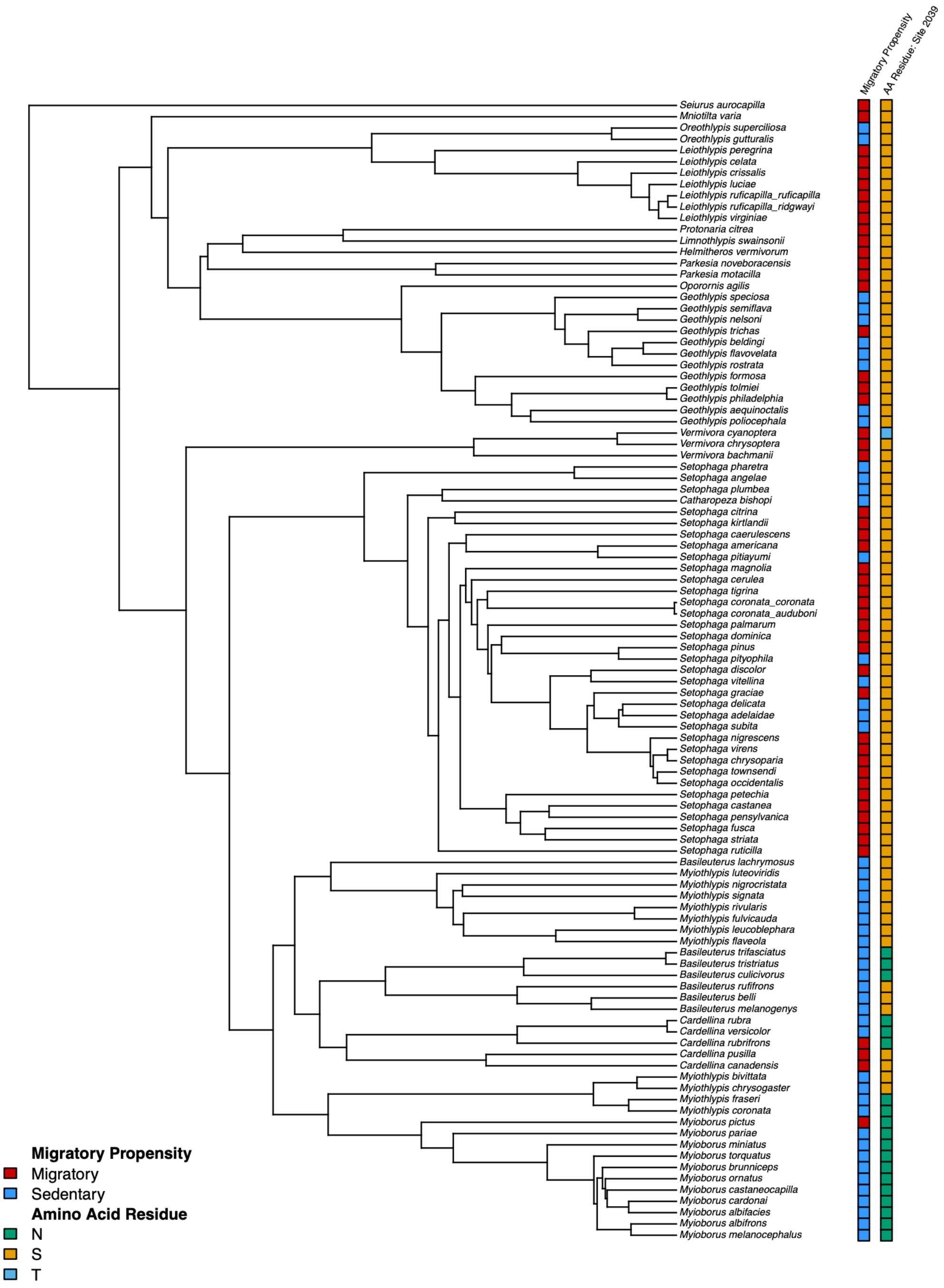

**FIGURE S47. Amino acid variation at *VPS13A* site 2039 mapped across the *VPS13A* gene tree for all parulid warblers.** Square labels at tips represent migratory propensity (red: migratory, blue: sedentary) and amino acid residue (green: asparagine, orange: serine, blue: threonine) for each species. Partially migratory species are indicated by asterisks next to their species name.

### SUPPLEMENTARY TABLES

**TABLE S1. Parulid warbler taxa included in the study, with sample size per taxon and assigned migratory status (S: sedentary, P: partially migratory, M: migratory).** For partially migratory species, whether the taxon was coded as migratory or sedentary for binary analyses is indicated in parentheses.

| Taxon | Sample Size | Migratory Status |
|---|---|---|
| *Basileuterus belli* | 5 | S |
| *Basileuterus culicivorus* | 5 | S |
| *Basileuterus lachrymosus* | 5 | P (S) |
| *Basileuterus melanogenys* | 5 | S |
| *Basileuterus rufifrons* | 5 | S |
| *Basileuterus trifasciatus* | 5 | S |
| *Basileuterus tristriatus* | 5 | S |
| *Cardellina canadensis* | 5 | M |
| *Cardellina pusilla* | 5 | M |
| *Cardellina rubra* | 5 | S |
| *Cardellina rubrifrons* | 5 | P (M) |
| *Cardellina versicolor* | 5 | S |
| *Catharopeza bishopi* | 2 | S |
| *Geothlypis aequinoctialis* | 5 | S |
| *Geothlypis beldingi* | 5 | S |
| *Geothlypis flavovelata* | 5 | S |
| *Geothlypis formosa* | 5 | M |
| *Geothlypis nelsoni* | 5 | S |

| | | |
|---|---|---|
| *Geothlypis philadelphia* | 5 | M |
| *Geothlypis poliocephala* | 5 | S |
| *Geothlypis rostrata* | 4 | S |
| *Geothlypis semiflava* | 5 | S |
| *Geothlypis speciosa* | 5 | S |
| *Geothlypis tolmiei* | 5 | M |
| *Geothlypis trichas* | 5 | P (M) |
| *Helmitheros vermivorum* | 5 | M |
| *Icteria virens* | 6 | M |
| *Leiothlypis celata* | 5 | M |
| *Leiothlypis crissalis* | 5 | M |
| *Leiothlypis luciae* | 4 | M |
| *Leiothlypis peregrina* | 4 | M |
| *Leiothlypis ruficapilla ridgwayi* | 12 | M |
| *Leiothlypis ruficapilla ruficapilla* | 12 | M |
| *Leiothlypis virginiae* | 5 | M |
| *Limnothlypis swainsonii* | 5 | M |
| *Mniotilta varia* | 5 | M |
| *Myioborus albifacies* | 1 | S |
| *Myioborus albifrons* | 3 | S |
| *Myioborus brunniceps* | 4 | S |
| *Myioborus cardonai* | 2 | S |
| *Myioborus castaneocapilla* | 5 | S |

| | | |
|---|---|---|
| *Myioborus melanocephalus* | 5 | S |
| *Myioborus miniatus* | 5 | S |
| *Myioborus ornatus* | 2 | S |
| *Myioborus pariae* | 1 | S |
| *Myioborus pictus* | 5 | P (M) |
| *Myioborus torquatus* | 5 | S |
| *Myiothlypis bivittata* | 4 | S |
| *Myiothlypis chrysogaster* | 2 | S |
| *Myiothlypis coronata* | 5 | S |
| *Myiothlypis flaveola* | 5 | S |
| *Myiothlypis fraseri* | 5 | S |
| *Myiothlypis fulvicauda* | 5 | S |
| *Myiothlypis leucoblephara* | 5 | S |
| *Myiothlypis luteoviridis* | 5 | S |
| *Myiothlypis nigrocristata* | 5 | S |
| *Myiothlypis rivularis* | 4 | S |
| *Myiothlypis signata* | 5 | S |
| *Oporornis agilis* | 5 | M |
| *Oreothlypis gutturalis* | 5 | S |
| *Oreothlypis superciliosa* | 5 | S |
| *Parkesia motacilla* | 5 | M |
| *Parkesia noveboracensis* | 4 | M |
| *Protonotaria citrea* | 5 | M |
| *Seiurus aurocapilla* | 5 | M |

| | | |
|---|---|---|
| *Setophaga adelaidae* | 4 | S |
| *Setophaga americana* | 5 | M |
| *Setophaga angelae* | 5 | S |
| *Setophaga caerulescens* | 5 | M |
| *Setophaga castanea* | 5 | M |
| *Setophaga cerulea* | 5 | M |
| *Setophaga chrysoparia* | 5 | M |
| *Setophaga citrina* | 4 | M |
| *Setophaga coronata auduboni* | 10 | P (M) |
| *Setophaga coronata coronata* | 5 | M |
| *Setophaga delicata* | 2 | S |
| *Setophaga discolor* | 4 | P (M) |
| *Setophaga dominica* | 5 | P (M) |
| *Setophaga fusca* | 4 | M |
| *Setophaga graciae* | 5 | P (M) |
| *Setophaga kirtlandii* | 5 | M |
| *Setophaga magnolia* | 5 | M |
| *Setophaga nigrescens* | 5 | M |
| *Setophaga occidentalis* | 5 | M |
| *Setophaga palmarum* | 4 | M |
| *Setophaga pensylvanica* | 5 | M |
| *Setophaga petechia* | 5 | P (M) |
| *Setophaga pharetra* | 4 | S |
| *Setophaga pinus* | 5 | P (M) |

| | | |
|---|---|---|
| *Setophaga pitiayumi* | 5 | P (S) |
| *Setophaga pityophila* | 4 | S |
| *Setophaga plumbea* | 2 | S |
| *Setophaga ruticilla* | 5 | M |
| *Setophaga striata* | 5 | M |
| *Setophaga subita* | 1 | S |
| *Setophaga tigrina* | 3 | M |
| *Setophaga townsendi* | 5 | M |
| *Setophaga virens* | 5 | M |
| *Setophaga vitellina* | 2 | S |
| *Vermivora bachmani* | 6 | M |
| *Vermivora chrysoptera* | 5 | M |
| *Vermivora cyanoptera* | 5 | M |

**Table S2. List of candidate genes included in this study.** Provided for each are name, location in the myrtle warbler reference genome, broad functional category, and citations for studies that either previously linked the gene with migratory variation or proposed it as a candidate migration gene on the basis of function.

| **Gene** | **Chr** | **Start position** | **Stop position** | **Function** | **Citations linking gene with migration** |
|---|---|---|---|---|---|
| AANAT (SNAT) | 18 | 5864962 | 5869272 | Circadian rhythm | Differentially expressed in *Turdus merula* differing in time to onset of migration (Franchini et al. 2017) |
| ARNTL (BMAL1) | 5 | 29574123 | 29600626 | Circadian rhythm | Proposed as candidate gene by Ruegg et al. (2014) |
| CLOCK | 4 | 27274053 | 27328112 | Circadian rhythm | Associated with migratory propensity, timing, and distance in many taxa—studies summarized in Parody-Merino et al. (2019), Le Clercq et al. (2023b,a) |
| CPNE4 | 2 | 43531933 | 43721676 | Circadian rhythm | Divergent between *Catharus ustulatus* differing in migration route (Ruegg et al. 2014) and associated with migratory timing in *Falco sparverius* (Bossu et al. 2022); differentially expressed in *Zonotrichia leucophrys* in migratory vs. non-migratory states (Jones et al. 2008) |
| CREB1 | 7 | 26104866 | 26125386 | Circadian rhythm | Divergent between *Catharus ustulatus* differing in migration route (Ruegg et al. 2014); associated with migratory propensity in *Geothlypis trichas* (Zamudio-Beltrán et al. 2024) |
| CRY1 | 1a | 51868910 | 51885584 | Circadian rhythm | Divergent between *Catharus ustulatus* differing in migration route (Delmore et al. 2015); differentially expressed in *Sylvia atricapilla* (Fusani et al. 2014) and *Emberiza bruniceps* (Sharma et al. 2018) in migratory vs. non-migratory states |
| CRY2 | 5 | 25404959 | 25429681 | Circadian rhythm | Proposed as candidate gene by Ruegg et al. (2014) |
| CSNK1E | 1a | 49633397 | 49652208 | Circadian rhythm | Divergent between *Catharus ustulatus* differing in migration route (Delmore et al. 2015) |
| DRD4 | 5 | 10745947 | 10747630 | Exploratory behavior | Associated with migratory propensity in *Sialia mexicana* (Sauve et al. 2021) and migration distance in *Melospiza melodia* (Posliff 2020) |
| HRSP12 (RIDA) | 2 | 134240943 | 134249420 | Cellular stress response | Associated with migratory propensity in broad comparison of passerine species (Lugo Ramos et al. 2017) and in *Geothlypis trichas* (Zamudio-Beltrán et al. 2024); Differentially expressed in *Zonotrichia leucophrys* in migratory vs. non-migratory states (Jones et al. 2008) |
| HSP90AA1 | 5 | 50707992 | 50713366 | Cellular stress response | Differentially expressed in *Zonotrichia leucophrys* (Jones et al. 2008) and *Emberiza bruniceps* (Sharma et al. 2021) in migratory vs. non-migratory states |

| | | | | | |
|---|---|---|---|---|---|
| HSP90B1 | 1a | 50801685 | 50811275 | Cellular stress response | Proposed as candidate gene by Ruegg et al. (2014) |
| HSPA5 | 17 | 8091583 | 8094872 | Cellular stress response | Associated with migratory propensity in broad comparison of passerine species (Lugo Ramos et al. 2017); Differentially expressed in *Zonotrichia leucophrys* in migratory vs. non-migratory states (Jones et al. 2008) |
| HSPA8 | Scaffold 45 | 14088 | 20168 | Cellular stress response | Differentially expressed in *Zonotrichia leucophrys* (Jones et al. 2008) and *Emberiza bruniceps* (Sharma et al. 2018) in migratory vs. non-migratory states |
| NEK2 | 3 | 21785715 | 21793413 | Cell cycle regulation | Proposed as candidate gene by Ruegg et al. (2014) |
| NFIL3 | z | 32060876 | 32073550 | Circadian rhythm | Associated with migratory propensity in Geothlypis trichas (Zamudio-Beltrán et al. 2024); differentially expressed in Turdus merula differing in time to onset of migration (Franchini et al. 2017) |
| NPAS2 | 1 | 83910156 | 83964023 | Circadian rhythm | Divergent between *Catharus ustulatus* differing in migration route (Ruegg et al. 2014) and associated with migration timing in male *Phylloscopus trochilus* (Bazzi et al. 2017); differentially expressed in *Emberiza melanocephala* in different migratory states (Trivedi et al. 2014). |
| PARL | 9 | 1407531 | 1419378 | Mitochondrial remodeling | Differentially expressed in *Zonotrichia leucophrys* in migratory vs. non-migratory states (Jones et al. 2008) |
| PER2 | 9 | 2679075 | 2723458 | Circadian rhythm | Divergent between *Catharus ustulatus* differing in migration route (Delmore et al. 2015); associated with migratory propensity in *Geothlypis trichas* (Zamudio-Beltrán et al. 2024); differentially expressed in *Turdus merula* differing in time to onset of migration (Franchini et al. 2017) and in *Emberiza bruniceps* (Sharma et al. 2018) in migratory vs. non-migratory states |
| PER3 | 21 | 3725224 | 3746426 | Circadian rhythm | Associated with migratory propensity in Geothlypis trichas (Zamudio-Beltrán et al. 2024) |
| SLC1A3 | Z | 61751978 | 61815338 | Neurotransmission | Differentially expressed in *Zonotrichia leucophrys* in migratory vs. non-migratory states (Jones et al. 2008) |
| SLC2A1 | 21 | 2282458 | 2293015 | Metabolism | Proposed as candidate gene by Ruegg et al. (2014) |
| TTR | 2 | 107395251 | 107403070 | Hormone transport | Differentially expressed in *Zonotrichia leucophrys* in migratory vs. non-migratory states (Jones et al. 2008) |
| VPS13A | Z | 39230511 | 39317236 | Lipid transport | Associated with differences in migration direction and distance in *Vermivora* warblers (Toews et al. 2019) |
| YPEL1 | 15 | 8030406 | 8048538 | Cell cycle regulation | Differentially expressed in *Zonotrichia leucophrys* in migratory vs. non-migratory states (Jones et al. 2008) |

**Table S3. Results from analyses comparing the topologies of candidate migration gene trees versus the parulid warbler ultra-conserved elements (UCE) species tree.** Reported for each candidate gene are gene name, length of full gene alignment in base pairs, number of parsimony informative sites, and best supported substitution model for the gene tree. We report two related measures of tree discordance for each candidate gene tree compared to the species tree: the Jaccard Robinson-Foulds (JRF) distance, and the normalized JRF distance, which ranges from 0 (trees identical) to 1 (no splits shared between trees). We also report for each gene tree the *D* statistic (a measure of phylogenetic signal; Fritz and Purvis 2010) calculated for migration propensity, the difference between *D* calculated for the gene tree vs. the species tree (more negative values indicate migration propensity has greater phylogenetic signal in the gene tree), and P-values from randomization tests assessing whether the observed gene significantly deviated from the random topology model ($P_{random}$) or the Brownian motion model ($P_{Brownian}$).

| Gene | Alignment Length | Informative Sites | Substitution Model | JRF Distance | Normalized JRF | D | $D_{gene}$ - $D_{species}$ | $P_{random}$ | $P_{Brownian}$ |
|---|---|---|---|---|---|---|---|---|---|
| AANAT (SNAT) | 2676 | 235 | TN+F+R3 | 68.7 | 0.351 | 0.081 | 0.265 | 0.000 | 0.345 |
| ARNTL (BMAL1) | 26252 | 1889 | GTR+F+R4 | 46.1 | 0.235 | -0.214 | -0.030 | 0.000 | 0.821 |
| CLOCK | 56363 | 4395 | GTR+F+R5 | 37.6 | 0.192 | -0.244 | -0.060 | 0.000 | 0.846 |
| CPNE4 | 198275 | 13781 | GTR+F+R5 | 27.6 | 0.141 | -0.381 | -0.197 | 0.000 | 0.937 |
| CREB1 | 21278 | 1489 | TIM3+F+I+R4 | 37.3 | 0.190 | -0.216 | -0.032 | 0.000 | 0.82 |
| CRY1 | 17312 | 1411 | GTR+F+R5 | 37.5 | 0.191 | -0.375 | -0.191 | 0.000 | 0.909 |
| CRY2 | 25238 | 1945 | GTR+F+R5 | 37.7 | 0.192 | -0.207 | -0.023 | 0.000 | 0.831 |
| CSNK1E | 19333 | 1586 | TIM3+F+R3 | 35.0 | 0.179 | -0.277 | -0.093 | 0.000 | 0.858 |
| DRD4 | 1693 | 108 | K2P+R3 | 68.8 | 0.351 | -0.029 | 0.155 | 0.000 | 0.576 |
| HRSP12 (RIDA) | 8773 | 588 | TIM2+F+R3 | 56.2 | 0.287 | -0.072 | 0.112 | 0.000 | 0.625 |

| | | | | | | | | | |
|---|---|---|---|---|---|---|---|---|---|
| HSP90AA1 | 5635 | 397 | GTR+F+R3 | 62.7 | 0.320 | -0.013 | 0.171 | 0.000 | 0.531 |
| HSP90B1 | 9884 | 869 | GTR+F+I+R2 | 36.5 | 0.186 | -0.474 | -0.290 | 0.000 | 0.957 |
| HSPA5 | 3329 | 133 | TN+F+R3 | 56.9 | 0.290 | -0.156 | 0.028 | 0.000 | 0.768 |
| HSPA8 | 4791 | 520 | HKY+F+R4 | 64.7 | 0.330 | -0.285 | -0.101 | 0.000 | 0.850 |
| NEK2 | 7959 | 492 | TPM3u+F+R3 | 59.4 | 0.303 | -0.029 | 0.155 | 0.000 | 0.547 |
| NFIL3 | 13147 | 929 | TIM2+F+I+R3 | 38.4 | 0.196 | -0.320 | -0.136 | 0.000 | 0.886 |
| NPAS2 | 55008 | 3750 | TIM2+F+R4 | 37.2 | 0.190 | -0.159 | 0.025 | 0.000 | 0.757 |
| PARL | 12231 | 993 | GTR+F+I+R3 | 45.1 | 0.230 | -0.141 | 0.043 | 0.000 | 0.739 |
| PER2 | 42573 | 3380 | GTR+F+R4 | 30.2 | 0.154 | -0.270 | -0.086 | 0.000 | 0.868 |
| PER3 | 22050 | 1319 | GTR+F+I+R4 | 44.8 | 0.228 | -0.175 | 0.009 | 0.000 | 0.784 |
| SLC1A3 | 66988 | 6560 | TVM+F+R5 | 28.2 | 0.144 | -0.254 | -0.070 | 0.000 | 0.865 |
| SLC2A1 | 8576 | 659 | TPM2u+F+I+R4 | 52.7 | 0.269 | -0.156 | 0.028 | 0.000 | 0.732 |
| TTR | 8271 | 752 | TVM+F+I+R4 | 56.4 | 0.288 | -0.315 | -0.131 | 0.000 | 0.908 |
| VPS13A | 91622 | 9352 | GTR+F+I+R6 | 31.0 | 0.158 | -0.361 | -0.177 | 0.000 | 0.916 |
| YPEL1 | 18843 | 1097 | GTR+F+R4 | 39.4 | 0.201 | -0.132 | 0.052 | 0.000 | 0.728 |

**Table S4. AIC results comparing models of discrete character evolution for migratory propensity in parulid warblers.** Four models were tested: equal rates (ER), all rates different (ARD), only transitions from migratory to sedentary (M→S) and only transitions from sedentary to migratory (S→M). Shown for each model are log likelihood (logL), degrees of freedom (df), AIC, and AIC weight. The best fitting model is highlighted in bold.

| **Model** | **log(L)** | **df** | **AIC** | **AIC weight** |
|---|---|---|---|---|
| ER | -45.66894 | 1 | 93.33788 | 0.710310 |
| ARD | -45.64163 | 2 | 95.28327 | 0.268543 |
| M→S | -49.55925 | 1 | 101.11849 | 0.014518 |
| S→M | -50.34318 | 1 | 102.68637 | 0.006629 |

**Table S5. Specifications of CODEML models of coding sequence evolution.** The "Model" setting specifies whether ω was fixed across branches (Model = 0) or allowed to vary between foreground and background branches (Model = 2). "NSsites" dictates whether the model assumes one site class for all sites (NSsites = 0), two classes (conserved and neutral; NSsites = 1), or three classes (conserved, neutral and positive; NSsites = 2). We also report which branches were designated as foreground, whether ω was fixed at a single value, and the starting values of ω used; most models were run twice with different starting ω values to ensure convergence.

| Model Name | Model | NSsites | Fixed ω? | Starting ω | Foreground |
|---|---|---|---|---|---|
| m0 | 0 | 0 | No | 0.5, 2 | None |
| migratory branch | 2 | 0 | No | 0.5, 2 | Migratory |
| sedentary branch | 2 | 0 | No | 0.5, 2 | Sedentary |
| migratory branch-site | 2 | 2 | No | 0.5, 2 | Migratory |
| migratory branch-site null | 2 | 2 | Yes | 1 | Migratory |
| sedentary branch-site | 2 | 2 | No | 0.5, 2 | Sedentary |
| sedentary branch-site null | 2 | 2 | Yes | 1 | Sedentary |
| site 1a (nearly neutral) | 0 | 1 | No | 0.5, 2 | None |
| site 2a (positive selection) | 0 | 2 | No | 0.5, 2 | None |

**TABLE S6. Branches of the parulid warbler species tree defined as foreground branches for CODEML analysis of coding sequence evolution.**

| | |
|---|---|
| Migratory foreground branches | Geothlypis trichas |
| | Myioborus pictus |
| | Cardellina rubrifrons |
| | Branch leading to the clade containing *Cardellina canadensis* and *C. pusilla* |
| Sedentary foreground branches | Branch leading to the clade containing *Oreothlypis guttaralis* and *O. superciliosa* |
| | Branch leading to the clade containing Geothlypis beldingi, G. flavovelata, G. trichas, G. rostrata, G. nelsoni, G. semiflava, and G. speciosa |
| | Branch leading to the clade containing Geothlypis aequinoctalis and Geothlypis poliocepha |
| | Setophaga pitiayumi |
| | Setophaga pityophila |
| | Setophaga vitellina |
| | Branch leading to the clade containing *Setophaga adelaidae*, *S. delicata*, and *S. subita* |

**Table S7. Amino acid sites with high posterior probability (Bayes Empirical Bayes P > 0.95) of positive selection (ω > 1) from CODEML site model analyses.** We report results for the five genes for which the site model allowing positive selection (M2a) provided a significantly better fit than the null site model constraining ω ≤ 1 (M1a). For each identified site, we report the position in the amino acid sequence, the posterior probability of ω > 1, the mean estimate of ω and standard error, and the amino acid variants observed at that site across sampled species.

| Gene | Position | Pr(ω>1) | Mean +/- SE for ω | Variants |
|---|---|---|---|---|
| HRSP12 | 93 | 1.000 | 8.382 +/- 1.809 | K / R |
| NEK2 | 73 | 1.000 | 8.029 +/- 1.464 | H / L |
| | 135 | 0.999 | 8.023 +/- 1.477 | D / E |
| | 238 | 0.994 | 7.987 +/- 1.558 | D / E |
| | 253 | 0.994 | 7.985 +/- 1.563 | F / L |
| | 349 | 0.981 | 7.896 +/- 1.743 | E / K |
| PER3 | 452 | 0.999 | 4.366 +/- 0.94 | S / G |
| | 622 | 0.953 | 4.222 +/- 1.16 | L / F |
| | 758 | 1.000 | 4.367 +/- 0.939 | T / A / P |
| | 854 | 1.000 | 4.367 +/- 0.938 | R / C / H / N / P |
| | 976 | 0.956 | 4.214 +/- 1.139 | G / D |
| TTR | 85 | 1.000 | 9.892 +/- 0.843 | R / G / Q |
| | 146 | 1.000 | 9.892 +- 0.843 | E / D |
| VPS13A | 92 | 0.998 | 2.639 +/- 0.359 | Q / R |
| | 420 | 0.994 | 2.633 +/- 0.375 | N / D |
| | 482 | 0.99 | 2.627 +/- 0.388 | K / E / T / Q |
| | 522 | 0.953 | 2.568 +/- 0.49 | C / R / S |
| | 642 | 0.995 | 2.635 +/- 0.37 | A / S / T |

| | | | |
|---|---|---|---|
| 1107 | 0.977 | 2.607 +/- 0.427 | V / L / M |
| 1610 | 0.984 | 2.617 +/- 0.408 | N / D |
| 2039 | 0.961 | 2.581 +/- 0.471 | S / N / T |

Le Clercq L.-S., Bazzi G., Cecere J.G., Gianfranceschi L., Grobler J.P., Kotzé A., Rubolini D., Liedvogel M., Dalton D.L. 2023a. Time trees and clock genes: a systematic review and comparative analysis of contemporary avian migration genetics. Biological Reviews. n/a.

Le Clercq L.-S., Bazzi G., Ferrer Obiol J., Cecere J.G., Gianfranceschi L., Grobler J.P., Kotzé A., Riutort León M., González-Solís J., Rubolini D., Liedvogel M., Dalton D.L. 2023b. Birds of a feather flock together: a dataset for *Clock* and *Adcyap1* genes from migration genetics studies. Sci Data. 10:787.

Lugo Ramos J.S., Delmore K.E., Liedvogel M. 2017. Candidate genes for migration do not distinguish migratory and non-migratory birds. J Comp Physiol A. 203:383–397.

Parody-Merino Á.M., Battley P.F., Conklin J.R., Fidler A.E. 2019. No evidence for an association between Clock gene allelic variation and migration timing in a long-distance migratory shorebird (*Limosa lapponica baueri*). Oecologia. 191:843–859.

Ruegg K., Anderson E.C., Boone J., Pouls J., Smith T.B. 2014. A role for migration-linked genes and genomic islands in divergence of a songbird. Molecular Ecology. 23:4757–4769.

Sauve D., Dale C.A., Tigano A., Ratcliffe L.M., Friesen V.L. 2021. Do candidate genes for migration and behavior explain migratory variation in bluebirds (*Sialia* spp.)? wils. 132:820–829.

Sharma A., Singh D., Gupta P., Bhardwaj S.K., Kaur I., Kumar V. 2021. Molecular changes associated with migratory departure from wintering areas in obligate songbird migrants. Journal of Experimental Biology. 224:jeb242153.

Sharma A., Singh D., Malik S., Gupta N.J., Rani S., Kumar V. 2018. Difference in control between spring and autumn migration in birds: insight from seasonal changes in hypothalamic gene expression in captive buntings. Proceedings of the Royal Society B: Biological Sciences. 285:20181531.

Toews D.P.L., Taylor S.A., Streby H.M., Kramer G.R., Lovette I.J. 2019. Selection on *VPS13A* linked to migration in a songbird. PNAS. 116:18272–18274.

Trivedi A.K., Kumar J., Rani S., Kumar V. 2014. Annual life history–dependent gene expression in the hypothalamus and liver of a migratory songbird: insights into the molecular regulation of seasonal metabolism. J Biol Rhythms. 29:332–345.

Zamudio-Beltrán L.E., Bossu C.M., Bueno-Hernández A.A., Dunn P.O., Sly N.D., Rayne C., Anderson E.C., Hernández-Baños B.E., Ruegg K.C. 2024. Parallel and convergent evolution in genes underlying seasonal migration. Evolution Letters.:qrae064.
